# Inferring Causal Direction Between Two Traits in the Presence of Horizontal Pleiotropy with GWAS Summary Data

**DOI:** 10.1101/2020.09.02.280263

**Authors:** Haoran Xue, Wei Pan

**Affiliations:** School of Statistics, University of Minnesota, Minneapolis, Minnesota 55455; Division of Biostatistics, School of Public Health, University of Minnesota, Minneapolis, Minnesota 55455

**Keywords:** Bi-directional MR, Instrumental variable, GWAS, Mendelian randomization (MR), Steiger’s method, TWAS

## Abstract

Orienting the causal relationship between pairs of traits is a fundamental task in scientific research with significant implications in practice, such as in prioritizing molecular targets and modifiable risk factors for developing therapeutic and interventional strategies for complex diseases. A recent method, called Steiger’s method, using a single SNP as an instrument variable (IV) in the framework of Mendelian randomization (MR), has since been widely applied. We report the following new contributions. First, we propose a single SNP-based alternative, overcoming a severe limitation of Steiger’s method in simply assuming, instead of inferring, the existence of a causal relationship. We also clarify a condition necessary for the validity of the methods in the presence of hidden confounding. Second, to improve statistical power, we propose combining the results from multiple, and possibly correlated, SNPs. as multiple instruments. Third, we develop three goodness-of-fit tests to check modeling assumptions, including those required for valid IVs. Fourth, by relaxing one of the three IV assumptions in MR, we propose methods, including one Egger regression-like approach and its multivariable version (analogous to multivariable MR), to account for horizontal pleiotropy of the SNPs/IVs, which is often unavoidable in practice. All our methods can simultaneously infer both the existence and (if so) the direction of a causal relationship, largely expanding their applicability over that of Steiger’s method. Although we focus on uni-directional causal relationships, we also briefly discuss an extension to bi-directional relationships. Through extensive simulations and an application to infer the causal directions between low density lipoprotein (LDL) cholesterol, or high density lipoprotein (HDL) cholesterol, and coronary artery disease (CAD), we demonstrate the superior performance and advantage of our proposed methods over Steiger’s method and bi-directional MR. In particular, after accounting for horizontal pleiotropy, our method confirmed the well known causal direction from LDL to CAD, while other methods, including bi-directional MR, failed.

**Author Summary:** In spite of its importance, due to technical challenges, orienting causal relationships between pairs of traits has been largely under-studied. Mendelian randomization (MR) Steiger’s method has become increasingly used in the last two years. Here we point out several limitations with MR Steiger’s method and propose alternative approaches. First, MR Steiger’s method is based on using only one single SNP as the instrument variable (IV), for which we propose a correlation ratio-based method, called Causal Direction-Ratio, or simply CD-Ratio. An advantage of CD-Ratio is its inference of both the existence and (if so) the direction of a causal relationship, in contrast to MR Steiger’s prior assumption of the existence and its poor performance if the assumption is violated. Furthermore, CD-Ratio can be extended to combine the results from multiple, possibly correlated, SNPs with improved statistical power. Second, we propose two methods, called CD-Egger and CD-GLS, for multiple and possibly correlated SNPs while allowing horizontal pleiotropy. Third, we propose three goodness-of-fit tests to check modeling assumptions for the three proposed methods. Finally, we introduce multivariable CD-Egger, analogous to multivariable MR, as a more robust approach, and an extension of CD-Ratio to cases with possibly bi-directional causal relationships. Our numerical studies demonstrated superior performance of our proposed methods over MR Steiger and bi-directional MR. Our proposed methods, along with freely available software, are expected to be useful in practice for causal inference.

## 1 Introduction

Mendelian Randomization (MR) has become increasingly popular and widely used to infer causal relationships between pairs of traits, which can be molecular (such as gene expression or DNA methylation), behavioral/educational or clinical [15, 16, 17]. MR (or two-sample MR) utilizes genetic variants, almost exclusively single nucleotide polymorphisms (SNPs), as instruments or instrumental variables (IVs) in two independent GWAS (summary) datasets for the two traits respectively. Although the gold standard for causal inference is randomized clinical trials (RCTs), RCTs are often too costly, unethical and infeasible, in which case MR offers a cost-effective approach. The recent years’ popularity of MR coincides with the ever-increasing availability of large-scale GWAS data and resources/tools like MR-Base [24]. As reviewed in [27], the potential of MR has been fully demonstrated by several notable examples in the studies of cardiovascular and metabolic diseases, such as MR analyses showing of Creactive protein’s and moderate alcohol consumption’s unlikely role in reducing coronary heart disease risk, and replicating RCT findings of several drug targets (e.g. Niemann-Pick C1-like protein). In most MR applications, one often assumes that the causal direction between two traits is known. Otherwise, bi-directional MR can be applied [50, 41]: one first applies MR to infer whether there is a causal relationship from one trait, say *X*, to another trait, say *Y*; then one reverses the roles of the two traits to infer whether there is a causal relationship from *Y* to *X*. If one of the two causal directions does hold, one would hope to detect it with bi-directional MR. Bi-directional MR can be also more intuitively applied graphically [38]: the estimated effect sizes on the two traits from the significant SNPs can be compared in two scatter plots, one from the SNPs significant for each trait; it is expected that, if the causal direction is *X* → *Y*, then there should be a strong relationship between the two sets of the effect sizes on the two traits of the SNPs significant for *X*, but not necessarily so for the SNPs significant for *Y*. However, whether bi-directional MR works depends on one critical, most often unknown, assumption: *two* sets of *valid* SNPs/IVs used in the two directions. That is, if an SNP primarily influences *X*, but influences *Y* only through *X*, then it should be used as an IV to infer *X* → *Y* in the first step; at the same time, it *cannot be used* as an IV to infer *Y* → *X*. In practice, due to limited knowledge, it is almost impossible to always choose valid IVs. Instead, often one just uses all significant (and independent) SNPs for *X* to infer *X* → *Y*, and vice versa for *Y* → *X*. As pointed out by [25], if an SNP primarily influences *X* (and influences *Y* completely through *X*), it will be associated with *both X* and *Y*. Hence, if the sample sizes are large enough, its associations with both *X* and *Y* will be detected, and it will be used as an IV in *both* steps or directions in bi-directional MR, leading to the incorrect conclusion that a causal relationship exists in *both* directions. This problem with bi-directional MR will be confirmed in our real data examples later. Relatedly, the result of bi-directional MR (or its graphical implementation) is confounded by possibly varying statistical power for the two traits (e.g., due to different sample sizes and different types of the traits such as one quantitative and the other binary). In the above example that an SNP primarily influences trait *X* but not *Y*, if the statistical power for association between the SNP and *Y* is large while that for the other pair is small, it is likely that the SNP will be significantly associated with *Y* but not with *X*, thus incorrectly used as an IV for *Y* but not for *X*, likely leading to the false conclusion of a causal effect of *Y* on *X*.

Recently a simple and effective approach, called MR Steiger’s method, was proposed [25]. It was discussed in the context of comparison with the main competitor of MR, mediation analysis, which requires one-sample data (i.e. both SNPs and the two traits are observed in the same sample). The proposed method is based on testing the difference of two (possibly correlated) Pearson’s correlation coefficients using Steiger’s test, and thus named. It was shown that MR Steiger’s method is more robust to measurement errors and hidden confounding than the causal inference test (CIT) [34] and likely other mediation analysis approaches. However, MR Steiger’s method has been most widely used in 2-sample MR applications. In [25], since MR Steiger was discussed in the context of mediation analysis, its derivation was mostly done in the absence of hidden confounding, though in an appendix the issue was briefly discussed under confounding; in contrast, an appealing advantage of MR is its valid causal inference while allowing the presence of hidden confounding, which is largely expected in practice. Therefore, first, we will formulate the problem in the presence of confounding; along the way we will point out for the first time that a key condition is needed for valid inference. Although the condition is expected to hold often in practice, it would be useful to have a method to check the condition; we propose such a method. Second, since MR Steiger’s method is based on using only a single SNP as IV, to improve both its statistical power and applicability, we extend it to allow multiple, and possibly correlated, SNPs. In particular, there is increasing interest in applying transcriptome-wide association studies (TWAS, or PrediXcan) to identify causal genes or other molecular/imaging endophenotypes by integrating GWAS with eQTL/xQTL data [20, 21, 64, 58, 59, 22, 47, 11, 29, 60]. In these applications, multiple correlated cis-SNPs near a gene are used as IVs to impute or predict the gene’s expression level (or another endophenotype) to infer whether the gene’s expression has a causal effect on a trait, say coronary artery disease (CAD); it is assumed that the causal relationship (if existing) is from the gene to the trait. Albeit reasonable, there is possibility of reverse causation: a gene’s expression change is not the cause, but the consequence, of having CAD. Hence, it would be useful to apply a method to confirm whether the causal direction is indeed from the gene to the trait (or vice versa). In such a case, due to often a small sample size of an eQTL study, it would be low powered to apply MR Steiger’s method with a single SNP; instead, it would be more powerful and thus more desirable to apply a method with all the cis-SNPs used to predict the gene’s expression level. Our proposed new methods are more suitable for such an application. The same argument was made for two traits based on large-scale GWAS when single SNP summary statistics-based MR (SMR) [64] was generalized to GSMR with multiple (possibly correlated) SNPs [65]. Third, we develop corresponding goodness-of-fit tests, similar to Cochran’s *Q* statistic, to check the IV and other modeling assumptions for our proposed methods. Fourth, most importantly, with multiple SNPs/IVs, we can relax one of the three key assumptions in MR for valid causal inference: allowing the presence of horizontal pleiotropy, which is expected to be commonplace as shown by previous studies [51, 55]. In particular, the likely violation of the IV assumptions for TWAS and its severe consequences on invalid causal conclusions have been increasingly recognized and discussed in the literature [58, 53, 33, 57]. We propose a valid approach similar to Egger regression, which has been widely used for exactly the same purpose (i.e. in the presence of horizontal pleiotropy) in MR [4]. As a side product of allowing horizontal pleiotropy, each of our methods can be applied to infer whether there is any causal relationship between two traits *and* if so, what is the causal direction; in contrast, MR Steiger’s method can only infer causal direction *after* assuming the existence of one of the two causal directions. Finally, we introduce a multivariable approach, analogous to multivariable MR (MV-MR) [8, 9], as another robust approach; although we focus on uni-directional causal relationships, we also briefly discuss an extension to cases with possibly bi-directional causal relationships.

To demonstrate the wide-ranging utility and superior performance of our proposed methods, we applied both the new and existing methods to infer the causal directions between low density lipoprotein (LDL) cholesterol, or high density lipoprotein (HDL) cholesterol, and CAD. Multiple previous MR analyses have clearly established the causal role of LDL cholesterol on CAD [19, 26, 52, 56], confirmed by meta-analyses of RCTs with statins and other cholesterol-lowering interventions [13, 12, 44]. Hence, the established causal direction of *LDL* → *CAD* can serve as a positive control to test various methods. Indeed, after accounting for horizontal pleiotropy, our methods reached the correct conclusion. In contrast, all other methods, including our method without accounting for horizontal pleiotropy and bi-directional MR (with various variants of MR methods ignoring or robust to or accounting for horizontal pleiotropy), failed to reach a conclusion. On the other hand, contrary to observational studies [18], HDL cholesterol does not appear to have a causal role on CAD, as evidenced by several MR analyses and recent RCTs [26, 52, 56, 2, 43]. Hence, it can serve as a negative control. Again our proposed and most other methods, as expected, could not establish any causal direction between HDL cholesterol and CAD. We also conducted extensive simulation studies to support the wide applicability and validity of our proposed methods.

## 2 Methods

We will first discuss the ideal case when all chosen SNPs are valid IVs, which means each SNP satisfies the three key IV assumptions: (i) each SNP is associated with the exposure; (ii) conditional on the exposure, each SNP is not associated with the outcome; i.e., there is no direct effect of each SNP on the outcome; (iii) each SNP is not associated with hidden confounders. Then we will consider the case when assumption (ii) is violated when one or more SNPs have direct effects, i.e. horizontal pleiotropic effects, on the outcome. We propose an Egger regression-like approach and an alternative based on the Generalized Least Squares (GLS) estimation method, both performing well and similarly. Finally, we introduce a multivariable extension to the Egger regression method, analogous to (MV-MR), as a more robust approach, and an extension to cases with possibly bi-directional causal relationships.

### 2.1 Ideal Case: With Valid IVs (and Hidden Confounding)

#### 2.1.1 True Model

If two traits *X* and *Y* are causally related, we’d like to determine the true causal direction, which can be either *X* → *Y* or *Y* → *X*. In the ideal case where all the *SNPs* used are valid IVs, two possible true models are shown in Figure 1. We use *g* to represent one (but later more SNPs) to be used as IV, and *U* is a hidden confounder (or an ensemble of all confounders). Each directed edge represents a direct and causal effect with the effect size above the line. As usual, we also assume that all the relationships in the models are linear.

**Figure 1:**
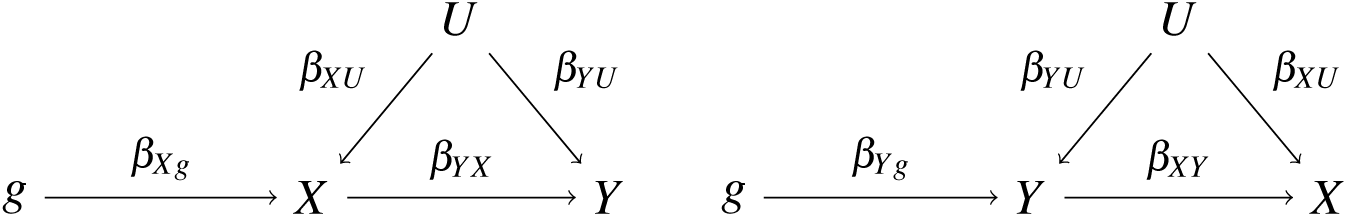
Two possible true models: left one is *X* → *Y*, right one is *Y* → *X*

For *X* → *Y*, the data-generative/causal models are:

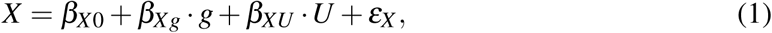

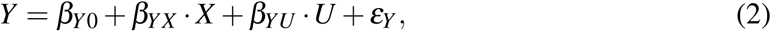

where *ε*_*X*_ and *ε*_*Y*_ are independent error terms.

For *Y* → *X*, we have the corresponding true models.

#### 2.1.2 Steiger’s method based on a single SNP

Based on true models (1) and (2), we have the (true) correlation between *X* and *g*, denoted by *ρ*_*Xg*_, and correlation between *Y* and *g*, denoted by *ρ*_*Yg*_, as

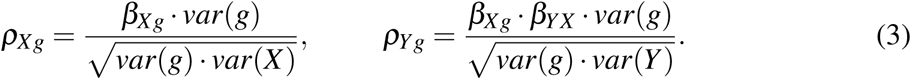

From equation (3) we obtain the correlation ratio:

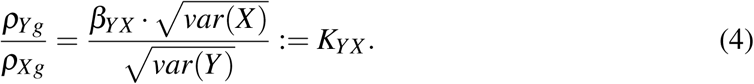

We note that the correlation ratio *K*_*YX*_ is a constant, independent of *g*. It is easy to see that, if there is no confounder *U* in the true models, and without loss of generality assuming *var*(*ε*_*Y*_) *>* 0, we have

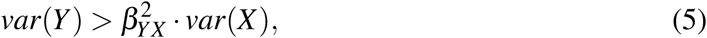

implying |*K*_*YX*_ |< 1. On the other hand, if the true model is *Y* → *X*, we can similarly derive that |*K*_*XY*_| < 1.

It is noted that the two conditions, |*K*_*YX*_| < 1 and |*ρ*_*Yg*_| < |*ρ*_*Xg*_|, are equivalent. While we will use the first one, the second one was used in MR Steiger’s method [25]. In spite of the trivial equivalence between the two, a key insight is that, because *K*_*YX*_ is independent of *g*, it can be used to combine over multiple SNPs/IVs; furthermore, the first one can be used to infer whether there is indeed a causal relationship between the two traits by checking whether *K*_*YX*_ = 0, equivalent to *β*_*YX*_ = 0, in contrast to simply assuming the existence of a causal relationship in Steiger’s method. These two key contributions here are to be elaborated on later.

From two independent GWAS summary datasets with sample sizes *n*_*X*_ and *n*_*Y*_ for traits *X* and *Y* respectively, for SNP *g*, we have its marginal effect estimate on *X* and the variance, 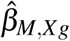 and 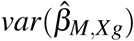, and the corresponding ones for *Y* as 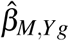 and 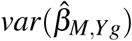. Then we can calculate its sample correlations with *X* and *Y*, denoted by *r*_*Xg*_ and *r*_*Yg*_, from the summary data:

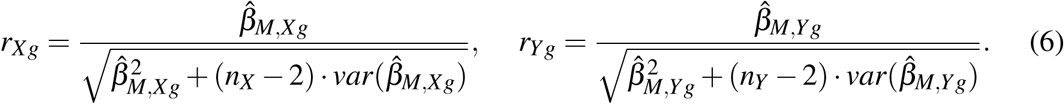

MR Steiger’s method [25] first applies Fisher’s Z-transformation to |*r*_*Xg*_|:

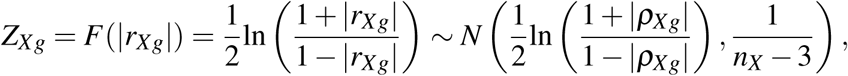

and calculate *Z*_*Yg*_ = *F*(|*r*_*Yg*_|). Then it infers the causal direction by comparing *Z*_*Xg*_ and *Z*_*Yg*_ to obtain *Z*_*g*_ = *Z*_*Xg*_ − *Z*_*Yg*_ and its variance *var*(*Z*_*g*_) = 1*/*(*n*_*X*_ − 3) + 1*/*(*n*_*Y*_ − 3). The test statistic 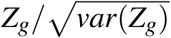 is compared to a standard normal distribution *N*(0, 1) to yield a p-value. If the p-value is larger than a given significance level, MR Steiger’s method would not make any conclusion; otherwise, depending on if *Z*_*g*_ *>* 0 or not, it would conclude that the causal direction is from *X* to *Y*, or from *Y* to *X*.

It is noted that, to test the suitable hypothesis on the magnitudes of the two correlations, MR Steiger’s method takes the Fisher’s transformation on the absolute values of the two correlations, i.e., |*r*_*Xg*_| and |*r*_*Yg*_|. For finite samples, using the non-negative |*r*_*Xg*_| and |*r*_*Yg*_| may lead to poor approximations by their normal distributions, though it is asymptotically valid (under usual regularity conditions, including each correlation is not 0 or *±*1). Figure S29 in the Supplementary gives an illustration.

It is clear that the key condition for Steiger’s method as proposed by [25] is inequality (5). It is expected to hold with the following intuitive explanation: based on true model (2), *var*(*Y*) represents the total variation of *Y* while 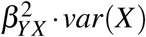 is only its partial variation mediated through *X*. However, in the presence of confounder *U*, as shown in equation (2), we have

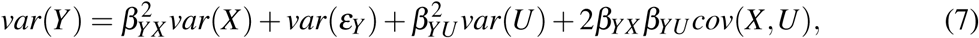

from which we cannot conclude that (5) holds, simply because the sum of the last three terms in *var*(*Y*) can be negative. A sufficient condition for (5) is that *β*_*YX*_ *β*_*YU*_ *cov*(*X,U*) is (i) positive, or (ii) negative but small in magnitude. Assumption (i) means that the directions of the correlations between *X* and *U* and between *Y* and *U* are consistent with the sign of *β*_*YX*_ : for example, if *β*_*YX*_ *>* 0, then the effects of *U* on *X* and *Y* are in the same direction; on the other hand, if *β*_*YX*_ < 0, then the effects of *U* on *X* and *Y* are in different directions. A sufficient condition for (ii) is the small confounding and small causal effect size. For complex traits, we believe that these assumptions (i) and (ii) are often reasonable, albeit untestable directly due to that the confounder *U* is unobserved. Furthermore, [25] showed numerically that, if the proportion of the variance explained between the two traits, 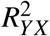, is less than 0.2, then the condition almost always holds. It is noted that for complex human traits *X* and *Y* that are influenced by many genetic *and* environmental factors, 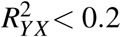 is expected to hold often.

Nevertheless, as an assumption, it would be desirable to check condition (5) in practice. Next we propose such a method. First, we can obtain an estimate 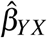 of the causal effect *β*_*YX*_, e.g. by MR Egger regression. Second, if we have individual level data, then we can estimate *var*(*X*) and *var*(*Y*) directly, thus verify whether the condition holds. It is noted that *var*(*X*) and *var*(*Y*) are just the marginal variances of the two traits, which can be consistently estimated by the corresponding sample variances. On the other hand, if we only have GWAS summary data for both traits, we propose the following procedure to estimate *var*(*X*) and *var*(*Y*). For each SNP *g*, we have its marginal association estimate 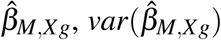 and *var*(*g*) = 2MAF(1 − MAF) in the marginal model 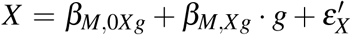, where MAF is the minor allele frequency of the SNP. Then it is easy to verify that

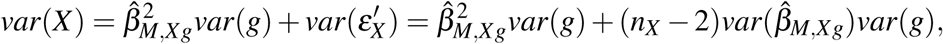

where *n*_*X*_ is the sample size. We can similarly estimate *var*(*Y*). Since such variance estimates may depend on the SNP *g* being used, we can obtain many such estimates based on each of many SNPs, then check the distribution of *var*(*Y*)*/var*(*X*) and compare to 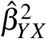 to verify condition (5).

In practice, as to be shown in our real data examples, in order to apply MR Steiger’s method and its extensions to be proposed next, we first assume that, if the causal direction is from *X* to *Y*, then condition (5) holds, then we apply our above proposed procedure to check it.

#### 2.1.3 New Method with Multiple and Possibly Correlated SNPs

Steiger’s method is based on using only one SNP as IV. There is a good reason to extend it to multiple and possibly correlated SNPs, e.g. from a genomic locus: to improve statistical efficiency with multiple S NPs. In particular, in some applications such as TWAS [20, 21], it is necessary to use multiple correlated cis SNPs to more effectively predict/impute one gene’s expression level; if we are interested in inferring whether the causal direction is from the gene to a GWAS trait, we have to use multiple correlated SNPs.

Now we describe the new method for multiple, and possibly correlated SNPs. Due to linkage disequilibrium (LD), it is likely that some SNPs at the same locus are correlated and are significantly associated with both *X* and *Y*. Suppose we have *m* such SNPs denoted in *g*; for each SNP *i*, from equation (6), with summary statistics we can get its sample correlations with *X* and *Y* respectively as 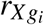 and 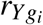. Denote 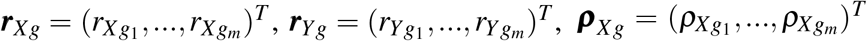 and 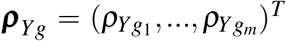. As discussed in [36], we obtain the asymptotic distributions of 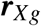 and 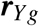 as

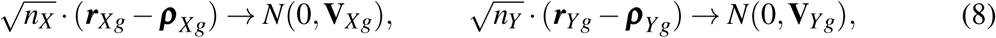

where the covariance matrices 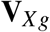 and 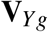depend on the correlation matrix of the SNPs in a complicated way, as shown in the Supplementary. With GWAS summary data, we use a reference panel, such as one from the 1000 Genomes Project [49], to estimate the correlation matrix of the SNPs. Given the asymptotic distributions of 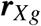 and 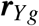, and the two independent GWAS datasets, 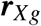 and 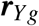 are also independent, leading to a block-diagonal covariance matrix for their joint distribution. By the delta method, we can obtain:

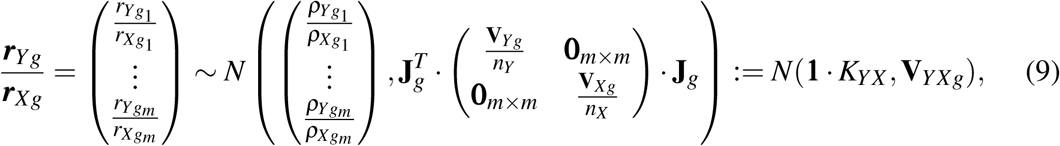

where “:=“represents “by definition”, **1** = (1,…, 1)^*T*^ and

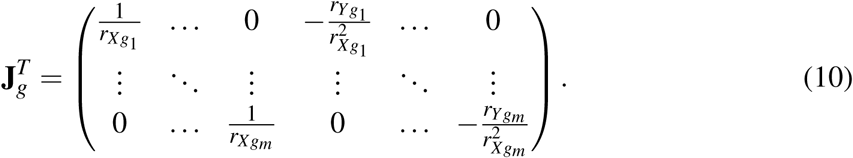

As we have already shown, 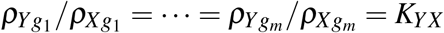 is a constant, independent of the SNPs. Hence, the GLS method is applied to estimate *K*_*YX*_ as

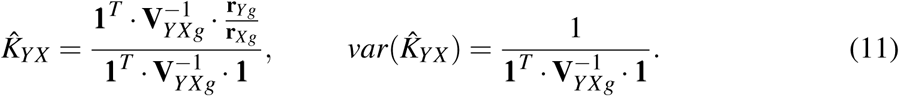

We call this method **Causal Direction-Ratio**, or **CD-Ratio** for short. It is noted that **CD-Ratio** can be applied to special situations with only one SNP, or with multiple independent SNPs. Note that for two independent SNPs, their sample correlations with the same trait are in general non-independent.

With each individual estimate, or the combined estimate 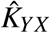 and and its 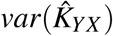, we can construct a confidence interval for *K*_*YX*_. Similarly, from the other direction we can estimate *K*_*XY*_ as 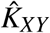 with 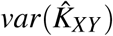. Assuming condition (5) is satisfied, we infer the causal direction between *X* and *Y* by comparing whether the confidence interval of *K*_*XY*_ (or *K*_*YX*_) is completely within or outside interval [-1, 1], as to be discussed later.

### 2.2 Extension to Case with Invalid IVs with Direct Effects

Now consider one or more SNPs/IVs with direct effects on the outcome as shown in Figure 2. The true causal models are

**Figure 2:**
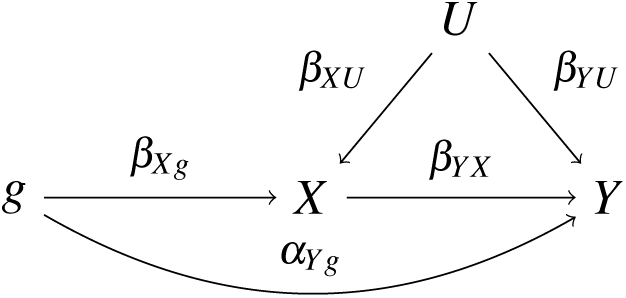
True model for causal direction from *X* to *Y* in the presence of pleiotropic/direct effect (i.e. 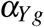).

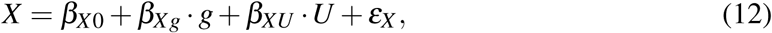

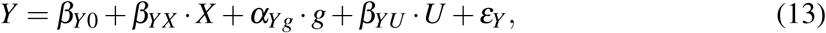

Implying

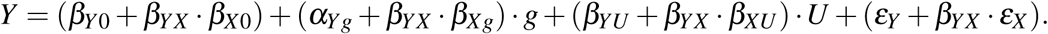

Accordingly, we calculate the (Pearson’s) correlations between the SNP and *X* and *Y*,

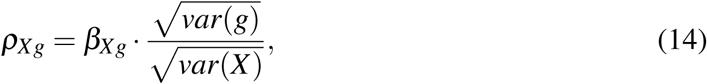

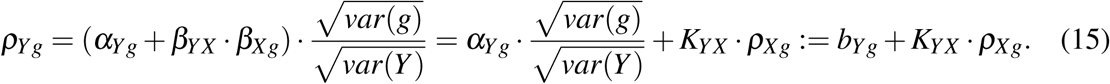

Again by definition 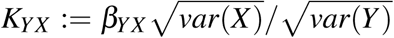; assuming condition (5) is satisfied, we have |*K*_*YX*_| < 1. Furthermore, *K*_*YX*_ = 0 is the sufficient and necessary condition for *β*_*YX*_ = 0, i.e. no causal relationship from *X* to *Y*. Different from that in the Ideal Model, as pleiotropic effects are allowed now, even there is no causal relationship between *X* and *Y*, there could still be significant SNPs *g* for both traits due to pleiotropy. By testing whether *K*_*YX*_ = 0 or not, we can determine the existence of a causal relationship.

It is noted that 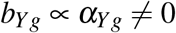. We next develop an estimation method by treating 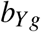’s as random effects, similar to that in MR Egger regression [4]. For simplicity we assume *g*’s are standardized to have mean 0 and variance 1, and the covariance matrix of *g*’s is Σ. Since *g*’s are possibly correlated, we may have correlated 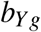’s. Assuming the direct effects from *g* to *Y* are independently from a normal distribution, we would have

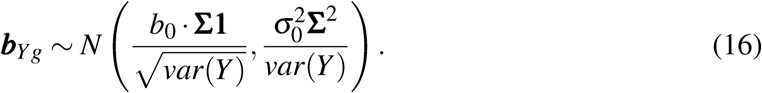

See Supplementary for the<u>deriva</u>tion. For simplicity, in the following we denote ***v*** = Σ**1** = (*v*_1_, …, *v*_*m*_)^*T*^, *b*_0_ for 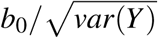, and 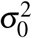 for 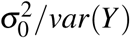. Using the sample correlation coefficients to approximate the population correlation coefficients in equation (16) (while ignoring the errors in 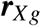), we have

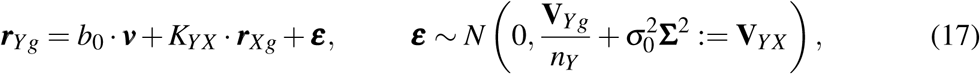

in which the InSIDE assumption (i.e. the independence between 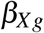 and 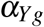) is imposed (as for MR Egger regression). Given 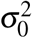, estimating (*b*_0_, *K*_*YX*_) is a GLS problem, thus we estimate (*b*_0_, *K*_*YX*_) and 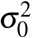 iteratively, and obtain their standard errors 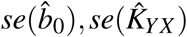; see Supplementary for details. Briefly, we simply regress the two sample correlations with an intercept term to estimate the slope parameter *K*_*YX*_ using the GLS method (which reduces to the weighted least squares with the weight inversely proportional to 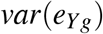) if the SNPs are independent). This is similar to Egger regression in MR, for which the effect size estimates, instead of the sample correlation coefficients, are regressed against each other. We call this method **Causal Direction-Egger**, or **CD-Egger** for short.

A possible downside of the regression approach is its ignoring the variability of 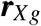 while account for that of 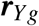, similar to that in MR Egger regression [61]. To overcome this short-coming, we propose a new method called **Causal Direction-GLS**, or simply **CD-GLS**. Since it performs similarly to CD-Egger (due to small variability of 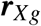 with a usually large sample size of GWAS), we relegate its detail to the Supplementary.

### 2.3 Remarks on Extension to Case with Invalid IVs Associated with Confounders

Now we consider an SNP with a direct effect on the hidden confounder, as shown in the DAG in the left panel of Figure 3. This is the case when the third key IV assumption is violated. Note that, as in [4], for simplicity we assume an effect size 1 of the confounder *U* on both *X* and *Y*, while it can be easily generalized. The case can be converted to that as if the second key IV assumption is violated, as shown in the right panel of Figure 3. It may appear that the two new methods developed in the previous subsection can be applied. However, the InSIDE assumption (i.e. the independence assumption of the IV effect on the exposure *X, γ* = *γ*_1_ + *β*, and that on the outcome *Y, α* = *α*_1_ + *β*) is clearly violated because of their shared component *β*. Hence, applying the two new methods developed in the previous section will lead biased inference, unless the direct effect *β* is small and thus negligible. We will not pursue this challenging topic further, which is beyond the scope of this paper.

**Figure 3:**
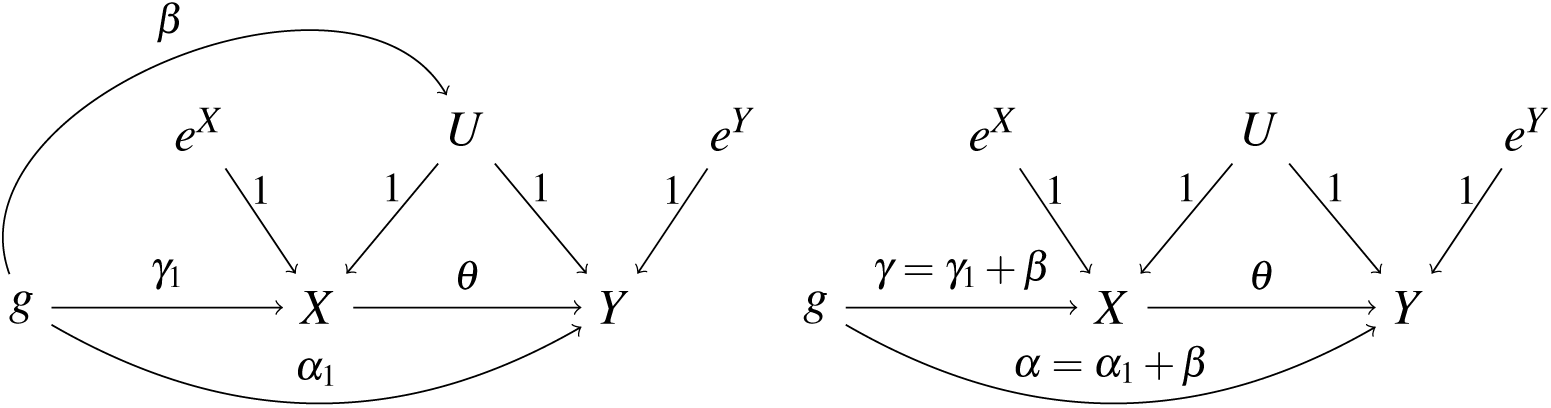
The case where the SNP/IV is directly associated with confounder *U* (left panel), which, by absorbing the effect of *SNP* → *U* into *SNP* → *X* and *SNP* → *Y*, leads to the case where the SNP/IV has a direct effect on *Y* (right panel).

### 2.4 Decision Rules

For each of three methods **CD-Ratio, CD-Egger**, and **CD-GLS**, we would have 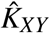 and 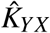. Along with their standard errors (SEs), we construct the confidence intervals at any significance level, say *α*, for *K*_*XY*_ and *K*_*YX*_ respectively as 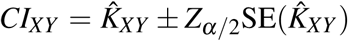 and 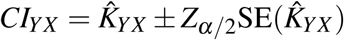. Assuming Key Condition (5) holds, then our decision rules are: (i) If *CI*_*XY*_ is completely outside interval [-1, 1], while *CI*_*YX*_ is inside [-1, 0) or (0, 1], we conclude that the causal direction is from *X* to *Y*; (ii) If *CI*_*YX*_ is completely outside interval [-1, 1], while *CI*_*XY*_ is inside [-1, 0) or (0, 1], we conclude that the causal direction is from *Y* to *X*; (iii) Otherwise, we cannot make a conclusion.

In the above, we have incorporated inferring whether there is indeed any causal relationship between the two traits *X* and *Y*, in contrast to Steiger’s method [25] in simply *assuming* the existence of a causal relationship. As to be shown in simulations, this is advantageous over Steiger’s method and expected to be useful in practice. Our proposal again exploits the use of our formulated estimating target *K*_*YX*_ (or *K*_*XY*_): the sufficient and necessary condition for no causal relationship with *β*_*XY*_ = *β*_*YX*_ = 0 is by definition equivalent to *K*_*YX*_ = *K*_*XY*_ = 0. Hence, we propose using their confidence intervals to determine the existence of a causal relationship.

### 2.5 Goodness-of-Fit Tests

In MR the intercept estimate and its SE from Egger regression can be used as a goodness-of-fit (GOF) test to check the assumptions of the valid IVs [4]; some extensions have also been developed [14]. Accordingly, we can also use our estimate 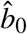 and its CI for such a purpose. In addition, as argued in [6], it is likely better and more powerful to conduct homogeneity testing using Cochran’s *Q* statistics (for the IVW method) and Rucker’s *Q*^*′*^ statistics (for MR-Egger regression). We develop the corresponding three tests in the current context, similar to but different from those for IVW or Egger regression in MR. Using the asymptotic distribution (9), we can construct a chi-squared GOF test statistic for CD-Ratio:

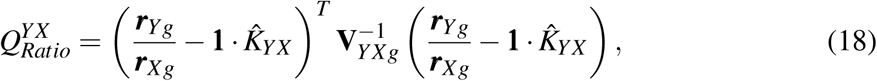

where 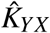 is the CD-Ratio (or any other consistent) estimate. Under the null hypothesis that all SNPs are valid IVs, the (asymptotic) null distribution is 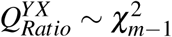, a chi-squared distribution with degrees of freedom *m* − 1, where 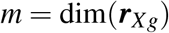 is the number of the SNPs/Ivs being used. Rejection of the null hypothesis may imply that the SNPs may have direct effects on the outcome, i.e. there is horizontal pleiotropy.

Based on the asymptotic distribution (17), we can construct a similar GOF test for CD-Egger after accounting for the possible presence of horizontal pleiotropy:

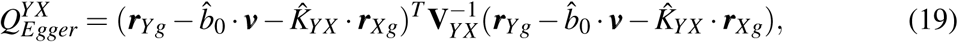

where 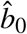and 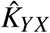 are the CD-Egger estimates of *b*_0_ and *K*_*YX*_ respectively. Under the null hypothesis that the model assumptions hold, the (asymptotic) null distribution is 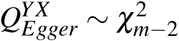. We can construct a similar GOF test based on CD-GLS, which we found performed similarly to 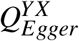 and thus is relegated to the Supplementary. Rejection of the null hypthesis by either test may imply that the SNPs/IVs are possibly correlated with some hidden confounders.

For the other direction *Y* to *X*, similarly we can perform corresponding chi-squared GOF tests with 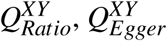 and 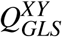.

### 2.6 Further Extensions

#### 2.6.1 Multivariable approach

Analogous to the extenstion of MR to multivariable MR (MV-MR) [8, 9], we extend our CD-Egger method to its multivariable version, denoted **MV-CD-Egger**, to explicitly account for pleiotropic/direct effects through a third (or more) trait. Specifically, we introduce the third trait *A*, which may be an observed confounder for the two traits of interest, *X* and *Y*. When the true causal direction is from *X* to *Y*, the true causal model is:

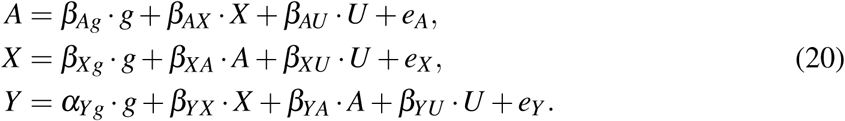

Here *e*_*A*_, *e*_*X*_, *e*_*Y*_ are independent random error terms. In model (20), we allow a general relationship between *A* and *X*, and between *A* and *Y*. When the true causal direction is from *Y* to *X*, we have a corresponding true model. The two possible true models are illustrated in Figure 4.

**Figure 4:**
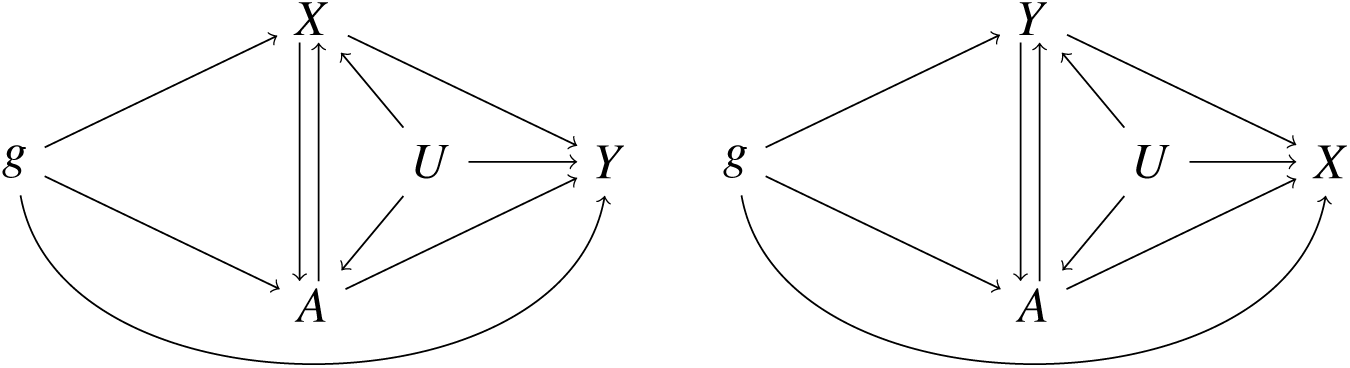
Two possible true models: the left one is for causal direction from *X* to *Y*, and the right one is for *Y* to *X*.

Under the true model of the causal direction from *X* to *Y*, since the IV *g* is independent of the hidden confounder *U* and error term *e*_*Y*_, we can calculate the covariance between *g* and *Y* as:

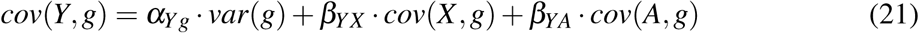

Let 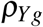 be the correlation between *Y* and *g*, and similarly define 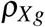 and 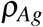, then we have

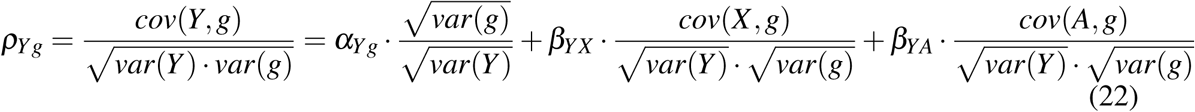

Let 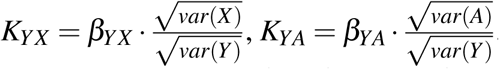 and 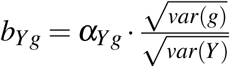. Paralleling with the key condition (5), we assume |*K*_*YX*_ | < 1 and |*K*_*YA*_| < 1. Hence we have

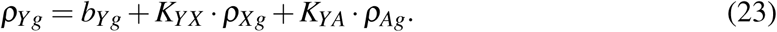

As for CD-Egger, we model the distribution of *b*_*Yg*_ similarly, and apply an iterative method to obtain the maximum likelihood estimates of parameters *K*_*YA*_ and *K*_*YX*_ with their standard errors as shown in section 2.2 in the Supplementary; the only change is to redefine **X** = 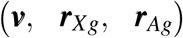. At the end, we construct the normal-based confidence intervals for *K*_*YX*_ and *K*_*XY*_, and apply the same decision rules as before.

#### 2.6.2 Bi-directional causal effects

So far we have restricted our discussion on uni-directional causal effects. Inspired by [48], as suggested by a reviewer, we now briefly discuss an extension of our CD-Ratio method to **bi-CD-Ratio** that works regardless whether there is no, a uni- or bi-directional causal relationship between two traits. In the presence of bi-directional causal effects, as discussed by [48], for the model identifiability, at least one valid IV is required for each trait as show in Fig 5, which is the case we consider here.

**Figure 5:**
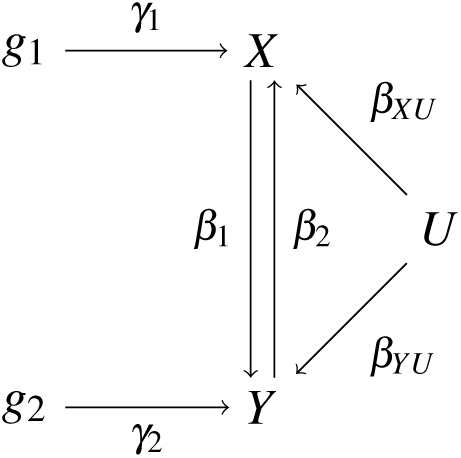
The true model for two traits *X* and *Y* with a bi-directional causal relationship: *g*_1_ and *g*_2_ are valid IVs for *X* and *Y* respectively, and *U* is unobserved confounder.

The true causal model is:

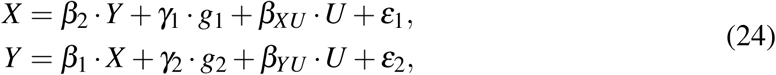

where *ε*_1_ and *ε*_2_ are independent errors for *X* and *Y* respectively. The reduced form of the true model is

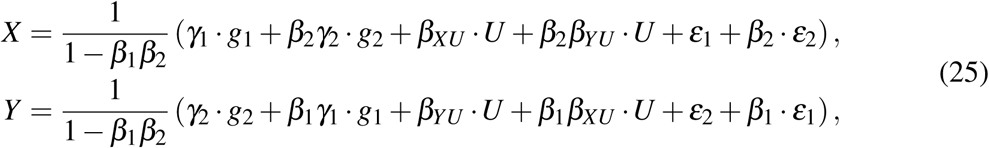

from which it is easy to calculate the correlation between *X* and *g*_1_, denoted by 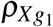, and similarly 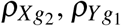 and 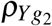 as:

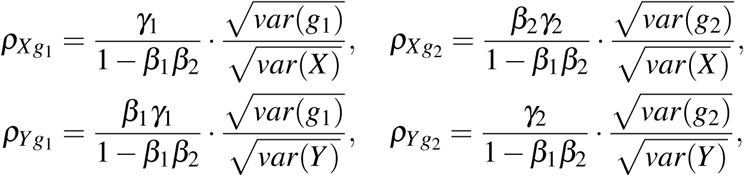

and thus

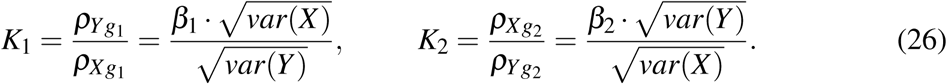

As before, since 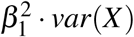 represents a part of the variance in *Y* due to the direct effect of *X* on *Y*, we assume 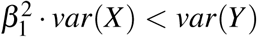, and thus |*K*_1_ | < 1. Similarly we assume |*K*_2_ | < 1. These two assumptions parallel with our key condition (5).

From the GWAS summary data for traits *X* and *Y*, we calculate the sample correlations 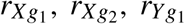 and 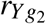. Then we apply our CD-Ratio method with *g*_1_ to estimate *K*_1_ as 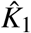 with its standard error 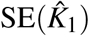, and construct its 100(1 − *α*)% confidence interval 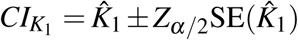. If 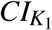 is completely inside [−1, 0) or (0, 1], we conclude that *X* has a causal effect on *Y*; if 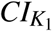 covers 0, then we conclude that *X* has no causal effect on *Y*. Similarly we construct a confidence interval for *K*_2_ and infer whether there is a causal effect from *Y* to *X*. In this way, we may conclude with bi-directional causal effects between *X* and *Y*.

As to be shown in simulations, in the case with valid IVs, both our bi-CD-Ratio and bi-directional MR performed well, being able to distinguish among no, uni- and bi-directional causal relationships between two traits. In contrast, Steiger’s method did not work because it cannot determine whether there is any causal relationship between two traits.

### 2.7 Other Methods

In addition to Steiger’s method, bi-directional MR, also called reciprocal MR, is popular. For the latter, it consists of two steps: first, an MR analysis is applied to infer the causal effect of *X* on *Y*, often using a set of (approximately) independent SNPs signficantly associated with *X* as IVs; second, a separate MR analysis is applied to infer the causal effect of *Y* on *X* using a possibly different set of independent SNPs that are significantly associated with *Y*. The decision rules are similar to ours: 1) if the result is significant in Step 1, but not in Step 2, we conclude that the causal direction is from *X* to *Y*; 2) If the result is significant in Step 2, but not in Step 1, we conclude that the causal direction is from *Y* to *X*; 3) Otherwise, we cannot determine the causal direction. However, an issue with bi-directional MR is that the differing sample sizes, or generally differing statistical power, between the two GWAS for traits *X* and *Y* respectively, would possibly confound the result. An extreme example for an ideal case is that, suppose that both GWAS are sufficiently powered to detect all associated SNPs; it is easy to see that, if there is indeed a causal relation from *X* to *Y* and that *Y* is completely determined by *X* genetically, then all the SNPs associated with *X* would be associated with *Y* too, leading to that both Steps 1 and 2 would give significant results.

We will apply a wide range of MR methods, representing both well-established and new ones, as implemented in R package **TwoSampleMR** for bi-directional MR. These include the inverse-variance–weighted (IVW) method [7], MR-Egger regression [4], the weighted median estimator [5], the weighted mode-based estimation [23, 10], and the Robust Adjusted Profile Score (RAPS) method [61]. To save space, we will only present the results of RAPS with the set-up of overdispersion and a squared-error loss; using other set-ups led to qualitatively similar results. The IVW is perhaps the most popular, requiring the three key assumptions on valid IVs, under which it is most efficient; the other ones either explicitly account for or are robust to horizontal pleiotropy. Since all these MR methods require using independent SNPs, we selected the most significant SNP from each locus if it contained more than one significant SNPs, then applying MR to only independent SNPs in any simulated or real data.

### 2.8 GWAS Data and Reference Panel

To draw inference on the causal direction between two traits using either our proposed methods or bi-directional MR, we only need two independent GWAS summary datasets for the two traits respectively, and a reference panel to estimate LD-matrices for correlated SNPs.

We showcase an application to infer causal directions between LDL and CAD, and between HDL and CAD. The large-scale GWAS summary data for traits LDL/HDL and CAD were drawn from two year 2017 publications [31, 35] with sample sizes up to 295826 for LDL, 316391 for HDL and 336860 for CAD of primary European ancestry respectively. We used the European samples of 489 unrelated individuals in the 1000 Genomes Project [49] as our reference panel to calculate the LD-matrix.

For our proposed methods and Steiger’s methoid, we chose the IVs from the SNPs significantly associated with both traits, denoted as before as *X* and *Y*. Based on [3], we partitioned the whole genome into 1703 approximately LD-independent loci and assigned the significant SNPs (associated with both *X* and *Y*) to one of these 1703 independent loci. We pruned out highly correlated SNPs falling inside each locus in the following two steps. Step 1, we ordered the significant SNPs in each locus according to their combined significance level with *X* and *Y*. Suppose there were *m* significant SNPs with their p-values **p**_*X*_ = (*p*_*X*1_, …, *p*_*Xm*_) and **p**_*Y*_ = (*p*_*Y*1_, …, *p*_*Ym*_) for two traits *X* and *Y* respectively. We sorted **p**_*X*_ and **p**_*Y*_ in an ascending order. For each SNP *i, i* = 1, …, *m*, we obtained the ranks of its p-values *p*_*Xi*_ and *p*_*Yi*_, denoted as *I*_*Xi*_ and *I*_*Yi*_. Then we calculated its rank sum *I*_*i*_ = *I*_*Xi*_ + *I*_*Yi*_, *i* = 1, …, *m*. After that, we reordered the SNPs in the ascending order of *I*_*i*_. Step 2, we pruned out some highly correlated (and significant) SNPs in each locus. We kept SNP 1, and for *i* = 2, …, *m*, if |*cor*(*SNP*_*i*_, *SNP*_*j*_) | *≤* 0.8, *j* = 1, …, *i* −1, we kept it; otherwise we removed it. At the end, we had a set of the significant SNPs with pairwise absolute correlations less than 0.8.

### 2.9 Main Simulation Set-ups

For our main simulations, to mimic the real data analysis for LDL and CAD, in which 22 SNPs from 12 loci were used, we generated simulated data with the same 22 SNPs. We extracted the genotype data of 489 European individuals from the 1000-Genomes Project, then replicated the genotype data of the 22 SNPs from the 489 individuals 200 times, obtaining a sample of size *n* = 97800.

Following [45], in each simulation, we generated the two traits from true models (12-13), where *g* was the vector of the genotype scores of the 22 SNPs, 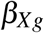 was the (row) vector of the effect sizes of the 22 SNPs on trait *X* with each element independently drawn from a truncated standard normal distribution (truncated at 0.5 to ensure 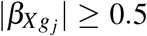), the confounder *U ∼ N*(0, 1); the causal effect size *β*_*YX*_ from *X* to *Y* was fixed at one value, the direct effects of the SNPs on *Y*, the elements of (row) vector *α*, were generated independently 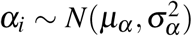; and both error terms were independently from *N*(0, 4). With various values of *β*_*YX*_, *µ*_*α*_ and *σ*_*α*_, we considered a total of 75 simulation set-ups, covering a wide range of situations from no pleiotropy (i.e. *µ*_*α*_ = *σ*_*α*_ = 0) to balanced pleiotropy (i.e. *µ*_*α*_ = 0 and *σ*_*α*_ *>* 0) and direction pleiotropy (i.e. *µ*_*α*_ ≠ 0 and *σ*_*α*_ *>* 0).

After generating each simulated dataset, we performed single SNP-based univariate/marginal analysis for each trait, obtaining the corresponding two GWAS summary datasets. The proposed methods, CD-Ratio, CD-Egger and CD-GLS, were then applied to the GWAS summary data, and for each method one of three possible decisions was made according to the decision rule. For comparison, we also applied MR Steiger’s method to each of the 22 SNPs; we then calculated the proportions of the three possible decisions, denoted by “Steiger-Prop”. In addition, we also pooled together the individual results from each of the 22 SNPs by majority voting to reach a final decision; this method is denoted as “Steiger-MV”.

### 2.10 Data Availability

GWAS summary data for CAD and LDL/HDL are publicly available at http://www.cardiogramplusc4d.org/data-downloads/, http://csg.sph.umich.edu/willer/public/lipids2017, respectively. R code implementing our proposed methods is available at https://github.com/xue-hr/Causal_Direction.

## 3 Results

### 3.1 Real Data Analysis Examples

#### 3.1.1 LDL and CAD

For LDL and CAD, at the significant cutoff of *p* < 5 *×* 10^−8^, out of 1703 (approximately) independent loci [3], we identified 39 significant loci for CAD, 102 for LDL, and 12 common ones for both traits from the two GWAS summary datasets. There were 40 SNPs in these 12 common loci significant for both LDL and CAD, after pruning out highly correlated ones, we obtained 22 SNPs in the 12 loci. We applied MR Steiger’s method and our proposed methods with these 22 SNPs as IVs. For each of the 22 SNPs, we applied MR Steiger’s method to obtain a 95% CI for *Z*_*g*_, as shown in Figure 6(a). If a CI covers 0, no conclusion would be made; otherwise, depending on whether it is in the right or left side of the zero point, we conclude the causal direction of LDL to CAD or the opposite. A problem with single SNP-based analysis is its lower power and thus possibly inconsistent results: as shown in Figure 6(a), although we’d conclude the causal direction of LDL driving CAD based on most of the SNPs, we would fail to conclude based on each of five SNPs while concluding CAD driving LDL on one SNP.

**Figure 6:**
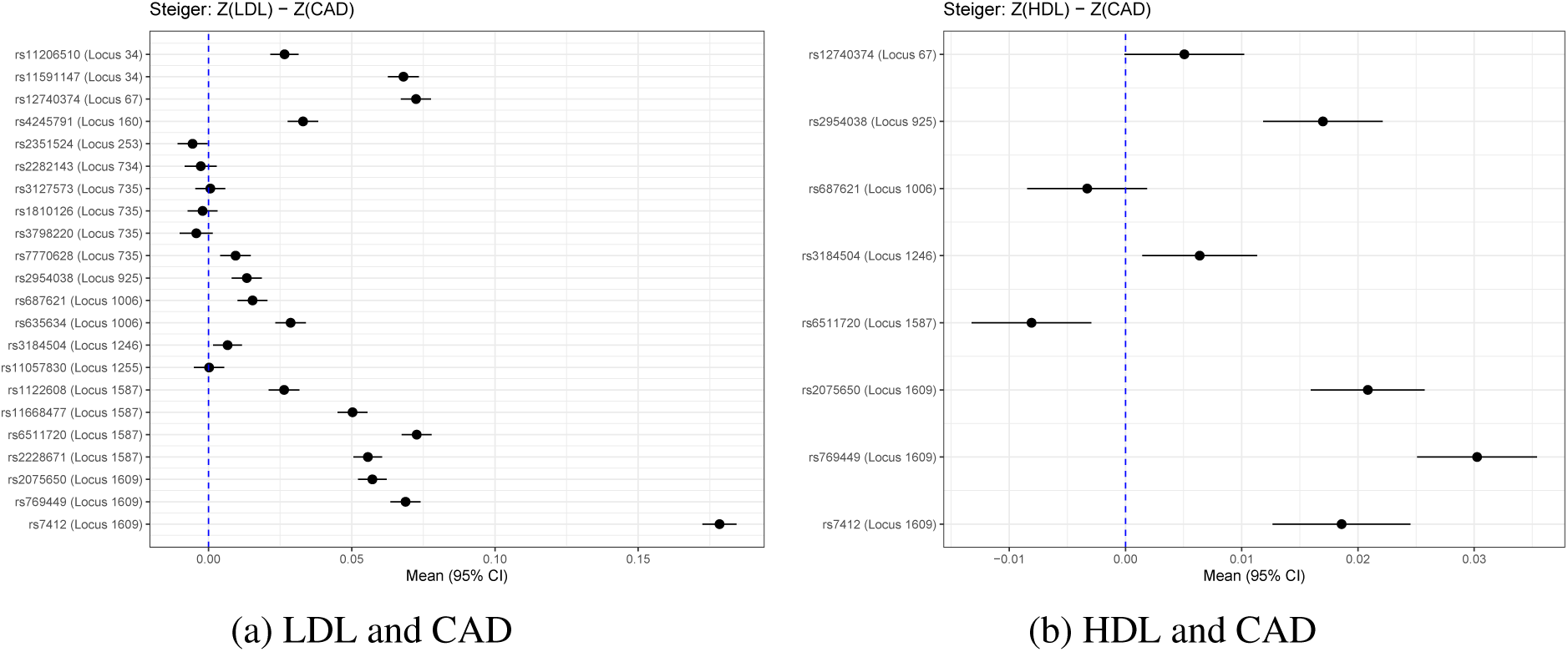
The results by Steiger’s method: 95% CI of the test statistics *Z*_*g*_ for each SNP.

As an alternative, under the Ideal Model, at each locus, we first applied CD-Ratio to all individual SNPs; if there were more than one SNP, we applied CD-Ratio to combine the SNP-specific results at the locus; then we applied CD-Ratio to all 22 SNPs from the 12 loci. For the Pleiotropy Model, we applied CD-Egger and CD-GLS directly to all the SNPs across all the loci. Figure 7 shows the results. First, we emphasize that, perhaps as expected, the single SNP-based results varied with the specific SNP being used as the IV, presumably due to that some SNPs were not valid IVs. For example, for the majority of the SNPs, CD-Ratio would give each a 95% confidence interval (CI) of *K*_*LDL*→*CAD*_ completely inside (0, 1], but a few outside, supporting the causal direction from LDL to CAD. Also note that none of the CIs covered 0, supporting the existence of a causal relationship between the two traits. Second, for the several loci with multiple SNPs, by combining the results over the SNPs, our proposed CD-Ratio improved statistical efficiency by giving shorter locus-specific CIs than the individual SNP-specific ones. Finally, most importantly, under the Ideal Model, after combining over all 22 SNPs from 12 loci, the CD-Ratio method could not conclude on the causal direction, because its CI for *K*_*LDL*→*CAD*_ was inside (0,1] while that for *K*_*CAD*→*LDL*_ covered 1, as given in Table 1. In contrast, after accounting for possible horizontal pleiotropy, our proposed methods CD-Egger and CD-GLS would both conclude the causal direction from LDL to CAD, in agreement with the previous MR analysis and RCT results reported in the literature [19, 13, 12, 44].

**Table 1:**
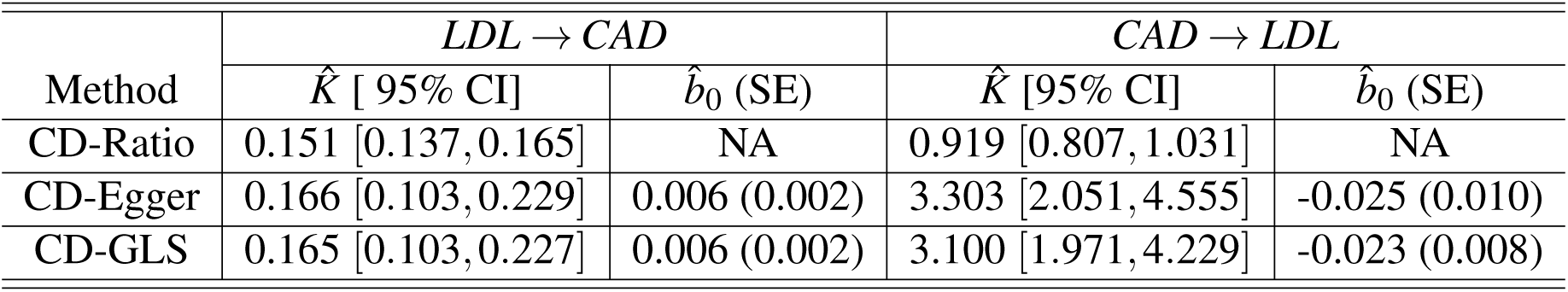
The results for inferring the causal direction between LDL and CAD after combining across all 22 SNPs in 12 loci.

| Method | $LDL \rightarrow CAD$ | | $CAD \rightarrow LDL$ | |
| --- | --- | --- | --- | --- |
| | $\hat{K}$ [95% CI] | $\hat{b}_0$ (SE) | $\hat{K}$ [95% CI] | $\hat{b}_0$ (SE) |
| CD-Ratio | 0.151 [0.137, 0.165] | NA | 0.919 [0.807, 1.031] | NA |
| CD-Egger | 0.166 [0.103, 0.229] | 0.006 (0.002) | 3.303 [2.051, 4.555] | -0.025 (0.010) |
| CD-GLS | 0.165 [0.103, 0.227] | 0.006 (0.002) | 3.100 [1.971, 4.229] | -0.023 (0.008) |

**Figure 7:**
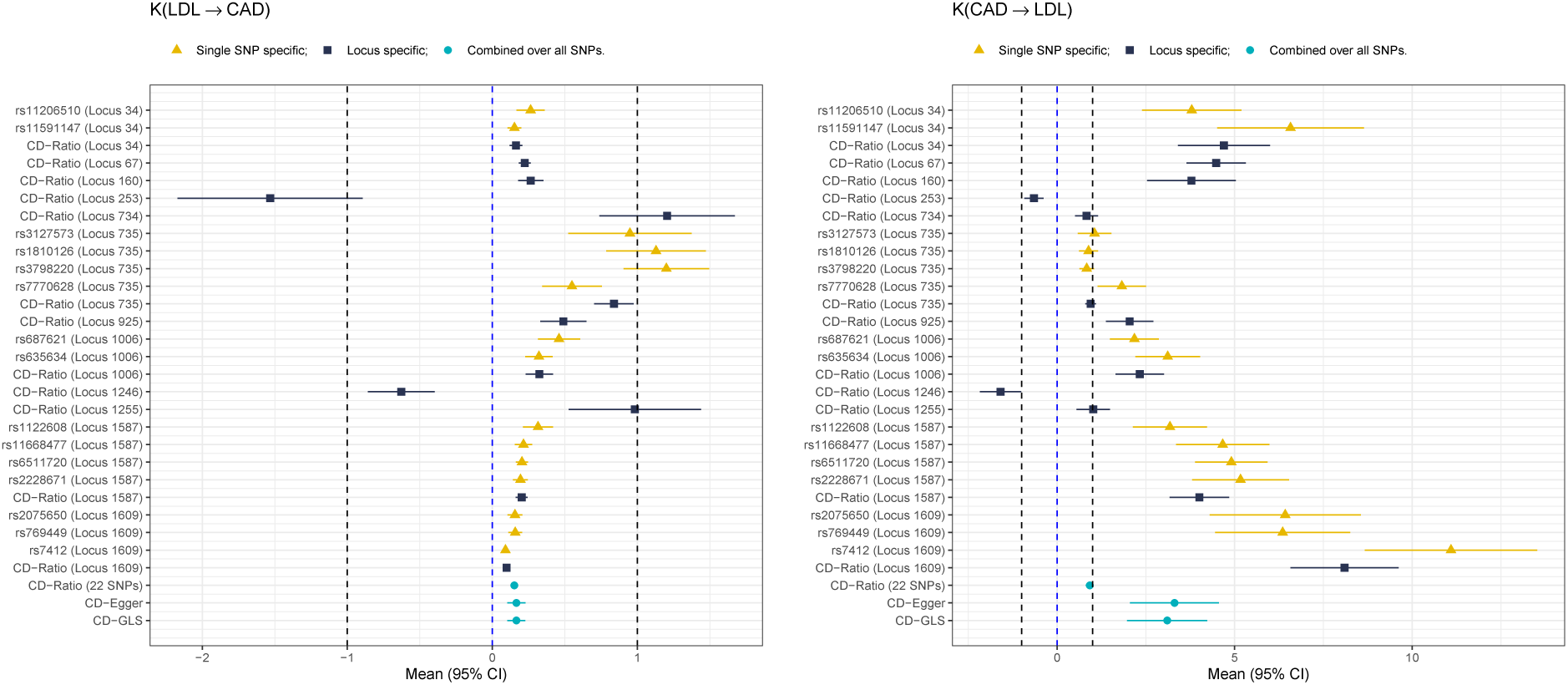
Forest plots for inferring the causal direction between LDL and CAD, showing the differently colored SNP- and locus-specific results and the combined results across all SNPs and loci, respectively.

It is noted that, Locus 253 and Locus 1246 showed a negative (i.e. beneficial) effect of LDL on CAD, in contrary to the known literature. Treating each as an outlier, after removing it, we applied the new methods, and all the three (including CD-Ratio) reached the same conclusion on the causal direction from LDL to CAD; see Supplementary Materials. On the other hand, if we only selected and used one SNP from each locus, CD-Rato could not conclude while both CD-Egger and CD-GLS reached the same conclusion of LDL to CAD, as shown in Supplementary Materials.

It is emphasized that the necessity of accounting for horizontal pleiotropy in this example was supported by the statistically significant non-zero estimates of *b*_0_ in either direction, which was interpreted as representing the mean of the pleiotropic effects; it was notably larger when considering causal direction *CAD* → *LDL* than that in the other direction (Table 1). This latter point was also confirmed by the results of MR Egger regression as to be shown. Furthermore, we conducted the GOF test with *Q*_*Ratio*_ = 337.1, leading to a *p*-value almost 0, rejecting the null hypothesis that the Ideal Model was adequate. The GOF test *Q*_*Ratio*_ = 527.2 for *CAD* → *LDL*, leading to a *p*-value almost 0 too. For CD-Egger, for the direction *LDL* → *CAD* we got *Q*_*Egger*_ = 22.2 and *p*-value=0.329; for the other direction, we obtained *Q*_*Egger*_ = 22.0 and value=0.340. The similar results were obtained from *Q*_*GLS*_, supporting the adequacy of the Pleiotropic Model and the good performance of CD-Egger and CD-GLS.

For comparison, we also applied bi-directional MR. In each of 39 loci significant for CAD, we picked up the most significant SNP (i.e. with the smallest p-value) as the IV, then applied various MR methods to estimate and test the causal effect of CAD on LDL. Similarly, in each of 102 significant loci for LDL, we used the most significant SNP as the IV, then applied MR to estimate and test the causal effect of LDL on CAD. The results are shown in Table 2. We can see that, at the significance level 0.05, IVW, Egger and RAPS (with all 4 combinations of its parameters) all showed some significant causal effects of CAD on LDL, and of LDL on CAD. Although Weighted Median and Weighted Mode concluded that CAD had no causal effect on LDL, and that LDL had causal effect on CAD, it could be due to the low statistical efficiency of the two methods; one evidence is that the two methods gave a negative effect of CAD on LDL, contrary to the known positive association between the two. Overall, across all the methods, there was much stronger statistical evidence to support the causal direction *LDL* → *CAD* than the other direction. Besides, MR-Egger gave an estimate of the mean pleiotropic effect for direction LDL to CAD as -0.003 with standard error 0.003, and -0.054 with standard error 0.028 for CAD to LDL. In summary, as expected, due to use of possibly invalid IVs, bi-directional MR tended to conclude causal effects from *both* directions.

**Table 2:**
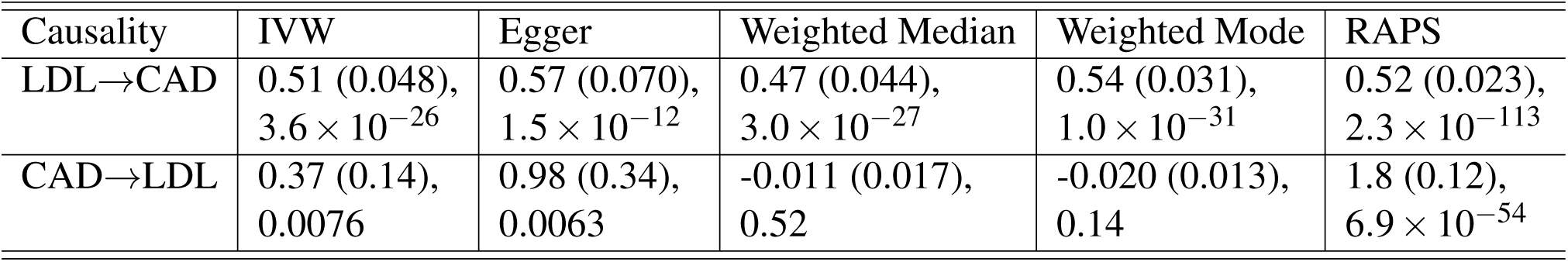
Results of bi-rectional MR for the causal relationship between LDL and CAD by various MR methods. Each cell gives “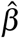 (SE), p-value”.

Finally, since our proposed CD-Egger and CD-GLS suggested the causal direction from LDL to CAD, we checked the key condition (5). Using all the 40 signfiicant SNPs in the 12 loci, we obtained the distribution of *var*(*CAD*)*/var*(*LDL*) ranging from 7.62 to 13.95 with mean 9.44, much larger than any MR 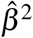 for LDL → CAD listed in Table 2, supporting that condition (5) held here.

#### 3.1.2 HDL and CAD

For HDL and CAD, at the significant cutoff 5 *×* 10^−8^, out of 1703 (approximately) independent loci, we identified 39 significant loci for CAD, 119 for HDL, and 6 common ones for both traits. There were 19 SNPs significant for both HDL and CAD from the 6 common loci, after pruning out highly correlated ones, we obtained 8 SNPs. We applied the MR Steiger’s method and our proposed methods to the 8 SNPs in the 6 common and significant loci. Figure 6(b) shows the results from Steiger’s method, while Figure 8 showing those for CD-Ratio, CD-Egger and CD-GLS.

**Figure 8:**
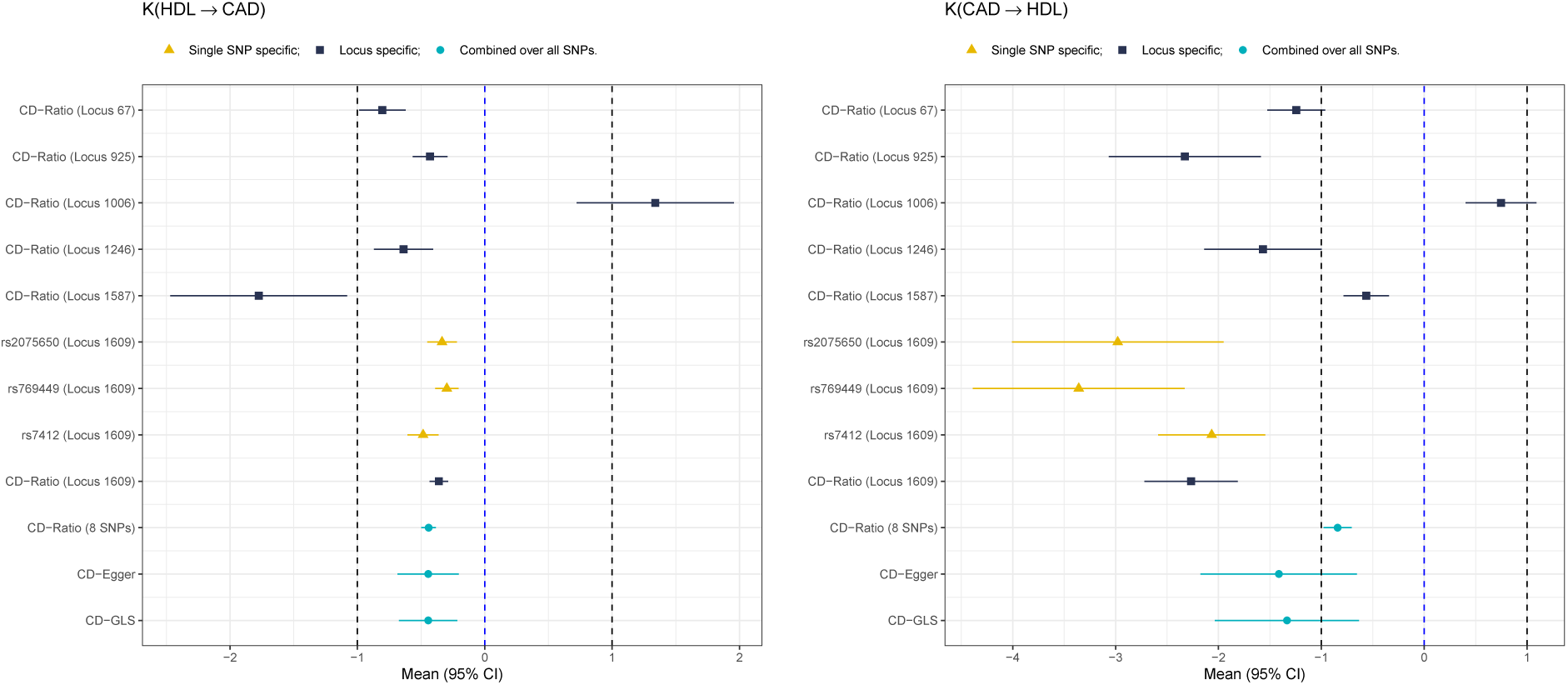
Forest plots for inferring the causal direction between HDL and CAD, showing the differently colored SNP- and locus-specific results and the combined results across all SNPs and loci, respectively.

Again, similar to that for LDL and CAD, the single SNP results varied with different SNPs being used. Combining multiple SNPs in a locus showed gains with higher sttaistical estimation efficiency: for example, at Locus 1609 with 3 SNPs, CD-Ratio gave a shorter CI than the 3 individual SNP-specific ones. As shown in Table 3, we cannot conclude on the causal direction using CD-Ratio under the Ideal Model, because its CIs for *K*_*HDL*→*CAD*_ and *K*_*CAD*→*HDL*_ were both inside [−1, 0). After accounting for possible pleiotropy, both CD-Egger and CD-GLS gave a stronger but not statistically signficiant result of the causal direction from HDL to CAD; both reached the same conclusion: although each gave a CI of *K*_*HDL*→*CAD*_ within (−1, 0), each gave an estimate of *K*_*CAD*→*HDL*_ < −1 but with a CI covering −1, failing to determine the causal direction.

**Table 3:** Results of inferring the causal direction between HDL and CAD after combining across all 8 SNPs in 6 loci.

| Method | $HDL \rightarrow CAD$ | | $CAD \rightarrow HDL$ | |
| --- | --- | --- | --- | --- |
| | $\hat{K}$ [95% CI] | $\hat{b}_0$ (SE) | $\hat{K}$ [95% CI] | $\hat{b}_0$ (SE) |
| CD-Ratio | -0.441 [-0.498, -0.385] | NA | -0.842 [-0.979, -0.706] | NA |
| CD-Egger | -0.445 [-0.685, -0.205] | -0.001 (0.003) | -1.414 [-2.174, -0.654] | -0.001 (0.006) |
| CD-GLS | -0.444 [-0.674, -0.215] | -0.001 (0.003) | -1.334 [-2.035, -0.634] | -0.001 (0.006) |

We also conducted GOF testing. First, we obtained *Q*_*Ratio*_ = 78.1, leading to a *p*-value almost 0, thus rejecting the null hypothesis that the Ideal MR Model for *HDL* → *CAD* was adequate. Similarly, we reached the same conclusion for *CAD* → *HDL* with *Q*_*Ratio*_ = 163.9 and a *p*-value almost 0. Next, for the Pleiotropic Model with CD-Egger, for the direction *HDL* → *CAD*, we obtained *Q*_*Egger*_ = 7.99 and *p*-value=0.239; for the other direction, we had *Q*_*Egger*_ = 8.07 and *p*-value 0.233. Similar results were obtained with CD-GLS. Hence, we concluded the adequacy of the Pleiotropic Model.

We applied bi-directional MR for comparison, results are shown in Table 4. At the significance level 0.05, IVW and RAPS showed significant causal effects of both HDL to CAD and CAD to HDL. MR Egger concluded with a significant causal effect of HDL to CAD but not CAD to HDL, while Weighted Median gave the opposite result. Weighted Mode showed causal effect from CAD to HDL was more significant than from HDL to CAD, though neither was significant at level 0.05. In summary, different bi-directional MR methods gave different and contradictory results.

**Table 4:** Results of bi-rectional MR for the causal relationship between HDL and CAD by various MR methods. Each cell gives “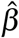 (SE), p-value”.

| Causality | IVW | Egger | Weighted Median | Weighted Mode | RAPS |
| --- | --- | --- | --- | --- | --- |
| HDL→CAD | -0.281(0.058),<br>$1 \times 10^{-6}$ | -0.197(0.096),<br>0.041 | -0.085(0.052),<br>0.100 | -0.083(0.061),<br>0.176 | -0.301(0.053),<br>$1.4 \times 10^{-8}$ |
| CAD→HDL | -0.0823(0.033),<br>0.013 | -0.104(0.084),<br>0.225 | -0.032(0.015),<br>0.029 | -0.027(0.014),<br>0.061 | -0.089(0.033),<br>0.006 |

Finally, given the inconclusive results by our methods, we checked the key condition (5) for both directions. Using all the 19 significant SNPs in the 6 loci, we obtained the distribution of *var*(*CAD*)*/var*(*HDL*) ranging from 8.37 to 11.77 with mean 9.84, much larger than all MR 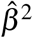 for HDL → CAD; on the other hand, *var*(*HDL*)*/var*(*CAD*) ranged from 0.0849 to 0.1195 with mean 0.1032, larger than any MR 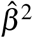 for CAD → HDL. Hence there was evidence supporting condition (5) (or its analog) in either direction.

#### 3.1.3 LDL/HDL and CAD: multivariable approaches

Since MV-MR has been shown to be more robust than MR, in particular in estimating causal effects of LDL and HDL on CAD [63], we also applied bi-directional MV-MR and MV-CD-Egger to infer the causal direction between LDL (or HDL) and CAD while adjusting for HDL (or LDL) as a potential confounder. As shown in Supplementary, our proposed MV-CD-Egger concluded with the causal direction from LDL to CAD, but no causal relationship between HDL and CAD. In contrast, both MV-MR-IVW and MV-MR-Egger concluded with LDL and HDL causal to CAD; in addition, MV-MR-Egger also identified a significant causal effect of CAD on LDL.

### 3.2 Main Simulation Results

#### 3.2.1 Estimation of *K*_*YX*_ : CD-Ratio performed well with no pleiotropy while CD-Egger and CD-GLS always performed well

First we show the simulation results in terms of estimating *K*_*YX*_ by CD-Ratio, CD-Egger and CD-GLS. For each simulation set-up and each method, we show the mean and standard deviation (sd) of its estimates of *K*_*YX*_ from 500 simulated datasets, which are compared to the true value of *K*_*YX*_ and the mean standard error (se) respectively; we would like them to be as close as possible. From Table 5, we can see that CD-Egger and CD-GLS always gave almost unbiased estimates (with good SE estimates too) in all situations, suggesting their correct (and similar) inference on the causal direction. In contrast, while the CD-Ratio estimator was almost unbiased in the absence of pleiotropy, it became more and more biased as the mean pleiotropic effect (i.e. *µ*_*α*_) increased. We obtained similar results and reached the same conclusion from many other simulation set-ups as shown in the Supplementary.

**Table 5:** Simulation results for estimating *K*_*YX*_. The means, standard deviations (sd) and SEs (se) from each method are shown for each set-up based on 500 simulated datasets. *µ*_*α*_ represents the mean of the pleiotropic/direct effects (with standard deviation *σ*_*α*_ = 0.1).

| No pleiotropy |  |  |  |  |  |  |  |  |  |  |
| --- | --- | --- | --- | --- | --- | --- | --- | --- | --- | --- |
| $\mu_\alpha$ | $K_{YX}$ | CD-Ratio | | | CD-Egger | | | CD-GLS | | |
| | | Mean($\hat{K}$) | sd($\hat{K}$) | Mean($se(\hat{K})$) | Mean($\hat{K}$) | sd($\hat{K}$) | Mean($se(\hat{K})$) | Mean($\hat{K}$) | sd($\hat{K}$) | Mean($se(\hat{K})$) |
| 0 | 0 | 0.0001 | 0.0034 | 0.0035 | 0.0001 | 0.0034 | 0.0036 | 0.0001 | 0.0034 | 0.0036 |
| 0 | 0.2257 | 0.2256 | 0.0033 | 0.0036 | 0.2256 | 0.0033 | 0.0036 | 0.2256 | 0.0033 | 0.0036 |
| 0 | -0.2344 | -0.2341 | 0.0032 | 0.0036 | -0.2341 | 0.0032 | 0.0036 | -0.2341 | 0.0032 | 0.0036 |
| Balanced pleiotropy |  |  |  |  |  |  |  |  |  |  |
| $\mu_\alpha$ | $K_{YX}$ | CD-Ratio | | | CD-Egger | | | CD-GLS | | |
| | | Mean($\hat{K}$) | sd($\hat{K}$) | Mean($se(\hat{K})$) | Mean($\hat{K}$) | sd($\hat{K}$) | Mean($se(\hat{K})$) | Mean($\hat{K}$) | sd($\hat{K}$) | Mean($se(\hat{K})$) |
| 0 | 0 | 0.0013 | 0.0563 | 0.0037 | -0.0007 | 0.0431 | 0.0410 | -0.0005 | 0.0436 | 0.0419 |
| 0 | 0.2257 | 0.2181 | 0.0531 | 0.0037 | 0.2201 | 0.0397 | 0.0393 | 0.2203 | 0.0402 | 0.0401 |
| 0 | -0.2344 | -0.2233 | 0.0545 | 0.0037 | -0.2296 | 0.0409 | 0.0408 | -0.2295 | 0.0414 | 0.0416 |
| Directional pleiotropy |  |  |  |  |  |  |  |  |  |  |
| $\mu_\alpha$ | $K_{YX}$ | CD-Ratio | | | CD-Egger | | | CD-GLS | | |
| | | Mean($\hat{K}$) | sd($\hat{K}$) | Mean($se(\hat{K})$) | Mean($\hat{K}$) | sd($\hat{K}$) | Mean($se(\hat{K})$) | Mean($\hat{K}$) | sd($\hat{K}$) | Mean($se(\hat{K})$) |
| 0.2 | 0 | -0.0112 | 0.0538 | 0.0039 | -0.0005 | 0.0386 | 0.0368 | -0.0004 | 0.0392 | 0.0375 |
| 0.4 | 0 | -0.0183 | 0.0448 | 0.0041 | -0.0004 | 0.0306 | 0.0291 | -0.0003 | 0.0311 | 0.0297 |
| 0.6 | 0 | -0.0317 | 0.0357 | 0.0042 | -0.0003 | 0.0240 | 0.0228 | -0.0002 | 0.0244 | 0.0233 |
| 1 | 0 | -0.0506 | 0.0241 | 0.0043 | -0.0002 | 0.0161 | 0.0153 | -0.0002 | 0.0164 | 0.0156 |
| 0.2 | 0.2027 | 0.1898 | 0.0514 | 0.0039 | 0.1987 | 0.0369 | 0.0355 | 0.1989 | 0.0375 | 0.0362 |
| 0.4 | 0.1614 | 0.1438 | 0.0448 | 0.0041 | 0.1593 | 0.0301 | 0.0285 | 0.1595 | 0.0307 | 0.0290 |
| 0.6 | 0.1272 | 0.0944 | 0.0362 | 0.0042 | 0.1261 | 0.024 | 0.0225 | 0.1263 | 0.0244 | 0.0230 |
| 1 | 0.0856 | 0.0313 | 0.0244 | 0.0043 | 0.0851 | 0.0162 | 0.0152 | 0.0852 | 0.0165 | 0.0155 |
| 0.2 | -0.2093 | -0.2172 | 0.0517 | 0.0039 | -0.2061 | 0.037 | 0.0366 | -0.206 | 0.0373 | 0.0373 |
| 0.4 | -0.1649 | -0.1796 | 0.0435 | 0.0041 | -0.1636 | 0.0298 | 0.0290 | -0.1635 | 0.0300 | 0.0296 |
| 0.6 | -0.1290 | -0.1552 | 0.0351 | 0.0042 | -0.1285 | 0.0236 | 0.0228 | -0.1285 | 0.0238 | 0.0233 |
| 1 | -0.0862 | -0.1301 | 0.0238 | 0.0043 | -0.0862 | 0.0159 | 0.0153 | -0.0862 | 0.0161 | 0.0156 |

#### 3.2.2 Inferring causal direction: CD-Ratio outperformed with no pleiotropy while CD-Egger and CD-GLS always outperformed Steiger’s and bidirectional MR methods

We next compare CD-Ratio, CD-Egger and CD-GLS with MR Steiger’s and bi-directional MR in terms of their decisions made on the causal direction. Figure 9 shows some representative results; more are given in the Supplementary. First, we consider the case of no causal relationship between the two traits (i.e. *β*_*YX*_ = *β*_*XY*_ = 0). It was confirmed that MR Steiger’s does not work if there is no causal relationship between two traits: it often incorrectly concluded with one of the two directions; in contrast, our CD-Ratio based on single SNPs overcame this shortcoming if there was no pleiotropy, and performed similarly to Steiger’s method otherwise (see Supplementary Materials). In general, bi-directional MR performed reasonably well, though most of them might make incorrect conclusions. For example, in the presence of balanced pleiotropy, all the five MR methods considered made incorrect decisions at least 10% time. In contrast, our proposed three new methods almost always reached the correct decision. Second, when the causal direction was from *X* to *Y*, our methods CD-Egger and CD-GLS had high (and similar) power to detect it across all the scenarios, regardless of the presence of balanced or directional pleiotropy. As expected, CD-Ratio performed well only when there was no pleiotropy or only balanced pleiotropy, but it completely lost power in the presence of directional pleiotropy. Steiger’s method was high powered except that it would often conclude with the incorrect causal direction in the presence of directional pleiotropy. On the other hand, all the five MR methods were low powered, and would often conclude with the incorrect causal direction in the presence of directional pleiotropy. Perhaps it is surprising to see low power of MR even in the ideal case of no pleiotropy. The reason is that bi-directional MR tends to detect both directions being significant (due to the use of invalid IVs), leading to no final conclusions by the decision rules (since we assume either no or a uni-directional causal relationship); this is confirmed by Figure 10, in which the significant results in either direction are shown separately (without being combined by the decision rules).

**Figure 9:**
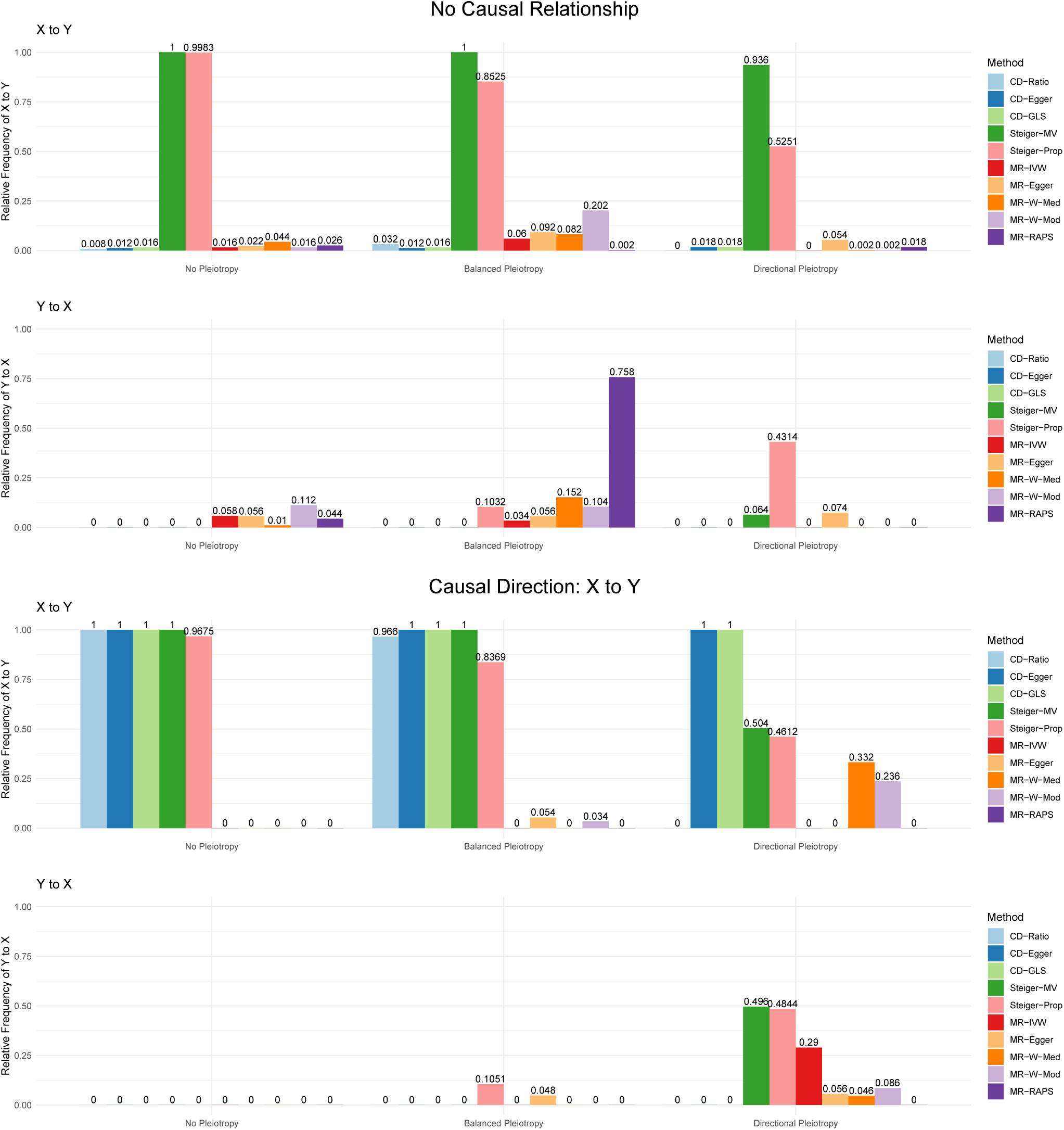
Relative frequencies of the causal directions reached by each method (from 500 simulations in each set-up) when there is no causal relatuonship (top two panels) or a causal direction is from *X* to *Y* (*β*_*YX*_ = 0.2; bottom two) with no pleiotropy, balanced pleiotropy (*µ*_*α*_ = 0, *σ*_*µ*_ = 0.1) and directional pleiotropy ((*µ*_*α*_ = 0.4, *σ*_*µ*_ = 0.1) respectively.

**Figure 10:**
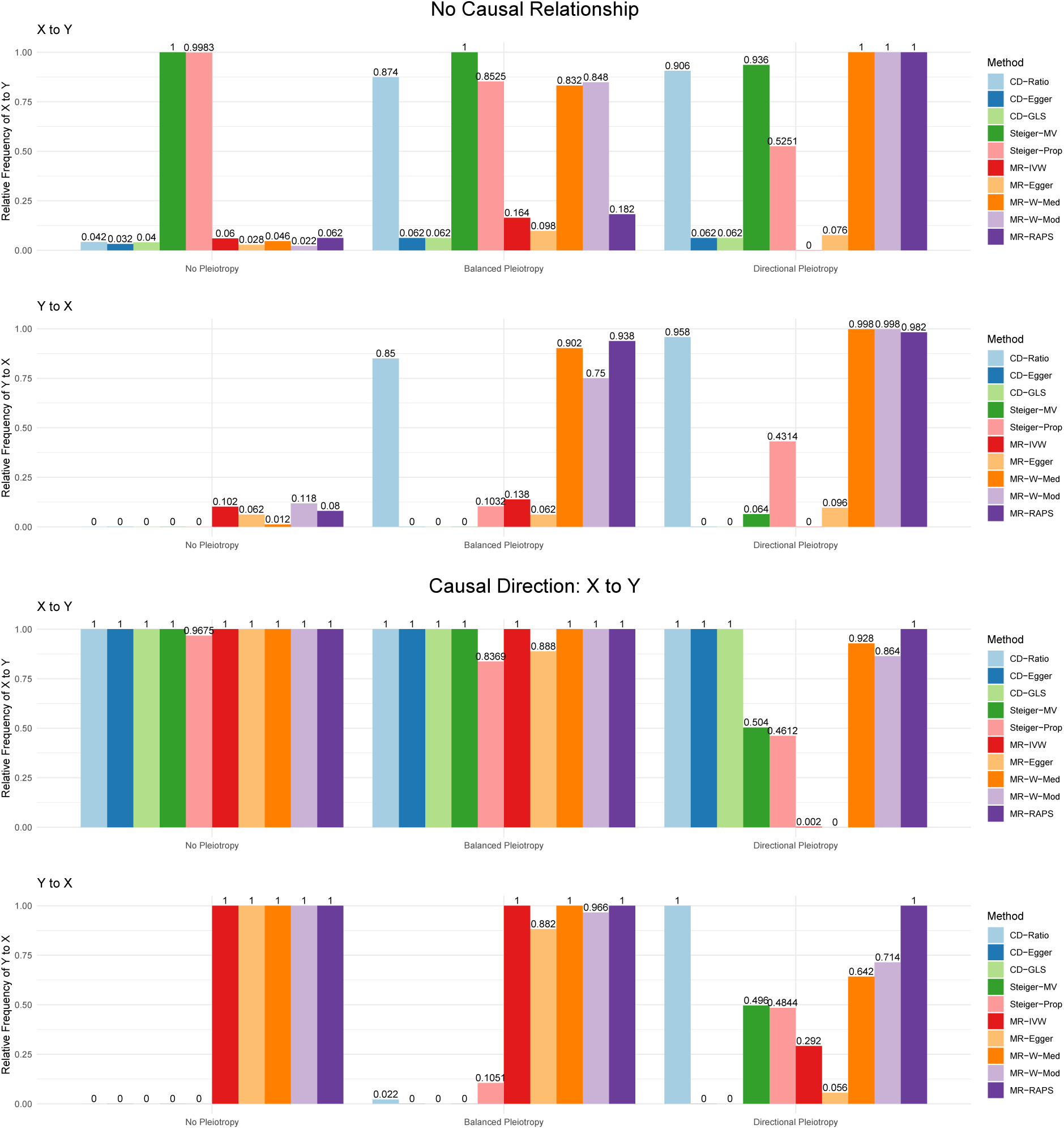
Relative frequencies of having significant results in each direction, in contrast to that of the final conclusions shown in Figure 9.

#### 3.2.3 The GOF tests performed well

In Supplementary Materials, we show the results of our proposed GOF tests: all performed well with controlled type I error rates and high power in our simulations.

### 3.3 Secondary Simulation Results

#### 3.3.1 Bi-CD-Ratio could detect bi-directional causal relationships

In Supplementary Materials, we also show the results for inferring no, uni- and bi-directional relationships between two traits with valid IVs. It is confirmed that our proposed bi-CD-Ratio and (bi-directional) MR-Wald-Ratio performed well, while Steiger’s method did not work due to its inability to determine whether there is any causal relationship between two traits as discussed earlier.

#### 3.3.2 CD-Ratio could perform in TWAS with relatively small sample sizes

We also did a simulation study to investigate how the asymptotic theory for the sample correlations and their ratios performs with smaller sample sizes as typical with molecular endophenotypes as in TWAS. We generated simulated data under realistic set-ups mimicking real eQTL datasets with sample sizes from around two hundreds to about a thousand, much smaller than those of typical GWAS as used in our main simulations. We also considered the presence of some highly correlated SNPs/IVs. As shown in Supplementary section 8, our proposed CD-ratio performed very well: it might be a bit conservative with empirical type I error rates smaller than the nominal level when the sample size was small at a few hundreds; but as the sample size increased to a few thousands, its type I error rates approached the nominal level. Importantly, it was confirmed that by combining the results across the SNPs, even including some highly correlated ones, CD-Ratio improved the statistical power over that based on single SNPs.

#### 3.3.3 Selection bias

We did simulations to study possible selection bias due to the double use of the same sample for SNP/IV selection and estimation. The simulation setups are the same as described in Section 2.9. We used 22 SNPs to generate simulated data, and in Section 3.2.1 we applied our proposed three methods to all 22 SNPs with the results shown in Table 5. Now for each simulated dataset, we used a p-value cutoff to select exactly *N* SNPs that were significantly associated with both *X* and *Y*, then we applied a method with these *N* selected SNPs. We tried with *N* = 10, 15 or 22; when *N* = 22, we used all 22 SNPs as before. Tables S41, S42 and S43 in the Supplementary show the simulation results with no pleiotropy, balanced pleiotropy, and directional pleiotropy respectively. We summarize the results below. First, when there is no pleiotropy, all three methods gave almost unbiased estimates of *K*_*YX*_ with any of the three values of *N*, suggesting no or negligible selection bias. Second, with balanced pleiotropy, there were small biases. Third, with directional pleiotropy, there could be large biases. However, interestingly, under the null hypothesis of *K*_*YX*_ = 0, CD-Egger and CD-GLS yielded only small biases, leading to barely any inflation of their Type I error rates; see Figures S29 to S34 in the Supplementary.

## 4 Conclusions and Discussion

We have proposed several extensions to the increasingly popular MR Steiger’s method to infer the causal direction between two traits. MR Steiger’s method is based on using a single SNP as a valid IV; if there is strong biological knowledge to justify the use of the SNP as a valid IV, it is both useful and simple to apply the method. However, in practice, due to incomplete knowledge, one often just uses genome-wide significant SNPs as IVs, in which case there are several potential problems. Some of the SNPs are likely invalid IVs due to wide-spread horizontal pleiotropy. In addition, the results may be low-powered and inconsistent, each critically depending on the SNP being used. Our proposed multiple SNP-based approaches would alleviate the above two problems. First, by using multiple SNPs, we allow and thus explicitly model possible pleiotropic effects, leading to two new methods, CD-Egger and CD-GLS, that are robust to and more powerful in the presence of pleiotropy. Second, combining the results over multiple SNPs is both more robust and more powerful than a single SNP-based method. In applications to inferring/confirming causal directions between molecular endophenotypes and complex diseases or quantitative traits, such as in TWAS, one would typically use multiple SNPs in LD from a local region to predict an endophenotype (e.g. a gene’s expression level) with relatively small sample sizes; in these applications, it may be necessary to apply a multiple correlated SNPs-based method like ours to achieve as high power as possible. Third, we can apply some goodness-of-fit tests to check the modeling assumptions. Furthermore, our methods can simultaneously infer both the existence and (if so) the direction of a causal relationship between two traits. For this reason, even for single SNPs, our CD-Ratio is a useful alternative to Steiger’s method because of the former’s ability to avoid incorrect conclusions of a causal direction when there is no causal relationship (and no pleiotropy). In addition, our methods have some advantages, i.e. higher statistical power and lower false positives, over, and are thus useful alternatives to, multiple SNP-based bi-directional MR. These advantages over bi-directional MR are perhaps due to that our proposed methods take advantage of and thus exploit some specific relationships between the Pearson correlations between each SNP/IV and the two traits implicitly implied by the true causal models. These points were confirmed in both extensive simulations and an application to infer the causal direction between LDL cholesterol and CAD. In particular, bi-directional MR was found to be non-robust for inferring causal directions in the presence of invalid IVs even when equipped with a robust MR method. In summary, our proposed methods are expected to be useful in the toolbox for causal inference, for which application of multiple complementary methods, called triangulation [30], is strongly advocated.

Our methods have some limitations. First and foremost, our methods are based on the standard asymptotic theory for sample correlations under the assumption of a large sample size (say *n*) and a small number of SNPs/IVs (say *m*). It has some important implications. Although including more relevant SNPs/IVs is expected to improve the statistical estimation efficiency and power in the ideal case, in practice there are some arguments against doing so: 1) including too many SNPs, i.e. making *m* large relative to *n*, the standard asymptotic theory may break down, leading to incorrect inference; 2) in particular, including some highly correlated SNPs may gain little while being at the risk of inducing a covariance matrix nearly singular, leading to bad estimates and inference with numerical problems; 3) our methods require estimating a large covariance matrix of dimension in the order of *m*^2^ *× m*^2^ for a sample correlation matrix in the order of *m × m*, demanding too much computer memory and computing time even if *m* is only in hundreds or thousands; 4) including more SNPs will increase the likelihood of violating IV assumptions; 5) it might be harder to accurately impute a high LD between two SNPs for GWAS summary data based on a not-too-large reference panel. For the above reasons, in our applications we pruned out those significant SNPs with pairwise correlations larger than 0.8. We also emphasize that we require the selected SNPs/IVs to be statistically significant for both traits. One reason is to similar to avoiding biases due to weak IVs in MR. Our methods are based on the ratios of sample correlations, say 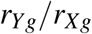. If the mean of the denominator 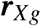 is 0, regardless of whether the normality (approximately) holds for each sample correlation, the ratio cannot be well approximated by a normal approximation; our methods are based on the asymptotic joint normal distribution for the correlation ratios. Second, we have mainly restricted to inferring uni-directional causal relationships. Although we have outlined an extension of our CD-Ratio method (along with bi-directional MR) to the case with bi-directional causal relationships, and shown some promising simulation results under the ideal situation with valid IVs, it remains to see how to extend and apply the methods in practice without knowing which IVs are valid, in which case there may be issues of model non-identifiability [48]. We note by passing that [48] considered a mediating analysis of how to estimate direct/indirect effects given the presence of a bi-directional causal relationship, differing from our task of estimating what causal direction if any. Third, as often done in applications with MR, we have used the same two GWAS samples both for selecting SNPs as IVs and for subsequent estimation and inference. As expected, this double use of the data could lead to selection bias. It was confirmed in our simulations in the presence of directional pleiotropy, though no or only small biases were observed in cases of no or only balanced pleiotropy, possibly due to our recommendation of selecting SNPs/IVs that are significant for both traits, alleviating the problem as compared to selecting SNPs/IVs based on only one trait as commonly adopted in MR. Nevertheless, this issue needs to be further studied; if possible, we may use a third independent GWAS sample for such selection to avoid selection bias as in the three-sample MR design [62]. Fourth, it is noted that our proposed methods are developed in the framework of two-sample MR, which is more common and less restrictive than the one-sample set-up (where both traits and SNPs are obtained from the same sample). Nevertheless, it will be usefull to extend our methods to the one-sample set-up. In particular, such methods may be usedful for mediation analysis [54, 42], in which methods to determine causal directions (i.e. to determine between two traits which is the mediator and which is the outcome) lack and are urgently needed while a bi-directional mediation analysis would be problematic as recently reported [32]. In addition, we have not compared with structure equation modeling [1], Bayesian network [28] and latent causal variable [37] approaches, mainly because of their lack of significance testing. Finally, we have pointed out a strong connection between our proposed CD-Egger method and MR Egger regression in dealing with horizontal pleiotropy. Taking advantage of this connection, one may be able to develop other extensions as developed for MR [9, 40, 51, 39, 45] in addition to those used here (e.g. weighted median and mode methods) for inference of causal direction. These interesting topics warrant future investigation.

## Supplementary Materials

In a Supplementary File we provide more details on theoretical derivations and numerical results.

## Supporting information

Supplementary_MRCD

## Acknowledgements

We thank the reviewers for many helpful and insightful comments and suggestions. This work was supported by NIH grants R01AG065636, R01HL116720, R01GM113250 and R01GM126002, and by the Minnesota Supercomputing Institute at the University of Minnesota.

