## Supplementary_MRCD for "Inferring Causal Direction Between Two Traits in the Presence of Horizontal Pleiotropy with GWAS Summary Data"

September 1, 2020

### 1 Asymptotic Covariance Matrix of $\boldsymbol{r}_{Xg}$ and $\boldsymbol{r}_{Yg}$

In the main text when we use multiple correlated SNPs to estimate correlation ratio, we use the asymptotic distributions of $\boldsymbol{r}_{Xg}$ and $\boldsymbol{r}_{Yg}$:

$$\sqrt{n_X}\cdot(\boldsymbol{r}_{Xg}-\boldsymbol{\rho}_{Xg})\to N(0,\mathbf{V}_X),\quad \sqrt{n_Y}\cdot(\boldsymbol{r}_{Yg}-\boldsymbol{\rho}_{Yg})\to N(0,\mathbf{V}_Y) \tag{1}$$

Here we show the calculation of asymptotic covariance matrix $\mathbf{V}_X$, and $\mathbf{V}_Y$ can be calculated similarly.

Suppose we have $m$ SNPs denoted by $g_1,g_2,\cdots,g_m$ with trait $X$. From GWAS summary statistics, we can get sample correlations $\boldsymbol{r}_{Xg}\in\mathbb{R}^{m\times 1}$ between $g$'s and $X$. From a reference panel like 1000-Genomes Project with $n$ individuals, we get individual level genotype data for $g_i$ as $\boldsymbol{g}_i=(g_{i1},g_{i2},\cdots,g_{in})^T$, $i=1,2,\cdots,m$. Because we only care about correlations, we can assume $g$'s and $X$ are standardized to have mean 0 and variance 1 in both GWAS data and reference panel. Denote $\mathbf{G}=(\boldsymbol{g}_1,\boldsymbol{g}_2,\cdots\boldsymbol{g}_m)\in\mathbb{R}^{n\times m}$. We can estimate the correlation matrix of $g_1,g_2,\cdots,g_m$, denoted by $\boldsymbol{\Sigma}=\mathbf{G}^T\mathbf{G}/n\in\mathbb{R}^{m\times m}$, as an approximation of sample correlation matrix of $g_1,g_2,\cdots,g_m$ calculated from the GWAS individual level data of $X$. So we can get the sample correlation matrix $\hat{\mathbf{P}}\in\mathbb{R}^{(m+1)\times(m+1)}$ of $x=(g_1,g_2,\cdots,g_m,X)^T$:

$$\hat{\mathbf{P}}=\begin{pmatrix}\boldsymbol{\Sigma} & \boldsymbol{r}_{Xg}\\ \boldsymbol{r}_{Xg}^T & 1\end{pmatrix} \tag{2}$$

And denote $\mathbf{P}\in\mathbb{R}^{(m+1)\times(m+1)}$ is the true correlation matrix of $g_1,g_2,\cdots,g_m$ and $X$. Then we can apply Theorem 2 from [1] to get:

$$\sqrt{n_X}\mathrm{vec}(\hat{\mathbf{P}}-\mathbf{P})\xrightarrow{D}N(0,\mathbf{A}) \tag{3}$$

Here $\mathbf{A}=[\mathbf{I}-\mathbf{M}_s(\mathbf{I}\otimes\mathbf{P})\mathbf{M}_d](\boldsymbol{\Lambda}^{-1/2}\otimes\boldsymbol{\Lambda}^{-1/2})\mathbf{V}(\boldsymbol{\Lambda}^{-1/2}\otimes\boldsymbol{\Lambda}^{-1/2})[\mathbf{I}-\mathbf{M}_d(\mathbf{I}\otimes\mathbf{P})\mathbf{M}_s]$. Here "$\otimes$" is Kronecker product of two matrices.

Matrices $\mathbf{M_s},\mathbf{M_d}\in\mathbb{R}^{(m+1)^2\times(m+1)^2}$ can be calculated from equations (2.9) and (2.13) from [1]. We can plug in $\hat{\mathbf{P}}$ to replace $\mathbf{P}$. And because we assume $g$'s and $X$ are standardized to have variance 1, $\boldsymbol{\Lambda}$ is identity matrix. Then we need to calculate $\mathbf{V}$.

Using equation (3.6) in [1], because we standardized $g$'s and $X$, the covariance matrix is the same as correlation matrix, we have:

$$\mathbf{V}=E[(x-\mu)(x-\mu)^T\otimes(x-\mu)(x-\mu)^T]-(\mathrm{vec}\mathbf{P})(\mathrm{vec}\mathbf{P})^T \tag{4}$$

Here $x=(g_1,g_2,\cdots,g_m,X)^T$ and $\mu=E(x)$. So we need $E[(x-\mu)(x-\mu)^T\otimes(x-\mu)(x-\mu)^T]$.

For $n$ individuals in the reference panel, denote their unobserved $X$ values as $\boldsymbol{X}=(X_1,X_2,\cdots,X_n)^T$, again $\boldsymbol{X}$ is standardized with mean 0 and variance 1. We fit a joint linear model of $X$ on $g_1,g_2,\cdots,g_m$:

$$X_i=\beta_1\cdot g_{1i}+\beta_2\cdot g_{2i}+\cdots+\beta_m\cdot g_{mi}+\varepsilon_i \tag{5}$$

Here $\varepsilon_i\sim N(0,\sigma^2)$ is random error. We have $\hat{\boldsymbol{\beta}}=(\mathbf{G}^T\mathbf{G})^{-1}\mathbf{G}\boldsymbol{X}=(\mathbf{G}^T\mathbf{G}/n)^{-1}\mathbf{G}\boldsymbol{X}/n$. Here $\mathbf{G}^T\mathbf{G}/n=\boldsymbol{\Sigma}$, and $\mathbf{G}\boldsymbol{X}/n$ are correlations of $g$'s with $X$ which could be replaced with $\boldsymbol{r}_{Xg}$ from GWAS summary statistics. So we get $\hat{\boldsymbol{\beta}}=\boldsymbol{\Sigma}^{-1}\boldsymbol{r}_{Xg}$, and $\hat{\sigma}^2=||\boldsymbol{X}-\mathbf{G}\hat{\boldsymbol{\beta}}||^2/n=1-\boldsymbol{r}_{Xg}^T\boldsymbol{\Sigma}^{-1}\boldsymbol{r}_{Xg}$. So we can approximate $X_i=\hat{\beta}_1\cdot g_{1i}+\hat{\beta}_2\cdot g_{2i}+\cdots+\hat{\beta}_{mi}\cdot g_m+\hat{\varepsilon}_i$ with independently generated $\hat{\varepsilon}_i\sim N(0,\hat{\sigma}^2)$. With this representation of $X_i$ in the reference panel, we can calculate the sample version of $E[(x-\mu)(x-\mu)^T\otimes(x-\mu)(x-\mu)^T]$ as its estimate. Thus, we get $\mathbf{V}$.

In summary we can estimate $\mathbf{A}$ in (3) which is the asymptotic covariance matrix of $\hat{\mathbf{P}}$, by extracting elements in $\mathbf{vec}(\hat{\mathbf{P}})$ corresponding to $\boldsymbol{r}_{Xg}$, we can get the asymptotic covariance matrix $\mathbf{V}_X$ of $\boldsymbol{r}_{Xg}$.

### 2 Details of CD-Egger and CD-GLS

#### 2.1 Distribution of $\boldsymbol{b}_{Yg}$

In the main text, when there are direct effects from $g$'s to $Y$, we got

$$\rho_{Yg} = (\alpha_{Yg} + \beta_{YX} \cdot \beta_{Xg}) \cdot \frac{\sqrt{var(g)}}{\sqrt{var(Y)}} = \alpha_{Yg} \cdot \frac{\sqrt{var(g)}}{\sqrt{var(Y)}} + K_{YX} \cdot \rho_{Xg} = b_{Yg} + K_{YX} \cdot \rho_{Xg}. \tag{6}$$

Here $\alpha_{Yg}$'s are the direct effects of $g$'s to $Y$ in the *marginal model*. $g$'s are standardized to have mean 0 and variance 1, and the covariance matrix of $g$'s is $\mathbf{\Sigma}$. If we assume the following joint model:

$$Y = \beta_{Y0} + \beta_{YX} \cdot X + \alpha^{(0)}_{Yg_1} \cdot g_1 + \cdots + \alpha^{(0)}_{Yg_m} \cdot g_m + U + \varepsilon_Y \tag{7}$$

Here $\alpha^{(0)}_{Yg_i}$'s are direct effects in the *joint model*, and assume $\alpha^{(0)}_{Yg_i}$'s are i.i.d. $N(b_0, \sigma_0^2)$. Then

$$\begin{aligned}
\boldsymbol{\rho}_{Yg} =& \frac{1}{\sqrt{var(Y)}} \begin{pmatrix} cov(Y,g_1) \\ \vdots \\ cov(Y,g_m) \end{pmatrix} = \frac{1}{\sqrt{var(Y)}} \left( \beta_{YX} \cdot \begin{pmatrix} cov(X,g_1) \\ \vdots \\ cov(X,g_m) \end{pmatrix} + \mathbf{\Sigma} \begin{pmatrix} \alpha^{(0)}_{Yg_1} \\ \vdots \\ \alpha^{(0)}_{Yg_m} \end{pmatrix} \right) \\
=& \beta_{YX} \frac{\sqrt{var(X)}}{\sqrt{var(Y)}} \begin{pmatrix} \frac{cov(X,g_1)}{\sqrt{var(X)}} \\ \vdots \\ \frac{cov(X,g_m)}{\sqrt{var(X)}} \end{pmatrix} + \frac{\mathbf{\Sigma}}{\sqrt{var(Y)}} \begin{pmatrix} \alpha^{(0)}_{Yg_1} \\ \vdots \\ \alpha^{(0)}_{Yg_m} \end{pmatrix} \\
=& K_{YX} \cdot \boldsymbol{\rho}_{Xg} + \frac{\mathbf{\Sigma}}{\sqrt{var(Y)}} \begin{pmatrix} \alpha^{(0)}_{Yg_1} \\ \vdots \\ \alpha^{(0)}_{Yg_m} \end{pmatrix}
\end{aligned} \tag{8}$$

So we have

$$\boldsymbol{b}_{Yg} = \frac{\mathbf{\Sigma}}{\sqrt{var(Y)}} \begin{pmatrix} \alpha^{(0)}_{Yg_1} \\ \vdots \\ \alpha^{(0)}_{Yg_m} \end{pmatrix} \sim N\left( \frac{b_0 \cdot \mathbf{\Sigma 1}}{\sqrt{var(Y)}}, \frac{\sigma_0^2 \mathbf{\Sigma}^2}{var(Y)} \right) \tag{9}$$

#### 2.2 CD-Egger: Estimation of $(b_0, K_{YX})$ and $\sigma_0^2$

For CD-Egger, we have:

$$\boldsymbol{r}_{Yg} = b_0 \cdot \boldsymbol{v} + K_{YX} \cdot \boldsymbol{r}_{Xg} + \boldsymbol{\varepsilon}, \boldsymbol{\varepsilon} \sim N(0, \frac{\mathbf{V}_{Yg}}{n_Y} + \sigma_0^2 \mathbf{\Sigma}^2 = \mathbf{V}_{YX}) \tag{10}$$

We use a iterative algorithm to estimate $(b_0, K_{YX})$ and $\sigma_0^2$. Set initial $\sigma_0^2 = 0$.

Step (1): Given $\sigma_0^2$, we can get $\mathbf{V}_{YX} = \frac{\mathbf{V}_{Yg}}{n_Y} + \sigma_0^2 \mathbf{\Sigma}^2$. Then estimating $(b_0, K_{YX})$ is a Generalize Least Square problem with design matrix $\mathbf{X}$ and covariance matrix of errors $\mathbf{\Omega}$:

$$\begin{aligned}
\mathbf{X} &= \begin{pmatrix} \boldsymbol{v} & \boldsymbol{r}_{Xg} \end{pmatrix} = \begin{pmatrix} v_1 & r_{Xg_1} \\ \vdots & \vdots \\ v_m & r_{Xg_m} \end{pmatrix} \\
\mathbf{\Omega} &= \mathbf{V}_{YX}
\end{aligned} \tag{11}$$

So we can get the estimation $(\hat{b}_0, \hat{K}_{YX})$ and their covariance matrix:

$$\begin{aligned} \begin{pmatrix} \hat{b}_0 \\ \hat{K}_{YX} \end{pmatrix} &= (\mathbf{X}^T \boldsymbol{\Omega}^{-1} \mathbf{X})^{-1} \mathbf{X}^T \boldsymbol{\Omega}^{-1} \boldsymbol{r}_{Yg} \\ cov \begin{pmatrix} \hat{b}_0 \\ \hat{K}_{YX} \end{pmatrix} &= (\mathbf{X}^T \boldsymbol{\Omega}^{-1} \mathbf{X})^{-1} \end{aligned} \tag{12}$$

Step (2): Given $(\hat{b}_0, \hat{K}_{YX})$, plug them in equation (10), denoting

$$\boldsymbol{x}_{YX} = \boldsymbol{r}_{Yg} - \hat{b}_0 \cdot \boldsymbol{v} - \hat{K}_{YX} \cdot \boldsymbol{r}_{Xg} \tag{13}$$

We get

$$\boldsymbol{\Sigma}^{-1} \boldsymbol{x}_{YX} \sim N(0, \frac{\boldsymbol{\Sigma}^{-1} \mathbf{V}_{Yg} \boldsymbol{\Sigma}^{-1}}{n_Y} + \sigma_0^2 \mathbf{I}) \tag{14}$$

Do the eigenvalue decomposition of

$$\frac{\boldsymbol{\Sigma}^{-1} \mathbf{V}_{Yg} \boldsymbol{\Sigma}^{-1}}{n_Y} = \mathbf{Q}^T \begin{pmatrix} \lambda_1 & \dots & 0 \\ \vdots & \ddots & \vdots \\ 0 & \dots & \lambda_m \end{pmatrix} \mathbf{Q} \tag{15}$$

Here $\mathbf{Q}$ is orthonormal, $\lambda_1, \cdots, \lambda_m$ are eigen-values. Denote $\mathbf{Q}\boldsymbol{\Sigma}^{-1}\boldsymbol{x}_{YX} = \boldsymbol{x}$, we get:

$$\boldsymbol{x} \sim N(0, \begin{pmatrix} \lambda_1 + \sigma_0^2 & \dots & 0 \\ \vdots & \ddots & \vdots \\ 0 & \dots & \lambda_m + \sigma_0^2 \end{pmatrix}) \tag{16}$$

Then the log-likelihood function

$$l = -\frac{1}{2}(\frac{x_1^2}{\lambda_1 + \sigma_0^2} + \cdots + \frac{x_m^2}{\lambda_m + \sigma_0^2}) - \frac{1}{2}(\ln(\lambda_1 + \sigma_0^2) + \cdots + \ln(\lambda_m + \sigma_0^2)) + Constant \tag{17}$$

To get the MLE of $\sigma_0^2$, it is equivalent to get root of

$$\frac{x_1^2}{(\lambda_1 + \sigma_0^2)^2} + \cdots + \frac{x_m^2}{(\lambda_m + \sigma_0^2)^2} = \frac{1}{\lambda_1 + \sigma_0^2} + \cdots + \frac{1}{\lambda_1 + \sigma_0^2} \tag{18}$$

We can use the bisection method to search root for (18) and get the root, i.e., MLE of $\sigma_0^2$, denoted as $\hat{\sigma}_0^2$.

Step (3): Update $\mathbf{V}_{YX}$ with $\hat{\sigma}_0^2$, then repeat Step (1) and Step (2) iteratively until convergence, to get the final estimations $(\hat{b}_0, \hat{K}_{YX})$ and $\hat{\sigma}_0^2$. And we can get $se(\hat{b}_0)$ and $se(\hat{K}_{YX})$ from (12).

##### 2.3 CD-GLS: Derivation and Estimation of $(b_0, K_{YX})$ and $\sigma_0^2$

In **CD-Egger** a possible downside is we ignore the variability in $\boldsymbol{r}_{Xg}$ while account for that of $\boldsymbol{r}_{Yg}$. We propose following method **Causal Direction-GLS**, and **CD-GLS** for short, to take into consider of both variation in $\boldsymbol{r}_{Xg}$ and $\boldsymbol{r}_{Yg}$.

Denote $\boldsymbol{b}^*_{Yg} = \boldsymbol{b}_{Yg} - b_0 \cdot \boldsymbol{v}$, so $\boldsymbol{b}^*_{Yg} \sim N(0, \sigma_0^2 \boldsymbol{\Sigma}^2)$. Then $\frac{\boldsymbol{\rho}_{Yg} - b_0 \cdot \boldsymbol{v}}{\boldsymbol{\rho}_{Xg}} = K_{YX} + \frac{\boldsymbol{b}^*_{Yg}}{\boldsymbol{\rho}_{Xg}}$, using Delta Method we will have:

$$\frac{\boldsymbol{r}_{Yg} - b_0 \cdot \boldsymbol{v}}{\boldsymbol{r}_{Xg}}_{|\boldsymbol{\rho}_{Yg}, \boldsymbol{\rho}_{Xg}} \sim N\left(\begin{pmatrix} \frac{\rho_{Yg_1} - b_0 \cdot v_1}{\rho_{Xg_1}} \\ \vdots \\ \frac{\rho_{Yg_m} - b_0 \cdot v_m}{\rho_{Xg_m}} \end{pmatrix}, \mathbf{J}_g^T \cdot \begin{pmatrix} \frac{\mathbf{V}_{Yg}}{n_Y} & \mathbf{0}_{m\times m} \\ \mathbf{0}_{m\times m} & \frac{\mathbf{V}_{Xg}}{n_X} \end{pmatrix} \cdot \mathbf{J}_g \right) := N(\mathbf{1} \cdot K_{YX} + \frac{\boldsymbol{b}^*_{Yg}}{\boldsymbol{\rho}_{Xg}}, \mathbf{V}_{YXg}), \quad (19)$$

Here

$$\mathbf{J}_g = \begin{pmatrix} \frac{1}{r_{Xg_1}} & \cdots & 0 & -\frac{r_{Yg_1} - b_0 \cdot v_1}{r^2_{Xg_1}} & \cdots & 0 \\ \vdots & \ddots & \vdots & \vdots & \ddots & \vdots \\ 0 & \cdots & \frac{1}{r_{Xg_m}} & 0 & \cdots & -\frac{r_{Yg_m} - b_0 \cdot v_m}{r^2_{Xg_m}} \end{pmatrix} \quad (20)$$

So we have:

$$\frac{\boldsymbol{r}_{Yg} - b_0 \cdot \boldsymbol{v}}{\boldsymbol{r}_{Xg}} = \begin{pmatrix} \frac{r_{Yg_1} - b_0 \cdot v_1}{r_{Xg_1}} \\ \vdots \\ \frac{r_{Yg_m} - b_0 \cdot v_m}{r_{Xg_m}} \end{pmatrix} \sim N(\mathbf{1} \cdot K_{YX}, \mathbf{V}_{YXg} + \sigma_0^2 \mathbf{P}) = N(\mathbf{1} \cdot K_{YX}, \mathbf{V}_{YX}) \quad (21)$$

Here

$$\mathbf{P} = \begin{pmatrix} \frac{1}{r_{Xg_1}} & & \\ & \ddots & \\ & & \frac{1}{r_{Xg_m}} \end{pmatrix} \boldsymbol{\Sigma}^2 \begin{pmatrix} \frac{1}{r_{Xg_1}} & & \\ & \ddots & \\ & & \frac{1}{r_{Xg_m}} \end{pmatrix} \quad (22)$$

Again, similar to CD-Egger, we can use iterative method to estimate $(b_0, K_{YX})$ and $\sigma_0^2$, and get standard errors of the estimations $se(\hat{b}_0), se(\hat{K}_{YX})$. Equation (21) is equivalent to

$$\frac{\boldsymbol{r}_{Yg}}{\boldsymbol{r}_{Xg}} = b_0 \cdot \frac{\boldsymbol{v}}{\boldsymbol{r}_{Xg}} + K_{YX} \cdot \mathbf{1} + \boldsymbol{\varepsilon}, \boldsymbol{\varepsilon} \sim N(0, \mathbf{V}_{YXg} + \sigma_0^2 \mathbf{P}) \quad (23)$$

Set initial $b_0 = 0$, $\sigma_0^2 = 0$.

Step (1): Given $b_0, \sigma_0^2$, we can get $\mathbf{V}_{YX} = \mathbf{V}_{YXg} + \sigma_0^2 \mathbf{P}$. Then estimating $(b_0, K_{YX})$ is a Generalize Least Square problem with design matrix $\mathbf{X}$ and covariance matrix of errors $\boldsymbol{\Omega}$:

$$\mathbf{X} = \begin{pmatrix} \frac{\boldsymbol{v}}{\boldsymbol{r}_{Xg}} & \mathbf{1} \end{pmatrix} = \begin{pmatrix} \frac{v_1}{r_{Xg_1}} & 1 \\ \vdots & \vdots \\ \frac{v_m}{r_{Xg_m}} & 1 \end{pmatrix}$$
$$\boldsymbol{\Omega} = \mathbf{V}_{YX} \quad (24)$$

So we can get the estimation $(\hat{b}_0, \hat{K}_{YX})$ and their covariance matrix:

$$\begin{pmatrix} \hat{b}_0 \\ \hat{K}_{YX} \end{pmatrix} = (\mathbf{X}^T \boldsymbol{\Omega}^{-1} \mathbf{X})^{-1} \mathbf{X}^T \boldsymbol{\Omega}^{-1} \frac{\boldsymbol{r}_{Yg}}{\boldsymbol{r}_{Xg}}$$
$$cov \begin{pmatrix} \hat{b}_0 \\ \hat{K}_{YX} \end{pmatrix} = (\mathbf{X}^T \boldsymbol{\Omega}^{-1} \mathbf{X})^{-1} \quad (25)$$

Step (2): Given $(\hat{b}_0, \hat{K}_{YX})$, plug them in equation (23), denoting

$$\boldsymbol{x}_{YX} = \frac{\boldsymbol{r}_{Yg}}{\boldsymbol{r}_{Xg}} - \hat{b}_0 \cdot \frac{\boldsymbol{v}}{\boldsymbol{r}_{Xg}} - \hat{K}_{YX} \cdot \mathbf{1}$$
$$\mathbf{U} = \begin{pmatrix} r_{Xg_1} & & \\ & \ddots & \\ & & r_{Xg_m} \end{pmatrix} \tag{26}$$

We get

$$\mathbf{\Sigma}^{-1}\mathbf{U}\boldsymbol{x}_{YX} \sim N(0, \mathbf{\Sigma}^{-1}\mathbf{U}\mathbf{V}_{YXg}\mathbf{U}\mathbf{\Sigma}^{-1} + \sigma_0^2\mathbf{I}) \tag{27}$$

Do the eigenvalue decomposition of

$$\mathbf{\Sigma}^{-1}\mathbf{U}\mathbf{V}_{YXg}\mathbf{U}\mathbf{\Sigma}^{-1} = \mathbf{Q}^T \begin{pmatrix} \lambda_1 & \dots & 0 \\ \vdots & \ddots & \vdots \\ 0 & \dots & \lambda_m \end{pmatrix} \mathbf{Q} \tag{28}$$

Here $\mathbf{Q}$ is orthonormal, $\lambda_1, \cdots, \lambda_m$ are eigen-values. Denote $\mathbf{Q}\mathbf{\Sigma}^{-1}\mathbf{U}\boldsymbol{x}_{YX} = \boldsymbol{x}$, we get:

$$\boldsymbol{x} \sim N(0, \begin{pmatrix} \lambda_1 + \sigma_0^2 & \dots & 0 \\ \vdots & \ddots & \vdots \\ 0 & \dots & \lambda_m + \sigma_0^2 \end{pmatrix}) \tag{29}$$

Then the log-likelihood function

$$l = -\frac{1}{2}(\frac{x_1^2}{\lambda_1 + \sigma_0^2} + \cdots + \frac{x_m^2}{\lambda_m + \sigma_0^2}) - \frac{1}{2}(\ln(\lambda_1 + \sigma_0^2) + \cdots + \ln(\lambda_m + \sigma_0^2)) + Constant \tag{30}$$

To get the MLE of $\sigma_0^2$, it is equivalent to get root of

$$\frac{x_1^2}{(\lambda_1 + \sigma_0^2)^2} + \cdots + \frac{x_m^2}{(\lambda_m + \sigma_0^2)^2} = \frac{1}{\lambda_1 + \sigma_0^2} + \cdots + \frac{1}{\lambda_1 + \sigma_0^2} \tag{31}$$

We can use the bisection method to search root for (31) and get the root, i.e., MLE of $\sigma_0^2$, denoted as $\hat{\sigma}_0^2$.

Step (3): Update $\mathbf{V}_{YX}$ with $\hat{b}_0$ and $\hat{\sigma}_0^2$, then repeat Step (1) and Step (2) iteratively until convergence, to get the final estimations $(\hat{b}_0, \hat{K}_{YX})$ and $\hat{\sigma}_0^2$. And we can get $se(\hat{b}_0)$ and $se(\hat{K}_{YX})$ from (25).

### 3 Degrees of Freedom for the Goodness-of-Fit Tests

#### 3.1 CD-Ratio

For **CD-Ratio**, from the main text we have:

$$\frac{\boldsymbol{r}_{Yg}}{\boldsymbol{r}_{Xg}} \sim N(\mathbf{1} \cdot K_{YX}, \mathbf{V}_{YXg}) \tag{32}$$

And

$$\hat{K}_{YX} = \frac{\mathbf{1}^T \cdot \mathbf{V}_{YXg}^{-1} \cdot \frac{\mathbf{r}_{Yg}}{\mathbf{r}_{Xg}}}{\mathbf{1}^T \cdot \mathbf{V}_{YXg}^{-1} \cdot \mathbf{1}} \tag{33}$$

For simplicity, we use $\boldsymbol{r}$ to represent $\frac{\boldsymbol{r}_{Yg}}{\boldsymbol{r}_{Xg}}$, $K$ to represent $K_{YX}$, $\hat{K}$ to represent $\hat{K}_{YX}$, and $\mathbf{V}$ to represent $\mathbf{V}_{YXg}$. Then the test statistic $Q_{Ratio}$ is:

$$Q_{Ratio} = \left(\boldsymbol{r} - \mathbf{1}\cdot\hat{K}\right)^T \mathbf{V}^{-1}\left(\boldsymbol{r} - \mathbf{1}\cdot\hat{K}\right) \tag{34}$$

And we have

$$\boldsymbol{r} - \mathbf{1}\cdot\hat{K} = \left(\mathbf{I} - \frac{\mathbf{1}\mathbf{1}^T\mathbf{V}^{-1}}{\mathbf{1}^T\mathbf{V}^{-1}\mathbf{1}}\right)\boldsymbol{r} \tag{35}$$

So

$$\mathbf{V}^{-\frac{1}{2}}(\boldsymbol{r} - \mathbf{1}\cdot\hat{K}) = \mathbf{V}^{-\frac{1}{2}}\left(\mathbf{I} - \frac{\mathbf{1}\mathbf{1}^T\mathbf{V}^{-1}}{\mathbf{1}^T\mathbf{V}^{-1}\mathbf{1}}\right)\boldsymbol{r} \sim N(\boldsymbol{\mu}, \mathbf{A}) \tag{36}$$

Here

$$\boldsymbol{\mu} = \mathbf{V}^{-\frac{1}{2}}\left(\mathbf{I} - \frac{\mathbf{1}\mathbf{1}^T\mathbf{V}^{-1}}{\mathbf{1}^T\mathbf{V}^{-1}\mathbf{1}}\right)\mathbf{1}\cdot K = \mathbf{0} \tag{37}$$

And

$$\begin{aligned}\mathbf{A} &= \mathbf{V}^{-\frac{1}{2}}\left(\mathbf{I} - \frac{\mathbf{1}\mathbf{1}^T\mathbf{V}^{-1}}{\mathbf{1}^T\mathbf{V}^{-1}\mathbf{1}}\right)\mathbf{V}\left(\mathbf{I} - \frac{\mathbf{V}^{-1}\mathbf{1}\mathbf{1}^T}{\mathbf{1}^T\mathbf{V}^{-1}\mathbf{1}}\right)\mathbf{V}^{-\frac{1}{2}} \\ &= \mathbf{I} - \frac{\mathbf{V}^{-\frac{1}{2}}\mathbf{1}\mathbf{1}^T\mathbf{V}^{-\frac{1}{2}}}{\mathbf{1}^T\mathbf{V}^{-1}\mathbf{1}}\end{aligned} \tag{38}$$

Since $\frac{\mathbf{V}^{-\frac{1}{2}}\mathbf{1}\mathbf{1}^T\mathbf{V}^{-\frac{1}{2}}}{\mathbf{1}^T\mathbf{V}^{-1}\mathbf{1}}$ is the projection matrix to vector $\mathbf{V}^{-\frac{1}{2}}\mathbf{1}$, eigenvalues of $\mathbf{A}$ are $(m-1)$ 1's and one 0, leading to $Q_{Ratio} \sim \chi^2_{m-1}$.

#### 3.2 CD-Egger

For **CD-Egger**, from the main text we have:

$$\boldsymbol{r}_{Yg} = b_0 \cdot \boldsymbol{v} + K_{YX}\cdot \boldsymbol{r}_{Xg} + \boldsymbol{\varepsilon}, \boldsymbol{\varepsilon} \sim N(0, \frac{\mathbf{V}_{Yg}}{n_Y} + \sigma_0^2\boldsymbol{\Sigma}^2 = \mathbf{V}_{YX}) \tag{39}$$

As we show in the Section 3 above, plug in estimation $\hat{\sigma}_0^2$ of $\sigma_0^2$ we get $\mathbf{V}_{YX}$, with notations in equation (11), we have

$$\begin{pmatrix}\hat{b}_0 \\ \hat{K}_{YX}\end{pmatrix} = (\mathbf{X}^T\boldsymbol{\Omega}^{-1}\mathbf{X})^{-1}\mathbf{X}^T\boldsymbol{\Omega}^{-1}\boldsymbol{r}_{Yg} \tag{40}$$

And the test statistic $Q_{Egger}$ is:

$$Q_{Egger} = (\boldsymbol{r}_{Yg} - \hat{b}_0\cdot\boldsymbol{v} - \hat{K}_{YX}\cdot\boldsymbol{r}_{Xg})^T\mathbf{V}_{YX}^{-1}(\boldsymbol{r}_{Yg} - \hat{b}_0\cdot\boldsymbol{v} - \hat{K}_{YX}\cdot\boldsymbol{r}_{Xg}) \tag{41}$$

Here

$$\begin{aligned}&\boldsymbol{r}_{Yg} - \hat{b}_0\cdot\boldsymbol{v} - \hat{K}_{YX}\cdot\boldsymbol{r}_{Xg} \\ &= \left(\mathbf{I} - \mathbf{X}(\mathbf{X}^T\boldsymbol{\Omega}^{-1}\mathbf{X})^{-1}\mathbf{X}^T\boldsymbol{\Omega}^{-1}\right)\boldsymbol{r}_{Yg} \\ &= \left(\mathbf{I} - \mathbf{X}(\mathbf{X}^T\boldsymbol{\Omega}^{-1}\mathbf{X})^{-1}\mathbf{X}^T\boldsymbol{\Omega}^{-1}\right)\boldsymbol{\varepsilon}\end{aligned} \tag{42}$$

So

$$\begin{aligned}&\mathbf{V}_{YX}^{-\frac{1}{2}}(\boldsymbol{r}_{Yg} - \hat{b}_0\cdot\boldsymbol{v} - \hat{K}_{YX}\cdot\boldsymbol{r}_{Xg}) \\ =&\mathbf{V}_{YX}^{-\frac{1}{2}}\left(\mathbf{I} - \mathbf{X}(\mathbf{X}^T\boldsymbol{\Omega}^{-1}\mathbf{X})^{-1}\mathbf{X}^T\boldsymbol{\Omega}^{-1}\right)\boldsymbol{\varepsilon} \sim N(\mathbf{0}, \mathbf{A})\end{aligned} \tag{43}$$

Here

$$\begin{aligned}\mathbf{A} &= \mathbf{V}_{YX}^{-\frac{1}{2}}\left(\mathbf{I}-\mathbf{X}(\mathbf{X}^T\boldsymbol{\Omega}^{-1}\mathbf{X})^{-1}\mathbf{X}^T\boldsymbol{\Omega}^{-1}\right)\mathbf{V}_{YX}\left(\mathbf{I}-\boldsymbol{\Omega}^{-1}\mathbf{X}(\mathbf{X}^T\boldsymbol{\Omega}^{-1}\mathbf{X})^{-1}\mathbf{X}^T\right)\mathbf{V}_{YX}^{-\frac{1}{2}} \\ &= \mathbf{I}-\boldsymbol{\Omega}^{-\frac{1}{2}}\mathbf{X}(\mathbf{X}^T\boldsymbol{\Omega}^{-1}\mathbf{X})^{-1}\mathbf{X}^T\boldsymbol{\Omega}^{-\frac{1}{2}}\end{aligned} \tag{44}$$

Since $\boldsymbol{\Omega}^{-\frac{1}{2}}\mathbf{X}(\mathbf{X}^T\boldsymbol{\Omega}^{-1}\mathbf{X})^{-1}\mathbf{X}^T\boldsymbol{\Omega}^{-\frac{1}{2}}$ is the projection matrix to dimension-2 space $\boldsymbol{\Omega}^{-\frac{1}{2}}\mathbf{X}$, eigenvalues of $\mathbf{A}$ are $(m-2)$ 2's and two 0's, leading to $Q_{Egger} \sim \chi^2_{m-2}$.

##### 3.3 CD-GLS

Based on the asymptotic distribution from above equation (21), for CD-GLS we can construct GOF test:

$$Q_{GLS} = \left(\frac{\boldsymbol{r}_{Yg}-\hat{b}_0\cdot\boldsymbol{v}}{\boldsymbol{r}_{Xg}} - \mathbf{1}\cdot\hat{K}_{YX}\right)^T \mathbf{V}_{YX}^{-1}\left(\frac{\boldsymbol{r}_{Yg}-\hat{b}_0\cdot\boldsymbol{v}}{\boldsymbol{r}_{Xg}} - \mathbf{1}\cdot\hat{K}_{YX}\right) \tag{45}$$

Similar to CD-Egger, the (asymptotic) null distribution is $Q_{GLS} \sim \chi^2_{m-2}$.

#### 4 More Results for LDL and CAD

##### 4.1 Removing Locus 253 and Locus 1246

For the real data example of LDL and CAD, we showed the results using 22 SNPs from 12 loci in the main text, Locus 253 and Locus 1246 showed negative effects of LDL on CAD. We can remove these 2 loci, then apply CD-Ratio, CD-Egger and CD-GLS to the rest 20 SNPs from the 10 loci. Figure S1 shows the corresponding results as forest plots, and Table S1 shows 95% confidence intervals for the combined results.

Figure S1: Forest plots for inferring the causal direction between LDL and CAD, showing SNP and locus-specific results and the combined results across 20 SNPs from 10 loci, after removing Locus 253 and Locus 1246.

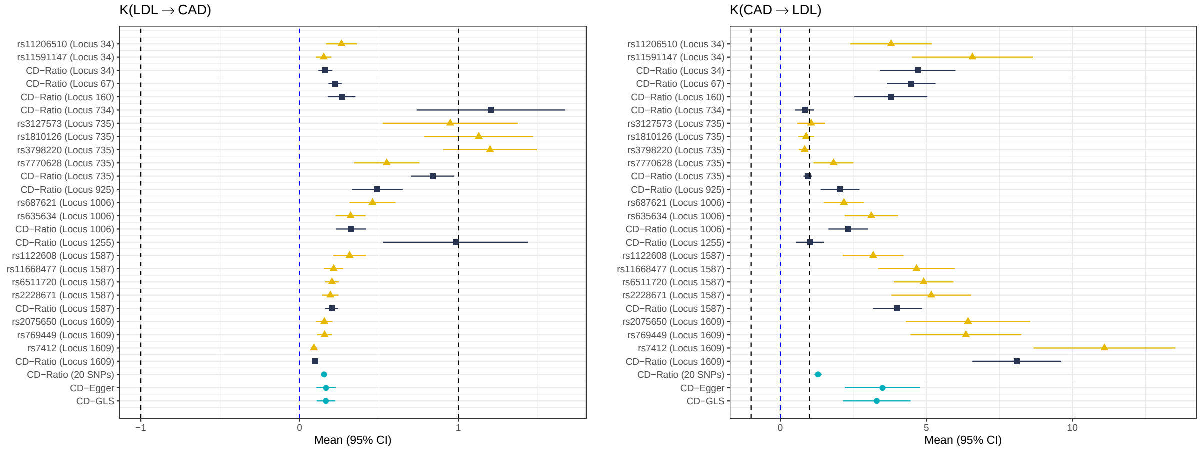

Table S1: Results of inferring the causal direction between LDL and CAD after combining across 20 SNPs in 10 loci after removing Locus 253 and Locus 1246.

| Method | *LDL → CAD* | | *CAD → LDL* | |
|---|---|---|---|---|
| | $\hat{K}$ [ 95% CI] | $\hat{b}_0$ (SE) | $\hat{K}$ [95% CI] | $\hat{b}_0$ (SE) |
| CD-Ratio | 0.154 [0.140, 0.168] | NA | 1.293 [1.168, 1.417] | NA |
| CD-Egger | 0.167 [0.106, 0.228] | 0.005 (0.002) | 3.500 [2.209, 4.790] | -0.021 (0.010) |
| CD-GLS | 0.166 [0.107, 0.225] | 0.005 (0.002) | 3.301 [2.144, 4.457] | -0.019 (0.009) |

#### 4.2 Only Using Independent SNPs

For the real data example of LDL and CAD, we showed the results using 22 SNPs from 12 loci in the main text. We can only pick the most significant SNP at each of 12 loci according to the combined significance level with LDL and CAD, as described in Section 2.7 in the main text, then we get 12 independent SNPs from the 12 loci. We can apply CD-Ratio to each SNPs and all 12 SNPs, and apply CD-Egger and CD-GLS to all 12 SNPs. Figure S2 shows the corresponding results as forest plots, and Table S2 shows 95% confidence intervals for the combined results.

Figure S2: Forest plots for inferring the causal direction between LDL and CAD, showing SNP results and the combined results across 12 independent SNPs from 12 loci

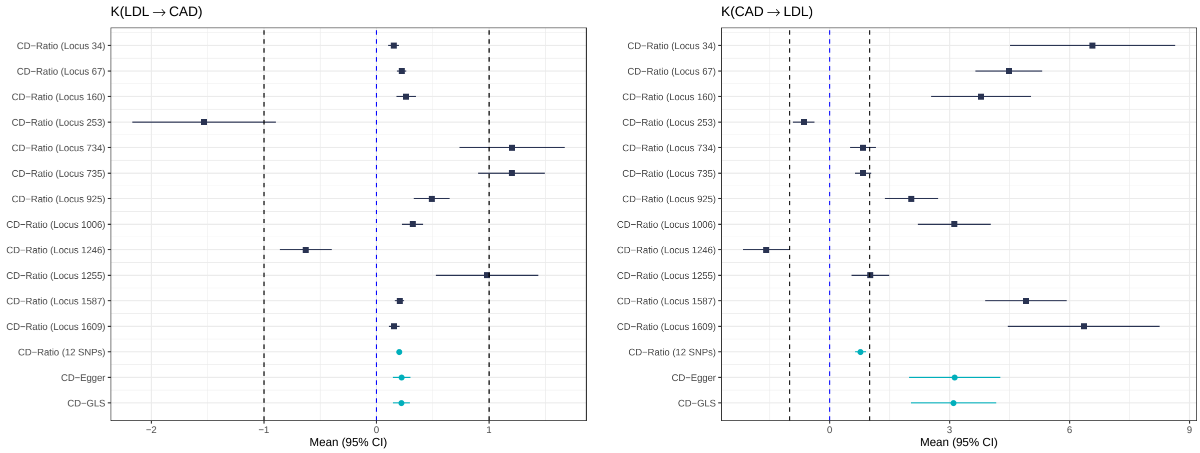


Table S2: Results of inferring the causal direction between LDL and CAD after combining across 12 independent SNPs in 12 loci.

| Method | *LDL → CAD* | | *CAD → LDL* | |
|---|---|---|---|---|
| | $\hat{K}$ [ 95% CI] | $\hat{b}_0$ (SE) | $\hat{K}$ [95% CI] | $\hat{b}_0$ (SE) |
| CD-Ratio | 0.203 [0.183, 0.223] | NA | 0.769 [0.633, 0.904] | NA |
| CD-Egger | 0.223 [0.145, 0.301] | 0.007 (0.002) | 3.126 [1.983, 4.268] | -0.023 (0.010) |
| CD-GLS | 0.222 [0.147, 0.297] | 0.007 (0.002) | 3.097 [2.029, 4.165] | -0.025 (0.009) |

### 5 Results for LDL/HDL and CAD with Multivariable Approaches

#### 5.1 LDL and CAD: MV-CD-Egger (adjusting for HDL)

For LDL and CAD, we started with the 22 SNPs from 12 independent loci as described in the main text. Among the $p$-values of these 22 SNPs for HDL, 8 of them were smaller than $5 \times 10^{-8}$, 2 of them between $5 \times 10^{-8}$ and $5 \times 10^{-6}$, 3 of them between $5 \times 10^{-6}$ and $5 \times 10^{-4}$, and 9 of them larger than $5 \times 10^{-4}$. We could use either all these 22 SNPs or only the 8 SNPs significant for all 3 traits. Table S3 and Table S4 show the results (after adjusting for HDL) with the two sets of the SNPs as IVs respectively.

Table S3: The results for inferring the causal direction between LDL and CAD combining across all 22 SNPs in 12 loci, and adjusting for HDL.

| | $LDL \to CAD$ | | $CAD \to LDL$ | |
|---|---|---|---|---|
| Method | $\hat{K}$ [ 95% CI] | $\hat{b}_0$ (SE) | $\hat{K}$ [95% CI] | $\hat{b}_0$ (SE) |
| CD-Egger | 0.172 $[0.078, 0.267]$ | 0.006 (0.002) | 2.158 $[0.996, 3.319]$ | -0.020 (0.008) |

Table S4: The results for inferring the causal direction between LDL and CAD combining across 8 SNPs from 6 loci, and adjusting for HDL.

| | $LDL \to CAD$ | | $CAD \to LDL$ | |
|---|---|---|---|---|
| Method | $\hat{K}$ [ 95% CI] | $\hat{b}_0$ (SE) | $\hat{K}$ [95% CI] | $\hat{b}_0$ (SE) |
| CD-Egger | 0.114 $[-0.008, 0.235]$ | 0.003 (0.004) | 2.680 $[-0.099, 5.458]$ | -0.033 (0.014) |

We can see that, using the 22 SNPs, the 95% CI of $K$ for $LDL \to CAD$ was [0.078,0.267], completely inside (0,1], while the 95% for $CAD \to LDL$ was [0.996,3.319], almost completely outside [-1,1]. So we have strong evidence to say that, after adjusting for HDL, we conclude that the causal direction is from LDL to CAD. Using the 8 SNPs, although the lengths of the confidence intervals were larger so we could not make a conclusion, the point estimate of $K$ for $LDL \to CAD$ was 0.114, inside (0,1] while that for $CAD \to LDL$ was 2.680, larger than 1 giving some evidence to support the causal effect of LDL on CAD.

#### 5.2 HDL and CAD: MV-CD-Egger (after adjusting for LDL)

For HDL and CAD, we selected the 8 SNPs from 6 independent loci as described in the main text, and all these 8 SNPs had $p$-values less than $5 \times 10^{-8}$ for LDL. Table S5 shows the results (after adjusting for LDL).

Table S5: The results for inferring the causal direction between HDL and CAD combining across all 8 SNPs in 6 loci, and adjusting for LDL.

| | $HDL \to CAD$ | | $CAD \to HDL$ | |
|---|---|---|---|---|
| Method | $\hat{K}$ [ 95% CI] | $\hat{b}_0$ (SE) | $\hat{K}$ [95% CI] | $\hat{b}_0$ (SE) |
| CD-Egger | -0.181 $[-0.531, 0.168]$ | 0.003 (0.004) | -0.660 $[-1.903, 0.584]$ | -0.007 (0.007) |

We can see that, for both directions the 95% CIs covered 0, and both point estimates were in [-1,0), so we cannot conclude with any causal relationship between HDL and CAD.

#### 5.3 MV-MR Results

Using R package **TwoSampleMR**, we applied two multivariable MR methods: the first one was MV-IVW with function `mv_ivw()` and the second MV-MR-Egger with `mv_multiple()` with an intercept term. Table S6 shows the results for CAD, LDL and HDL with each as the outcome respectively. As before, we used 2017 lipid data and 2017 CAD data. We set the significance cutoff at $5 \times 10^{-8}$ to choose independent SNPs as instruments. When CAD was the outcome we used 169 IVs: 89 significant for LDL, 106 significant for HDL, and 26 for both; when LDL was the outcome, we use 142 IVs: 34 significant for CAD, 114 significant for HDL, and 6 for both; when HDL was the outcome, we used 127 IVs: 38 significant for CAD, 101 significant for LDL, and 12 for both.

Table S6: Multivariable MR results with one of CAD, LDL and HDL as the outcome and the other two as exposure based on 2017 lipids GWAS summary data and 2017 CAD GWAS summary data.

| CAD as outcome | | | | | | |
|---|---|---|---|---|---|---|
| | MV-MR-IVW | | | MV-MR-Egger | | |
| Covariate | $\hat{\beta}$ | SE($\hat{\beta}$) | $p$-value | $\hat{\beta}$ | SE($\hat{\beta}$) | $p$-value |
| LDL | 0.492 | 0.053 | $1.1 \times 10^{-20}$ | 0.485 | 0.047 | $7.2 \times 10^{-25}$ |
| HDL | -0.219 | 0.045 | $1.2 \times 10^{-6}$ | -0.214 | 0.044 | $1.2 \times 1.2^{-6}$ |
| **LDL as outcome** | | | | | | |
| | MV-MR-IVW | | | MV-MR-Egger | | |
| Covariate | $\hat{\beta}$ | SE($\hat{\beta}$) | $p$-value | $\hat{\beta}$ | SE($\hat{\beta}$) | $p$-value |
| CAD | 0.187 | 0.148 | 0.206 | 0.402 | 0.077 | $1.7 \times 10^{-7}$ |
| HDL | 0.134 | 0.077 | 0.083 | 0.034 | 0.080 | 0.673 |
| **HDL as outcome** | | | | | | |
| | MV-MR-IVW | | | MV-MR-Egger | | |
| Covariate | $\hat{\beta}$ | SE($\hat{\beta}$) | $p$-value | $\hat{\beta}$ | SE($\hat{\beta}$) | $p$-value |
| CAD | -0.044 | 0.033 | 0.177 | -0.125 | 0.074 | 0.092 |
| LDL | 0.029 | 0.085 | 0.736 | -0.034 | 0.066 | 0.614 |

Through R package **TwoSampleMR**, the functions can automatically extracts IVs from 2013 lipid data and 2015 CAD data from its database, we show the corresponding results in Table S7. We set cutoff $5 \times 10^{-8}$ to choose independent SNPs as instruments, when CAD is outcome we use 132 IVs, 68 significant for LDL, 75 significant for HDL, and 11 for both; when LDL is outcome we use 102 IVs, 26 significant for CAD, 79 significant for HDL, and 3 for both; when HDL is outcome we use 95 IVs, 26 significant for CAD, 74 significant for LDL, and 5 for both.

Table S7: Multivariable MR results with one of CAD, LDL, HDL as outcome and the other two as covariates, using 2013 lipids GWAS summary data and 2015 CAD GWAS summary data.

| CAD as outcome | | | | | | |
|---|---|---|---|---|---|---|
| | MV-MR-IVW | | | MV-MR-Egger | | |
| Covariate | $\hat{\beta}$ | SE($\hat{\beta}$) | $p$-value | $\hat{\beta}$ | SE($\hat{\beta}$) | $p$-value |
| LDL | 0.397 | 0.056 | $1.3 \times 10^{-12}$ | 0.389 | 0.050 | $8.4 \times 10^{-15}$ |
| HDL | -0.142 | 0.054 | $8.1 \times 10^{-3}$ | -0.141 | 0.057 | 0.014 |
| **LDL as outcome** | | | | | | |
| | MV-MR-IVW | | | MV-MR-Egger | | |
| Covariate | $\hat{\beta}$ | SE($\hat{\beta}$) | $p$-value | $\hat{\beta}$ | SE($\hat{\beta}$) | $p$-value |
| CAD | 0.067 | 0.070 | 0.336 | 0.144 | 0.057 | 0.011 |
| HDL | 0.017 | 0.066 | 0.797 | -0.045 | 0.063 | 0.479 |
| **HDL as outcome** | | | | | | |
| | MV-MR-IVW | | | MV-MR-Egger | | |
| Covariate | $\hat{\beta}$ | SE($\hat{\beta}$) | $p$-value | $\hat{\beta}$ | SE($\hat{\beta}$) | $p$-value |
| CAD | -0.018 | 0.028 | 0.513 | -0.054 | 0.073 | 0.459 |
| LDL | -0.039 | 0.102 | 0.700 | -0.080 | 0.071 | 0.258 |

Results in Table S6 and Table S7 are similar. We can see at significant level 0.05, both MV-MR methods would conclude with LDL and HDL has causal effect on CAD, and MV-MR-Egger concludes CAD has causal effect on LDL.

### 6 More Simulation Results

#### 6.1 Details of Simulation Setups

We used three parameters $(\beta_{YX}, \mu_\alpha, \sigma_\alpha)$ in each simulation setup. $\beta_{YX}$ controls the causal effect size, $\mu_\alpha$ controls the mean direct effect size, $\sigma_\alpha$ controls the variation of the (random) direct effect sizes. We tried 75 different combinations of $\beta_{YX} = -0.2, -0.1, 0, 0.1, 0.2, \mu_\alpha = 0, 0.2, 0.4, 0.6, 1, \sigma_\alpha = 0, 0.1, 0.2$. For each setup we first generated $\boldsymbol{\beta}_{Xg} = (\beta_{Xg_1}, \beta_{Xg_2}, ..., \beta_{Xg_{22}})^T$, the vector of the effect sizes of the 22 SNPs on $X$, with each element independently drawn from a truncated standard normal distribution (truncated at 0.5 to ensure $|\beta_{Xg_j}| \geq 0.5$). Then in each simulation, we generated the two traits from the true models:

$$X_i = \boldsymbol{Z}_i \boldsymbol{\beta}_{Xg} + U_i + \varepsilon_{Xi}, \qquad Y_i = \beta_{YX} \cdot X_i + \boldsymbol{Z}_i \boldsymbol{\alpha} + U_i + \varepsilon_{Yi},$$

where $\boldsymbol{Z}_i$ was the (row) vector of the genotype scores of the 22 SNPs for subject $i = 1, 2, ..., n$, $U_i \sim N(0,1)$ was a confounder; the direct effects of the SNPs on $Y$, $\boldsymbol{\alpha} = (\alpha_1, ..., \alpha_{22})^T$, were generated independently $\alpha_i \sim N(\mu_\alpha, \sigma_\alpha^2)$; and both error terms $\varepsilon_{Xi}, \varepsilon_{Yi}$ were independently from $N(0,4)$.

#### 6.2 Estimating $K_{YX}$

Table S8 shows simulation results for estimating $K_{YX}$ of CD-Ratio, CD-Egger, and CD-GLS, for the setup $\mu_\alpha = 0$, $\sigma_\alpha = 0$, and various $\beta_{YX}$ levels. In each row we list the value $\beta_{YX}$ in the first column, and corresponding true value $K_{YX}$ in the second column, and for each of three methods we show the estimation results similar to Table 5 in the main text. Table S9 to Table S22 show similar results for other setups. When $\beta_{YX} \neq 0$ the true causal direction is from $X$ to $Y$, we can calculate the true value of

$K_{YX} = \beta_{YX} \cdot \frac{\sqrt{var(X)}}{\sqrt{var(Y)}}$, here

$$var(X) = \boldsymbol{\beta}_{Xg}^T \boldsymbol{\Sigma} \boldsymbol{\beta}_{Xg} + var(U) + var(\varepsilon_X)$$
$$var(Y) = (\mu_\alpha \cdot \mathbf{1} + \beta_{YX} \cdot \boldsymbol{\beta}_{Xg})^T \boldsymbol{\Sigma} (\mu_\alpha \cdot \mathbf{1} + \beta_{YX} \cdot \boldsymbol{\beta}_{Xg}) + (\beta_{YX}+1)^2 var(U) + \beta_{YX}^2 var(\varepsilon_X) + var(\varepsilon_Y) \tag{46}$$

$\boldsymbol{\Sigma}$ is the covariance matrix of SNPs. When $\beta_{YX} = 0$, we have $K_{YX} = 0$.

Table S8: Results of $\mu_\alpha = 0$, $\sigma_\alpha = 0$: $K_{YX}$

| | | CD-Ratio | | | CD-Egger | | | CD-GLS | | |
|---|---|---|---|---|---|---|---|---|---|---|
| $\beta_{YX}$ | $K_{YX}$ | Mean($\hat{K}$) | sd($\hat{K}$) | Mean$(se(\hat{K}))$ | Mean($\hat{K}$) | sd($\hat{K}$) | Mean$(se(\hat{K}))$ | Mean($\hat{K}$) | sd($\hat{K}$) | Mean$(se(\hat{K}))$ |
| -0.2 | -0.442 | -0.4415 | 0.0028 | 0.0036 | -0.4415 | 0.0028 | 0.0035 | -0.4415 | 0.0028 | 0.0036 |
| -0.1 | -0.2344 | -0.2341 | 0.0032 | 0.0036 | -0.2341 | 0.0032 | 0.0036 | -0.2341 | 0.0032 | 0.0036 |
| 0 | 0 | 0.0001 | 0.0034 | 0.0035 | 0.0001 | 0.0034 | 0.0036 | 0.0001 | 0.0034 | 0.0036 |
| 0.1 | 0.2257 | 0.2256 | 0.0033 | 0.0036 | 0.2256 | 0.0033 | 0.0036 | 0.2256 | 0.0033 | 0.0036 |
| 0.2 | 0.4139 | 0.4137 | 0.003 | 0.0036 | 0.4137 | 0.003 | 0.0035 | 0.4137 | 0.003 | 0.0036 |

Table S9: Results of $\mu_\alpha = 0.2$, $\sigma_\alpha = 0$: $K_{YX}$

| | | CD-Ratio | | | CD-Egger | | | CD-GLS | | |
|---|---|---|---|---|---|---|---|---|---|---|
| $\beta_{YX}$ | $K_{YX}$ | Mean($\hat{K}$) | sd($\hat{K}$) | Mean$(se(\hat{K}))$ | Mean($\hat{K}$) | sd($\hat{K}$) | Mean$(se(\hat{K}))$ | Mean($\hat{K}$) | sd($\hat{K}$) | Mean$(se(\hat{K}))$ |
| -0.2 | -0.3995 | -0.4106 | 0.003 | 0.0038 | -0.3992 | 0.0026 | 0.0034 | -0.3992 | 0.0026 | 0.0035 |
| -0.1 | -0.2093 | -0.2191 | 0.0032 | 0.0039 | -0.2091 | 0.0029 | 0.0035 | -0.2091 | 0.0029 | 0.0035 |
| 0 | 0 | -0.0097 | 0.0034 | 0.0039 | 0 | 0.0031 | 0.0035 | 0 | 0.0031 | 0.0035 |
| 0.1 | 0.2027 | 0.1947 | 0.0034 | 0.0039 | 0.2026 | 0.003 | 0.0035 | 0.2026 | 0.003 | 0.0036 |
| 0.2 | 0.3773 | 0.3675 | 0.0031 | 0.0039 | 0.3771 | 0.0028 | 0.0035 | 0.3771 | 0.0028 | 0.0036 |

Table S10: Results of $\mu_\alpha = 0.4$, $\sigma_\alpha = 0$: $K_{YX}$

| | | CD-Ratio | | | CD-Egger | | | CD-GLS | | |
|---|---|---|---|---|---|---|---|---|---|---|
| $\beta_{YX}$ | $K_{YX}$ | Mean($\hat{K}$) | sd($\hat{K}$) | Mean$(se(\hat{K}))$ | Mean($\hat{K}$) | sd($\hat{K}$) | Mean$(se(\hat{K}))$ | Mean($\hat{K}$) | sd($\hat{K}$) | Mean$(se(\hat{K}))$ |
| -0.2 | -0.3203 | -0.3298 | 0.0031 | 0.004 | -0.3202 | 0.0022 | 0.0032 | -0.3202 | 0.0022 | 0.0033 |
| -0.1 | -0.1649 | -0.1777 | 0.0032 | 0.0041 | -0.1648 | 0.0023 | 0.0033 | -0.1648 | 0.0023 | 0.0033 |
| 0 | 0 | -0.0158 | 0.0033 | 0.0041 | 0 | 0.0025 | 0.0033 | 0 | 0.0025 | 0.0034 |
| 0.1 | 0.1614 | 0.1464 | 0.0033 | 0.0041 | 0.1613 | 0.0025 | 0.0034 | 0.1613 | 0.0025 | 0.0034 |
| 0.2 | 0.3079 | 0.2956 | 0.0032 | 0.0041 | 0.3078 | 0.0024 | 0.0034 | 0.3078 | 0.0024 | 0.0035 |

Table S11: Results of $\mu_\alpha = 0.6$, $\sigma_\alpha = 0$: $K_{YX}$

| | | CD-Ratio | | | CD-Egger | | | CD-GLS | | |
|---|---|---|---|---|---|---|---|---|---|---|
| $\beta_{YX}$ | $K_{YX}$ | Mean($\hat{K}$) | sd($\hat{K}$) | Mean$(se(\hat{K}))$ | Mean($\hat{K}$) | sd($\hat{K}$) | Mean$(se(\hat{K}))$ | Mean($\hat{K}$) | sd($\hat{K}$) | Mean$(se(\hat{K}))$ |
| -0.2 | -0.2535 | -0.2713 | 0.0033 | 0.0041 | -0.2535 | 0.0018 | 0.0031 | -0.2535 | 0.0018 | 0.0032 |
| -0.1 | -0.129 | -0.1538 | 0.0034 | 0.0042 | -0.129 | 0.0019 | 0.0031 | -0.129 | 0.0019 | 0.0032 |
| 0 | 0 | -0.0304 | 0.0034 | 0.0042 | 0 | 0.002 | 0.0032 | 0 | 0.002 | 0.0032 |
| 0.1 | 0.1272 | 0.0955 | 0.0034 | 0.0042 | 0.1272 | 0.002 | 0.0033 | 0.1272 | 0.002 | 0.0033 |
| 0.2 | 0.2469 | 0.2181 | 0.0033 | 0.0042 | 0.2468 | 0.002 | 0.0034 | 0.2468 | 0.002 | 0.0034 |

Table S12: Results of $\mu_\alpha = 1$, $\sigma_\alpha = 0$: $K_{YX}$

| | | CD-Ratio | | | CD-Egger | | | CD-GLS | | |
|---|---|---|---|---|---|---|---|---|---|---|
| $\beta_{YX}$ | $K_{YX}$ | Mean($\hat{K}$) | sd($\hat{K}$) | Mean$(se(\hat{K}))$ | Mean($\hat{K}$) | sd($\hat{K}$) | Mean$(se(\hat{K}))$ | Mean($\hat{K}$) | sd($\hat{K}$) | Mean$(se(\hat{K}))$ |
| -0.2 | -0.171 | -0.2063 | 0.0036 | 0.0042 | -0.171 | 0.0013 | 0.003 | -0.171 | 0.0012 | 0.003 |
| -0.1 | -0.0862 | -0.1297 | 0.0036 | 0.0043 | -0.0862 | 0.0013 | 0.003 | -0.0862 | 0.0013 | 0.003 |
| 0 | 0 | -0.0503 | 0.0036 | 0.0043 | 0 | 0.0013 | 0.0031 | 0 | 0.0013 | 0.0031 |
| 0.1 | 0.0856 | 0.0315 | 0.0035 | 0.0043 | 0.0855 | 0.0014 | 0.0032 | 0.0856 | 0.0014 | 0.0032 |
| 0.2 | 0.1687 | 0.1144 | 0.0035 | 0.0043 | 0.1687 | 0.0014 | 0.0033 | 0.1687 | 0.0014 | 0.0033 |

Table S13: Results of $\mu_\alpha = 0$, $\sigma_\alpha = 0.1$: $K_{YX}$

| | | CD-Ratio | | | CD-Egger | | | CD-GLS | | |
|---|---|---|---|---|---|---|---|---|---|---|
| $\beta_{YX}$ | $K_{YX}$ | Mean($\hat{K}$) | sd($\hat{K}$) | Mean$(se(\hat{K}))$ | Mean($\hat{K}$) | sd($\hat{K}$) | Mean$(se(\hat{K}))$ | Mean($\hat{K}$) | sd($\hat{K}$) | Mean$(se(\hat{K}))$ |
| -0.2 | -0.442 | -0.4254 | 0.0477 | 0.0037 | -0.4335 | 0.0345 | 0.0386 | -0.4335 | 0.0347 | 0.0394 |
| -0.1 | -0.2344 | -0.2233 | 0.0545 | 0.0037 | -0.2296 | 0.0409 | 0.0408 | -0.2295 | 0.0414 | 0.0416 |
| 0 | 0 | 0.0013 | 0.0563 | 0.0037 | -0.0007 | 0.0431 | 0.041 | -0.0005 | 0.0436 | 0.0419 |
| 0.1 | 0.2257 | 0.2181 | 0.0531 | 0.0037 | 0.2201 | 0.0397 | 0.0393 | 0.2203 | 0.0402 | 0.0401 |
| 0.2 | 0.4139 | 0.4019 | 0.0459 | 0.0037 | 0.4056 | 0.0332 | 0.0362 | 0.4059 | 0.0334 | 0.0369 |

Table S14: Results of $\mu_\alpha = 0.2$, $\sigma_\alpha = 0.1$: $K_{YX}$

| | | CD-Ratio | | | CD-Egger | | | CD-GLS | | |
|---|---|---|---|---|---|---|---|---|---|---|
| $\beta_{YX}$ | $K_{YX}$ | Mean($\hat{K}$) | sd($\hat{K}$) | Mean$(se(\hat{K}))$ | Mean($\hat{K}$) | sd($\hat{K}$) | Mean$(se(\hat{K}))$ | Mean($\hat{K}$) | sd($\hat{K}$) | Mean$(se(\hat{K}))$ |
| -0.2 | -0.3995 | -0.4048 | 0.0463 | 0.0038 | -0.3936 | 0.0328 | 0.035 | -0.3936 | 0.0326 | 0.0357 |
| -0.1 | -0.2093 | -0.2172 | 0.0517 | 0.0039 | -0.2061 | 0.037 | 0.0366 | -0.206 | 0.0373 | 0.0373 |
| 0 | 0 | -0.0112 | 0.0538 | 0.0039 | -0.0005 | 0.0386 | 0.0368 | -0.0004 | 0.0392 | 0.0375 |
| 0.1 | 0.2027 | 0.1898 | 0.0514 | 0.0039 | 0.1987 | 0.0369 | 0.0355 | 0.1989 | 0.0375 | 0.0362 |
| 0.2 | 0.3773 | 0.3612 | 0.0452 | 0.0039 | 0.3711 | 0.0326 | 0.0331 | 0.3714 | 0.0331 | 0.0338 |

Table S15: Results of $\mu_\alpha = 0.4$, $\sigma_\alpha = 0.1$: $K_{YX}$

| | | CD-Ratio | | | CD-Egger | | | CD-GLS | | |
|---|---|---|---|---|---|---|---|---|---|---|
| $\beta_{YX}$ | $K_{YX}$ | Mean($\hat{K}$) | sd($\hat{K}$) | Mean$(se(\hat{K}))$ | Mean($\hat{K}$) | sd($\hat{K}$) | Mean$(se(\hat{K}))$ | Mean($\hat{K}$) | sd($\hat{K}$) | Mean$(se(\hat{K}))$ |
| -0.2 | -0.3203 | -0.3301 | 0.0412 | 0.004 | -0.3176 | 0.028 | 0.0282 | -0.3176 | 0.028 | 0.0288 |
| -0.1 | -0.1649 | -0.1796 | 0.0435 | 0.0041 | -0.1636 | 0.0298 | 0.029 | -0.1635 | 0.03 | 0.0296 |
| 0 | 0 | -0.0183 | 0.0448 | 0.0041 | -0.0004 | 0.0306 | 0.0291 | -0.0003 | 0.0311 | 0.0297 |
| 0.1 | 0.1614 | 0.1438 | 0.0448 | 0.0041 | 0.1593 | 0.0301 | 0.0285 | 0.1595 | 0.0307 | 0.029 |
| 0.2 | 0.3079 | 0.2925 | 0.0424 | 0.0041 | 0.3046 | 0.0284 | 0.0272 | 0.3049 | 0.029 | 0.0278 |

Table S16: Results of $\mu_\alpha = 0.6$, $\sigma_\alpha = 0.1$: $K_{YX}$

| | | CD-Ratio | | | CD-Egger | | | CD-GLS | | |
|---|---|---|---|---|---|---|---|---|---|---|
| $\beta_{YX}$ | $K_{YX}$ | Mean($\hat{K}$) | sd($\hat{K}$) | Mean$(se(\hat{K}))$ | Mean($\hat{K}$) | sd($\hat{K}$) | Mean$(se(\hat{K}))$ | Mean($\hat{K}$) | sd($\hat{K}$) | Mean$(se(\hat{K}))$ |
| -0.2 | -0.2535 | -0.2721 | 0.0341 | 0.0041 | -0.2524 | 0.0228 | 0.0224 | -0.2524 | 0.0228 | 0.0229 |
| -0.1 | -0.129 | -0.1552 | 0.0351 | 0.0042 | -0.1285 | 0.0236 | 0.0228 | -0.1285 | 0.0238 | 0.0233 |
| 0 | 0 | -0.0317 | 0.0357 | 0.0042 | -0.0003 | 0.024 | 0.0228 | -0.0002 | 0.0244 | 0.0233 |
| 0.1 | 0.1272 | 0.0944 | 0.0362 | 0.0042 | 0.1261 | 0.024 | 0.0225 | 0.1263 | 0.0244 | 0.023 |
| 0.2 | 0.2469 | 0.217 | 0.0359 | 0.0042 | 0.2451 | 0.0234 | 0.0219 | 0.2454 | 0.0239 | 0.0224 |

Table S17: Results of $\mu_\alpha = 1$, $\sigma_\alpha = 0.1$: $K_{YX}$

| | | CD-Ratio | | | CD-Egger | | | CD-GLS | | |
|---|---|---|---|---|---|---|---|---|---|---|
| $\beta_{YX}$ | $K_{YX}$ | Mean($\hat{K}$) | sd($\hat{K}$) | Mean$(se(\hat{K}))$ | Mean($\hat{K}$) | sd($\hat{K}$) | Mean$(se(\hat{K}))$ | Mean($\hat{K}$) | sd($\hat{K}$) | Mean$(se(\hat{K}))$ |
| -0.2 | -0.171 | -0.2066 | 0.0236 | 0.0042 | -0.1708 | 0.0157 | 0.0151 | -0.1708 | 0.0158 | 0.0154 |
| -0.1 | -0.0862 | -0.1301 | 0.0238 | 0.0043 | -0.0862 | 0.0159 | 0.0153 | -0.0862 | 0.0161 | 0.0156 |
| 0 | 0 | -0.0506 | 0.0241 | 0.0043 | -0.0002 | 0.0161 | 0.0153 | -0.0002 | 0.0164 | 0.0156 |
| 0.1 | 0.0856 | 0.0313 | 0.0244 | 0.0043 | 0.0851 | 0.0162 | 0.0152 | 0.0852 | 0.0165 | 0.0155 |
| 0.2 | 0.1687 | 0.1143 | 0.0247 | 0.0043 | 0.1681 | 0.0161 | 0.015 | 0.1682 | 0.0165 | 0.0153 |

Table S18: Results of $\mu_\alpha = 0$, $\sigma_\alpha = 0.2$: $K_{YX}$

| | | CD-Ratio | | | CD-Egger | | | CD-GLS | | |
|---|---|---|---|---|---|---|---|---|---|---|
| $\beta_{YX}$ | $K_{YX}$ | Mean($\hat{K}$) | sd($\hat{K}$) | Mean$(se(\hat{K}))$ | Mean($\hat{K}$) | sd($\hat{K}$) | Mean$(se(\hat{K}))$ | Mean($\hat{K}$) | sd($\hat{K}$) | Mean$(se(\hat{K}))$ |
| -0.2 | -0.442 | -0.3933 | 0.0936 | 0.0038 | -0.4113 | 0.0669 | 0.0727 | -0.4111 | 0.0674 | 0.0742 |
| -0.1 | -0.2344 | -0.2037 | 0.1043 | 0.0039 | -0.2168 | 0.0771 | 0.0763 | -0.2166 | 0.0781 | 0.078 |
| 0 | 0 | 0.0031 | 0.107 | 0.0038 | -0.0017 | 0.0804 | 0.0768 | -0.0015 | 0.0815 | 0.0784 |
| 0.1 | 0.2257 | 0.2031 | 0.1021 | 0.0038 | 0.2066 | 0.0751 | 0.0738 | 0.2068 | 0.076 | 0.0754 |
| 0.2 | 0.4139 | 0.3767 | 0.0909 | 0.0038 | 0.385 | 0.0644 | 0.0685 | 0.3853 | 0.0648 | 0.07 |

Table S19: Results of $\mu_\alpha = 0.2$, $\sigma_\alpha = 0.2$: $K_{YX}$

| | | CD-Ratio | | | CD-Egger | | | CD-GLS | | |
|---|---|---|---|---|---|---|---|---|---|---|
| $\beta_{YX}$ | $K_{YX}$ | Mean($\hat{K}$) | sd($\hat{K}$) | Mean$(se(\hat{K}))$ | Mean($\hat{K}$) | sd($\hat{K}$) | Mean$(se(\hat{K}))$ | Mean($\hat{K}$) | sd($\hat{K}$) | Mean$(se(\hat{K}))$ |
| -0.2 | -0.3995 | -0.3871 | 0.089 | 0.0039 | -0.3775 | 0.0632 | 0.0668 | -0.3773 | 0.0633 | 0.0682 |
| -0.1 | -0.2093 | -0.2079 | 0.099 | 0.0039 | -0.1971 | 0.0705 | 0.0695 | -0.1968 | 0.0712 | 0.071 |
| 0 | 0 | -0.0129 | 0.1023 | 0.0039 | -0.0012 | 0.0733 | 0.0698 | -0.0009 | 0.0744 | 0.0712 |
| 0.1 | 0.2027 | 0.1764 | 0.0977 | 0.0039 | 0.1889 | 0.0703 | 0.0675 | 0.1893 | 0.0715 | 0.0689 |
| 0.2 | 0.3773 | 0.3409 | 0.0874 | 0.0039 | 0.3555 | 0.0629 | 0.0633 | 0.3559 | 0.0639 | 0.0646 |

Table S20: Results of $\mu_\alpha = 0.4$, $\sigma_\alpha = 0.2$: $K_{YX}$

| | | CD-Ratio | | | CD-Egger | | | CD-GLS | | |
|---|---|---|---|---|---|---|---|---|---|---|
| $\beta_{YX}$ | $K_{YX}$ | Mean($\hat{K}$) | sd($\hat{K}$) | Mean$(se(\hat{K}))$ | Mean($\hat{K}$) | sd($\hat{K}$) | Mean$(se(\hat{K}))$ | Mean($\hat{K}$) | sd($\hat{K}$) | Mean$(se(\hat{K}))$ |
| -0.2 | -0.3203 | -0.3305 | 0.0782 | 0.004 | -0.3097 | 0.0546 | 0.0548 | -0.3096 | 0.0546 | 0.0559 |
| -0.1 | -0.1649 | -0.1836 | 0.0837 | 0.004 | -0.1594 | 0.0578 | 0.0562 | -0.1592 | 0.0584 | 0.0574 |
| 0 | 0 | -0.0248 | 0.0871 | 0.004 | -0.0009 | 0.0593 | 0.0563 | -0.0006 | 0.0603 | 0.0575 |
| 0.1 | 0.1614 | 0.1345 | 0.0868 | 0.0041 | 0.1544 | 0.0584 | 0.0551 | 0.1547 | 0.0596 | 0.0562 |
| 0.2 | 0.3079 | 0.2805 | 0.082 | 0.0041 | 0.2962 | 0.0552 | 0.0527 | 0.2965 | 0.0564 | 0.0538 |

Table S21: Results of $\mu_\alpha = 0.6$, $\sigma_\alpha = 0.2$: $K_{YX}$

| | | CD-Ratio | | | CD-Egger | | | CD-GLS | | |
|---|---|---|---|---|---|---|---|---|---|---|
| $\beta_{YX}$ | $K_{YX}$ | Mean($\hat{K}$) | sd($\hat{K}$) | Mean$(se(\hat{K}))$ | Mean($\hat{K}$) | sd($\hat{K}$) | Mean$(se(\hat{K}))$ | Mean($\hat{K}$) | sd($\hat{K}$) | Mean$(se(\hat{K}))$ |
| -0.2 | -0.2535 | -0.2754 | 0.0659 | 0.0041 | -0.2487 | 0.0448 | 0.044 | -0.2486 | 0.0449 | 0.0449 |
| -0.1 | -0.129 | -0.1595 | 0.0685 | 0.0041 | -0.1268 | 0.0463 | 0.0447 | -0.1266 | 0.0468 | 0.0456 |
| 0 | 0 | -0.0361 | 0.0706 | 0.0042 | -0.0007 | 0.0471 | 0.0447 | -0.0005 | 0.0479 | 0.0457 |
| 0.1 | 0.1272 | 0.0901 | 0.0716 | 0.0042 | 0.1236 | 0.047 | 0.0441 | 0.1239 | 0.048 | 0.045 |
| 0.2 | 0.2469 | 0.2122 | 0.0706 | 0.0042 | 0.2408 | 0.0459 | 0.0428 | 0.2411 | 0.0469 | 0.0437 |

Table S22: Results of $\mu_\alpha = 1$, $\sigma_\alpha = 0.2$: $K_{YX}$

| | | CD-Ratio | | | CD-Egger | | | CD-GLS | | |
|---|---|---|---|---|---|---|---|---|---|---|
| $\beta_{YX}$ | $K_{YX}$ | Mean($\hat{K}$) | sd($\hat{K}$) | Mean$\big(se(\hat{K})\big)$ | Mean($\hat{K}$) | sd($\hat{K}$) | Mean$\big(se(\hat{K})\big)$ | Mean($\hat{K}$) | sd($\hat{K}$) | Mean$\big(se(\hat{K})\big)$ |
| -0.2 | -0.171 | -0.2087 | 0.0464 | 0.0042 | -0.1699 | 0.031 | 0.03 | -0.1699 | 0.0312 | 0.0306 |
| -0.1 | -0.0862 | -0.1321 | 0.0471 | 0.0042 | -0.0859 | 0.0315 | 0.0302 | -0.0858 | 0.0319 | 0.0309 |
| 0 | 0 | -0.0523 | 0.0479 | 0.0043 | -0.0005 | 0.0318 | 0.0302 | -0.0004 | 0.0324 | 0.0309 |
| 0.1 | 0.0856 | 0.0301 | 0.0486 | 0.0043 | 0.0842 | 0.032 | 0.03 | 0.0843 | 0.0326 | 0.0307 |
| 0.2 | 0.1687 | 0.1134 | 0.0492 | 0.0043 | 0.1665 | 0.0319 | 0.0296 | 0.1667 | 0.0326 | 0.0303 |

#### 6.3 Goodness-of-Fit Tests

We show the results of the Goodness-of-Fit tests for CD-Ratio, CD-Egger, and CD-GLS. Table S23 shows the results under setup $\mu_\alpha = 0, \sigma_\alpha = 0$, and various levels of $\beta_{YX}$. For each of the three methods, in each of 500 simulations we can calculate the test statistics $Q^{YX}$ and $Q^{XY}$, then perform the corresponding $\chi^2$-test to get $p$-value $p_{YX}$ for direction $X$ to $Y$, and $p$-value $p_{XY}$ for direction $Y$ to $X$, respectively. Then we show the proportion out of 500 simulated datasets $p_{YX}$'s less than 0.05, indicated by "$p_{YX} < 0.05$", and proportion of $p_{XY}$'s less than 0.05, indicated by "$p_{XY} < 0.05$". Table S24 to S37 shows the results for other setups.

Table S23: GOF Testing Results of $\mu_\alpha = 0$, $\sigma_\alpha = 0$

| | CD-Ratio | | CD-Egger | | CD-GLS | |
|---|---|---|---|---|---|---|
| $\beta_{YX}$ | $p_{YX} < 0.05$ | $p_{XY} < 0.05$ | $p_{YX} < 0.05$ | $p_{XY} < 0.05$ | $p_{YX} < 0.05$ | $p_{XY} < 0.05$ |
| -0.2 | 0.002 | 0.012 | 0 | 0.014 | 0 | 0 |
| -0.1 | 0.012 | 0.188 | 0 | 0 | 0 | 0 |
| 0 | 0.054 | 0.042 | 0.002 | 0 | 0.002 | 0.026 |
| 0.1 | 0.022 | 0.18 | 0.002 | 0 | 0 | 0 |
| 0.2 | 0 | 0.022 | 0 | 0.006 | 0 | 0 |

Table S24: GOF Results of $\mu_\alpha = 0.2$, $\sigma_\alpha = 0$

| | CD-Ratio | | CD-Egger | | CD-GLS | |
|---|---|---|---|---|---|---|
| $\beta_{YX}$ | $p_{YX} < 0.05$ | $p_{XY} < 0.05$ | $p_{YX} < 0.05$ | $p_{XY} < 0.05$ | $p_{YX} < 0.05$ | $p_{XY} < 0.05$ |
| -0.2 | 1 | 1 | 0 | 0.014 | 0 | 0 |
| -0.1 | 1 | 1 | 0 | 0 | 0 | 0 |
| 0 | 1 | 1 | 0 | 0 | 0 | 0 |
| 0.1 | 1 | 1 | 0 | 0 | 0 | 0 |
| 0.2 | 1 | 1 | 0 | 0.006 | 0 | 0 |

Table S25: GOF Results of $\mu_\alpha = 0.4$, $\sigma_\alpha = 0$

| | CD-Ratio | | CD-Egger | | CD-GLS | |
|---|---|---|---|---|---|---|
| $\beta_{YX}$ | $p_{YX} < 0.05$ | $p_{XY} < 0.05$ | $p_{YX} < 0.05$ | $p_{XY} < 0.05$ | $p_{YX} < 0.05$ | $p_{XY} < 0.05$ |
| -0.2 | 1 | 1 | 0 | 0.014 | 0 | 0 |
| -0.1 | 1 | 1 | 0 | 0 | 0 | 0 |
| 0 | 1 | 1 | 0 | 0 | 0 | 0 |
| 0.1 | 1 | 1 | 0 | 0 | 0 | 0 |
| 0.2 | 1 | 1 | 0 | 0.006 | 0 | 0 |

Table S26: GOF Results of $\mu_\alpha = 0.6$, $\sigma_\alpha = 0$

| | CD-Ratio | | CD-Egger | | CD-GLS | |
|---|---|---|---|---|---|---|
| $\beta_{YX}$ | $p_{YX} < 0.05$ | $p_{XY} < 0.05$ | $p_{YX} < 0.05$ | $p_{XY} < 0.05$ | $p_{YX} < 0.05$ | $p_{XY} < 0.05$ |
| -0.2 | 1 | 1 | 0 | 0.014 | 0 | 0 |
| -0.1 | 1 | 1 | 0 | 0 | 0 | 0 |
| 0 | 1 | 1 | 0 | 0 | 0 | 0 |
| 0.1 | 1 | 1 | 0 | 0 | 0 | 0 |
| 0.2 | 1 | 1 | 0 | 0.006 | 0 | 0 |

Table S27: GOF Results of $\mu_\alpha = 1$, $\sigma_\alpha = 0$

| | CD-Ratio | | CD-Egger | | CD-GLS | |
|---|---|---|---|---|---|---|
| $\beta_{YX}$ | $p_{YX} < 0.05$ | $p_{XY} < 0.05$ | $p_{YX} < 0.05$ | $p_{XY} < 0.05$ | $p_{YX} < 0.05$ | $p_{XY} < 0.05$ |
| -0.2 | 1 | 1 | 0 | 0.014 | 0 | 0 |
| -0.1 | 1 | 1 | 0 | 0 | 0 | 0 |
| 0 | 1 | 1 | 0 | 0 | 0 | 0 |
| 0.1 | 1 | 1 | 0 | 0 | 0 | 0 |
| 0.2 | 1 | 1 | 0 | 0.006 | 0 | 0 |

Table S28: GOF Results of $\mu_\alpha = 0$, $\sigma_\alpha = 0.1$

| | CD-Ratio | | CD-Egger | | CD-GLS | |
|---|---|---|---|---|---|---|
| $\beta_{YX}$ | $p_{YX} < 0.05$ | $p_{XY} < 0.05$ | $p_{YX} < 0.05$ | $p_{XY} < 0.05$ | $p_{YX} < 0.05$ | $p_{XY} < 0.05$ |
| -0.2 | 1 | 1 | 0 | 0 | 0 | 0 |
| -0.1 | 1 | 1 | 0 | 0 | 0 | 0 |
| 0 | 1 | 1 | 0 | 0 | 0 | 0 |
| 0.1 | 1 | 1 | 0 | 0 | 0 | 0 |
| 0.2 | 1 | 1 | 0 | 0 | 0 | 0 |

Table S29: GOF Results of $\mu_\alpha = 0.2$, $\sigma_\alpha = 0.1$

| | CD-Ratio | | CD-Egger | | CD-GLS | |
|---|---|---|---|---|---|---|
| $\beta_{YX}$ | $p_{YX} < 0.05$ | $p_{XY} < 0.05$ | $p_{YX} < 0.05$ | $p_{XY} < 0.05$ | $p_{YX} < 0.05$ | $p_{XY} < 0.05$ |
| -0.2 | 1 | 1 | 0 | 0 | 0 | 0 |
| -0.1 | 1 | 1 | 0 | 0 | 0 | 0 |
| 0 | 1 | 1 | 0 | 0 | 0 | 0 |
| 0.1 | 1 | 1 | 0 | 0 | 0 | 0 |
| 0.2 | 1 | 1 | 0 | 0 | 0 | 0 |

Table S30: GOF Results of $\mu_\alpha = 0.4$, $\sigma_\alpha = 0.1$

| | CD-Ratio | | CD-Egger | | CD-GLS | |
|---|---|---|---|---|---|---|
| $\beta_{YX}$ | $p_{YX} < 0.05$ | $p_{XY} < 0.05$ | $p_{YX} < 0.05$ | $p_{XY} < 0.05$ | $p_{YX} < 0.05$ | $p_{XY} < 0.05$ |
| -0.2 | 1 | 1 | 0 | 0 | 0 | 0 |
| -0.1 | 1 | 1 | 0 | 0 | 0 | 0 |
| 0 | 1 | 1 | 0 | 0 | 0 | 0 |
| 0.1 | 1 | 1 | 0 | 0 | 0 | 0 |
| 0.2 | 1 | 1 | 0 | 0 | 0 | 0 |

Table S31: GOF Results of $\mu_\alpha = 0.6$, $\sigma_\alpha = 0.1$

| | CD-Ratio | | CD-Egger | | CD-GLS | |
|---|---|---|---|---|---|---|
| $\beta_{YX}$ | $p_{YX} < 0.05$ | $p_{XY} < 0.05$ | $p_{YX} < 0.05$ | $p_{XY} < 0.05$ | $p_{YX} < 0.05$ | $p_{XY} < 0.05$ |
| -0.2 | 1 | 1 | 0 | 0 | 0 | 0 |
| -0.1 | 1 | 1 | 0 | 0 | 0 | 0 |
| 0 | 1 | 1 | 0 | 0 | 0 | 0 |
| 0.1 | 1 | 1 | 0 | 0 | 0 | 0 |
| 0.2 | 1 | 1 | 0 | 0 | 0 | 0 |

Table S32: GOF Results of $\mu_\alpha = 1$, $\sigma_\alpha = 0.1$

| | CD-Ratio | | CD-Egger | | CD-GLS | |
|---|---|---|---|---|---|---|
| $\beta_{YX}$ | $p_{YX} < 0.05$ | $p_{XY} < 0.05$ | $p_{YX} < 0.05$ | $p_{XY} < 0.05$ | $p_{YX} < 0.05$ | $p_{XY} < 0.05$ |
| -0.2 | 1 | 1 | 0 | 0 | 0 | 0 |
| -0.1 | 1 | 1 | 0 | 0 | 0 | 0 |
| 0 | 1 | 1 | 0 | 0 | 0 | 0 |
| 0.1 | 1 | 1 | 0 | 0 | 0 | 0 |
| 0.2 | 1 | 1 | 0 | 0 | 0 | 0 |

Table S33: GOF Results of $\mu_\alpha = 0$, $\sigma_\alpha = 0.2$

| | CD-Ratio | | CD-Egger | | CD-GLS | |
|---|---|---|---|---|---|---|
| $\beta_{YX}$ | $p_{YX} < 0.05$ | $p_{XY} < 0.05$ | $p_{YX} < 0.05$ | $p_{XY} < 0.05$ | $p_{YX} < 0.05$ | $p_{XY} < 0.05$ |
| -0.2 | 1 | 1 | 0 | 0 | 0 | 0 |
| -0.1 | 1 | 1 | 0 | 0 | 0 | 0 |
| 0 | 1 | 1 | 0 | 0 | 0 | 0 |
| 0.1 | 1 | 1 | 0 | 0 | 0 | 0 |
| 0.2 | 1 | 1 | 0 | 0 | 0 | 0 |

Table S34: GOF Results of $\mu_\alpha = 0.2$, $\sigma_\alpha = 0.2$

| | CD-Ratio | | CD-Egger | | CD-GLS | |
|---|---|---|---|---|---|---|
| $\beta_{YX}$ | $p_{YX} < 0.05$ | $p_{XY} < 0.05$ | $p_{YX} < 0.05$ | $p_{XY} < 0.05$ | $p_{YX} < 0.05$ | $p_{XY} < 0.05$ |
| -0.2 | 1 | 1 | 0 | 0 | 0 | 0 |
| -0.1 | 1 | 1 | 0 | 0 | 0 | 0 |
| 0 | 1 | 1 | 0 | 0 | 0 | 0 |
| 0.1 | 1 | 1 | 0 | 0 | 0 | 0 |
| 0.2 | 1 | 1 | 0 | 0 | 0 | 0 |

Table S35: GOF Results of $\mu_\alpha = 0.4$, $\sigma_\alpha = 0.2$

| | CD-Ratio | | CD-Egger | | CD-GLS | |
|---|---|---|---|---|---|---|
| $\beta_{YX}$ | $p_{YX} < 0.05$ | $p_{XY} < 0.05$ | $p_{YX} < 0.05$ | $p_{XY} < 0.05$ | $p_{YX} < 0.05$ | $p_{XY} < 0.05$ |
| -0.2 | 1 | 1 | 0 | 0 | 0 | 0 |
| -0.1 | 1 | 1 | 0 | 0 | 0 | 0 |
| 0 | 1 | 1 | 0 | 0 | 0 | 0 |
| 0.1 | 1 | 1 | 0 | 0 | 0 | 0 |
| 0.2 | 1 | 1 | 0 | 0 | 0 | 0 |

Table S36: GOF Results of $\mu_\alpha = 0.6$, $\sigma_\alpha = 0.2$

| | CD-Ratio | | CD-Egger | | CD-GLS | |
|---|---|---|---|---|---|---|
| $\beta_{YX}$ | $p_{YX} < 0.05$ | $p_{XY} < 0.05$ | $p_{YX} < 0.05$ | $p_{XY} < 0.05$ | $p_{YX} < 0.05$ | $p_{XY} < 0.05$ |
| -0.2 | 1 | 1 | 0 | 0 | 0 | 0 |
| -0.1 | 1 | 1 | 0 | 0 | 0 | 0 |
| 0 | 1 | 1 | 0 | 0 | 0 | 0 |
| 0.1 | 1 | 1 | 0 | 0 | 0 | 0 |
| 0.2 | 1 | 1 | 0 | 0 | 0 | 0 |

Table S37: GOF Results of $\mu_\alpha = 1$, $\sigma_\alpha = 0.2$

| | CD-Ratio | | CD-Egger | | CD-GLS | |
|---|---|---|---|---|---|---|
| $\beta_{YX}$ | $p_{YX} < 0.05$ | $p_{XY} < 0.05$ | $p_{YX} < 0.05$ | $p_{XY} < 0.05$ | $p_{YX} < 0.05$ | $p_{XY} < 0.05$ |
| -0.2 | 1 | 1 | 0 | 0 | 0 | 0 |
| -0.1 | 1 | 1 | 0 | 0 | 0 | 0 |
| 0 | 1 | 1 | 0 | 0 | 0 | 0 |
| 0.1 | 1 | 1 | 0 | 0 | 0 | 0 |
| 0.2 | 1 | 1 | 0 | 0 | 0 | 0 |

#### 6.4 Comparing the Decisions

We compare CD-Ratio, CD-Egger and CD-GLS with MR Steiger's and bi-directional MR in terms of their decisions made on the causal direction. Similar to Steiger's method, we also applied CD-Ratio to each of the 22 SNPs, then calculated the proportions of the three possible decisions, denoted by "CD-Ratio-Prop". We also pooled together the individual results from each of the 22 SNPs by majority voting to reach a final decision, denoted as "CD-Ratio-MV". For other methods, their notations are the same as in Figure 7 in the main text. Figure S3 shows the comparison results for setup $\beta_{YX} = 0, \mu_\alpha = 0$, and $\sigma_\alpha = 0, 0.1, 0.2$. Figure S4 to S27 show the comparison results for other simulation setups.

Figure S3: Relative frequencies of decisions for causal direction of all methods: $\beta_{YX} = 0$, $\mu_\alpha = 0$

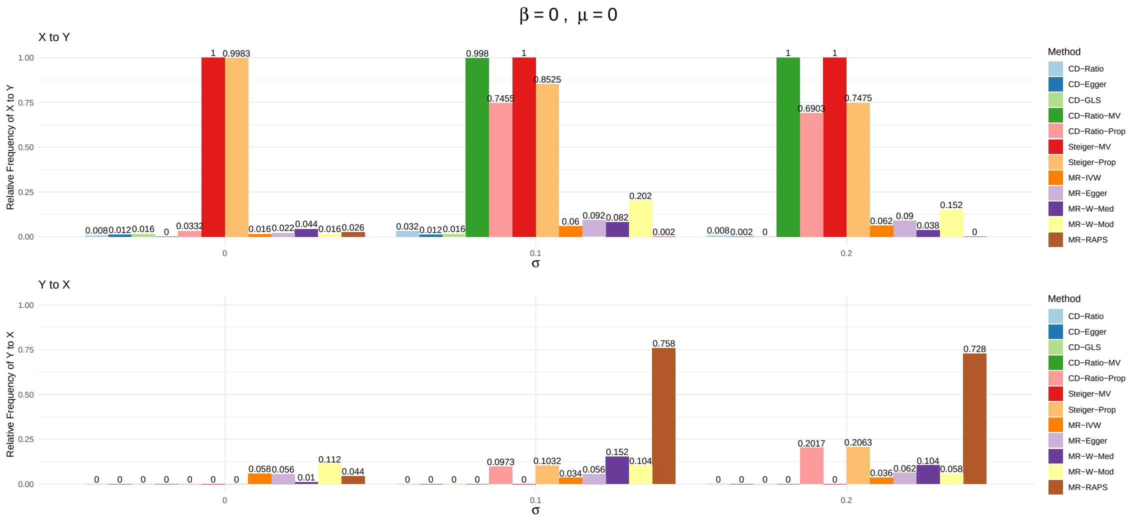

Figure S4: Relative frequencies of decisions for causal direction of all methods: $\beta_{YX} = 0$, $\mu_\alpha = 0.2$

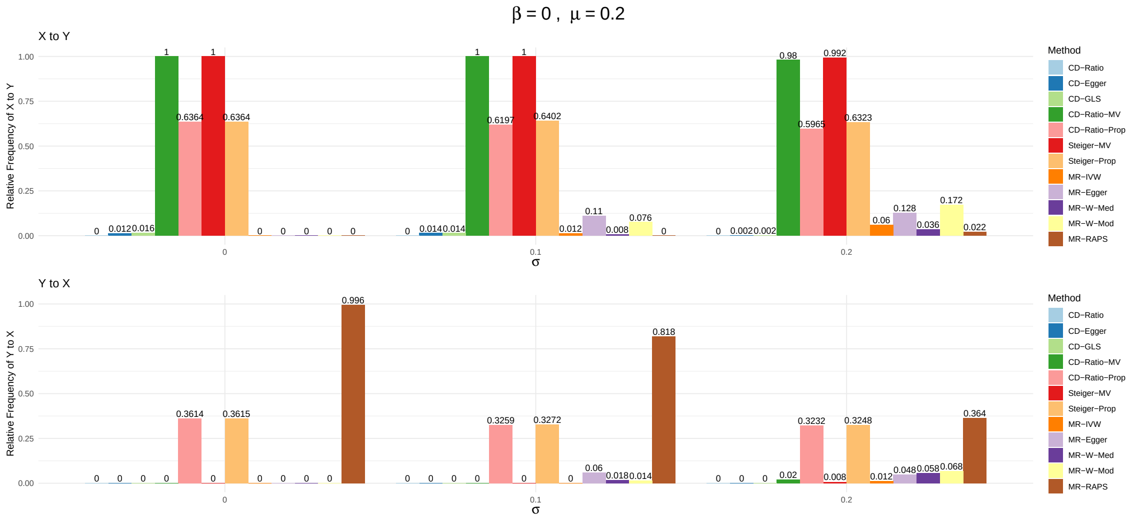


Figure S5: Relative frequencies of decisions for causal direction of all methods: $\beta_{YX} = 0$, $\mu_\alpha = 0.4$

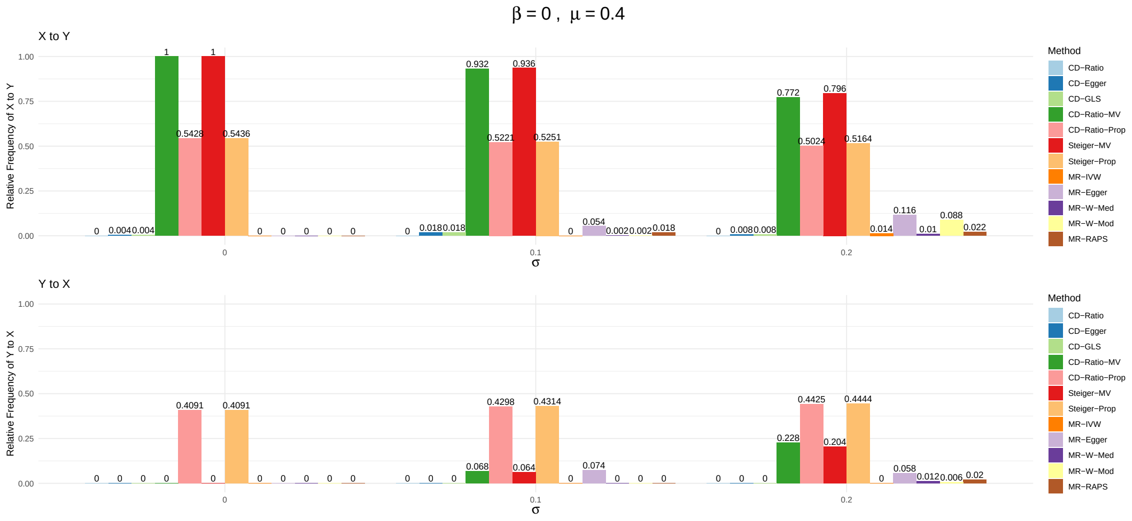

Figure S6: Relative frequencies of decisions for causal direction of all methods: $\beta_{YX} = 0$, $\mu_\alpha = 0.6$

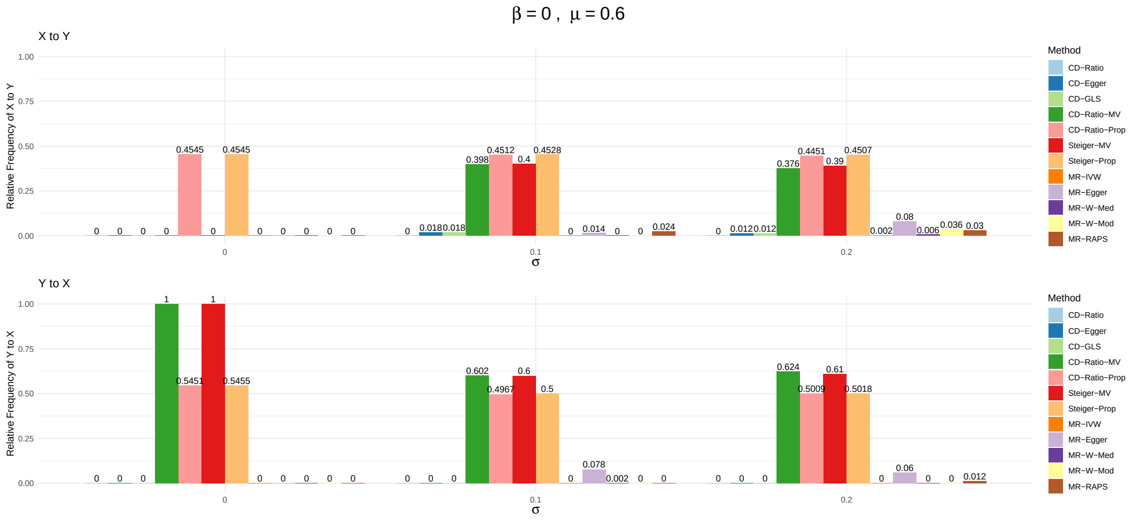


Figure S7: Relative frequencies of decisions for causal direction of all methods: $\beta_{YX} = 0$, $\mu_\alpha = 1$

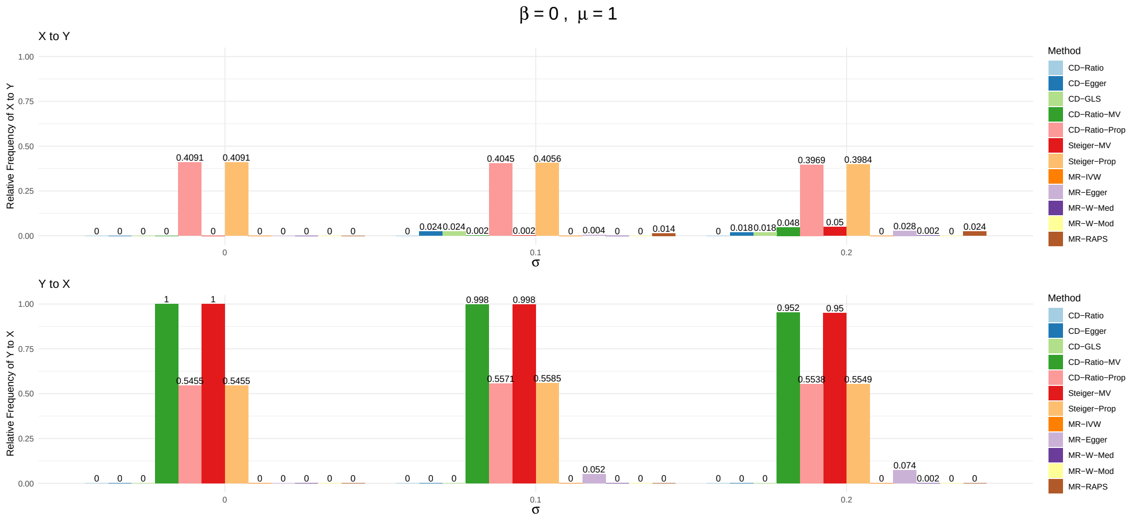

Figure S8: Relative frequencies of decisions for causal direction of all methods: $\beta_{YX} = 0.1$, $\mu_\alpha = 0$

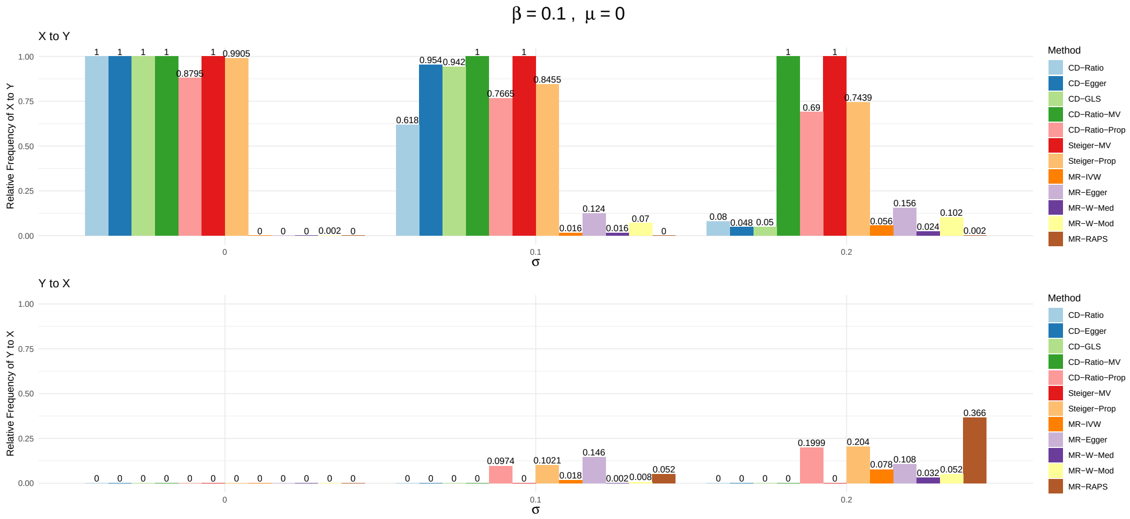


Figure S9: Relative frequencies of decisions for causal direction of all methods: $\beta_{YX} = 0.1$, $\mu_\alpha = 0.2$

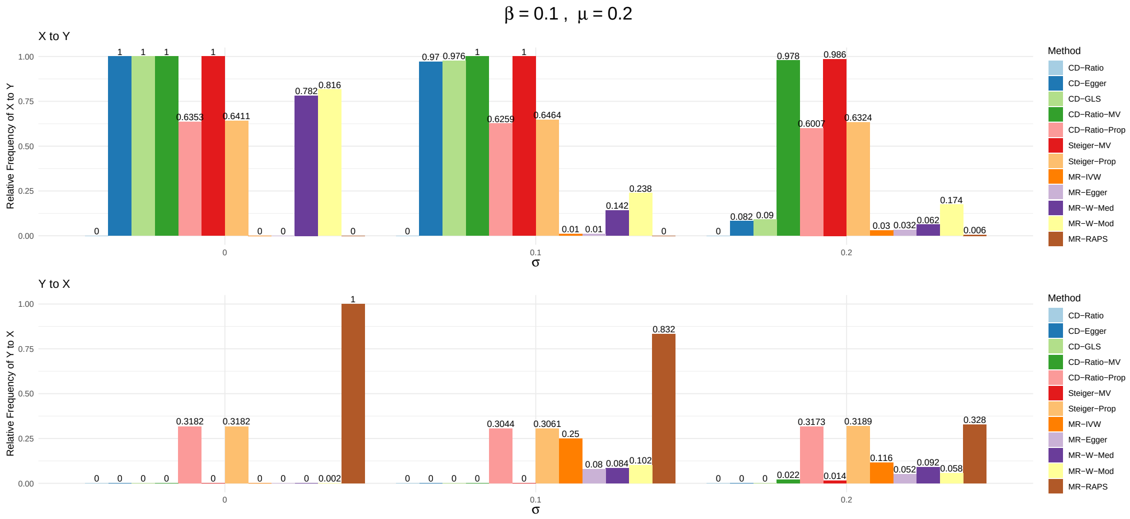

Figure S10: Relative frequencies of decisions for causal direction of all methods: $\beta_{YX} = 0.1$, $\mu_\alpha = 0.4$

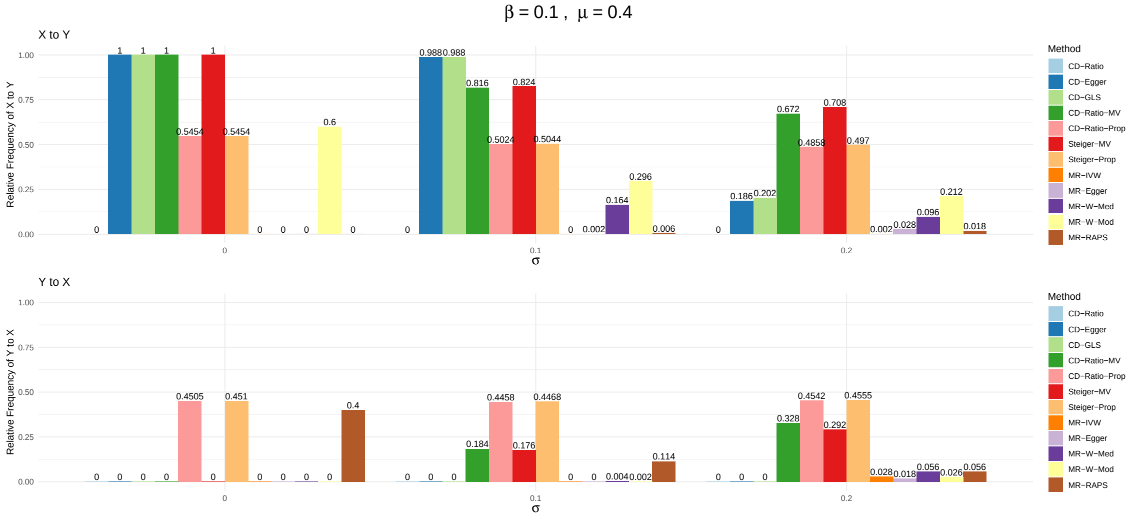


Figure S11: Relative frequencies of decisions for causal direction of all methods: $\beta_{YX} = 0.1$, $\mu_\alpha = 0.6$

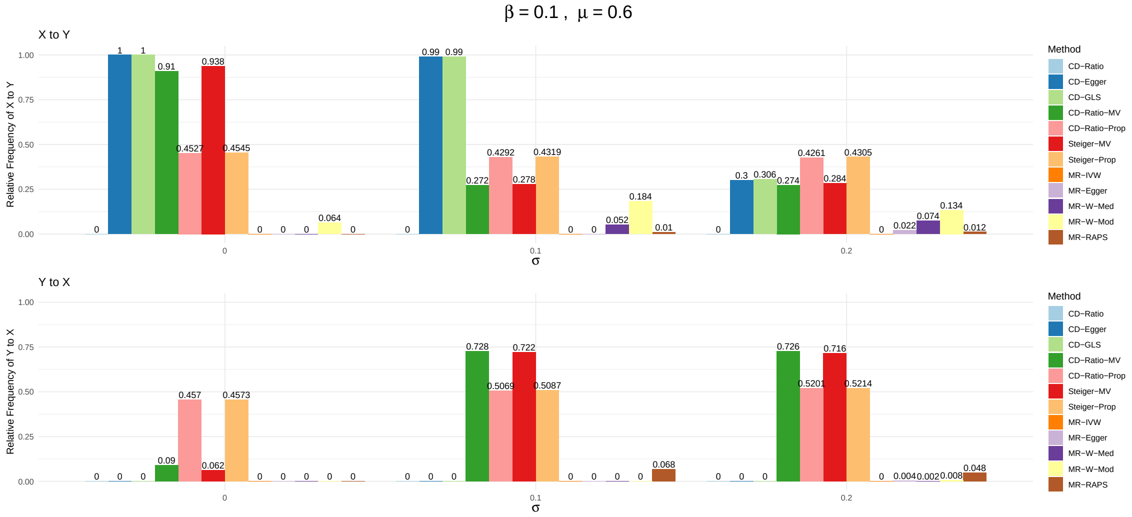

Figure S12: Relative frequencies of decisions for causal direction of all methods: $\beta_{YX} = 0.1$, $\mu_\alpha = 1$

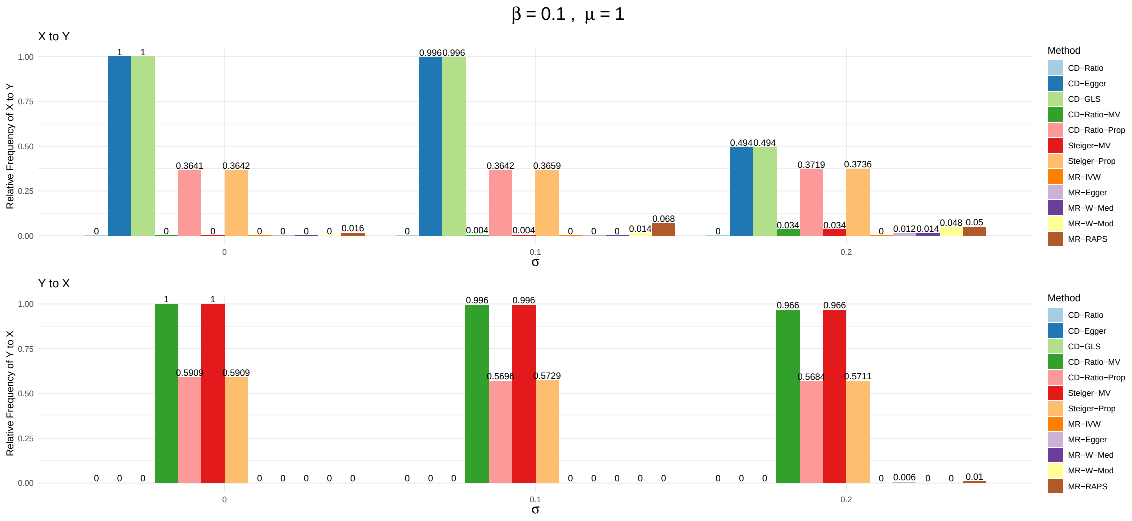


Figure S13: Relative frequencies of decisions for causal direction of all methods: $\beta_{YX} = -0.1$, $\mu_\alpha = 0$

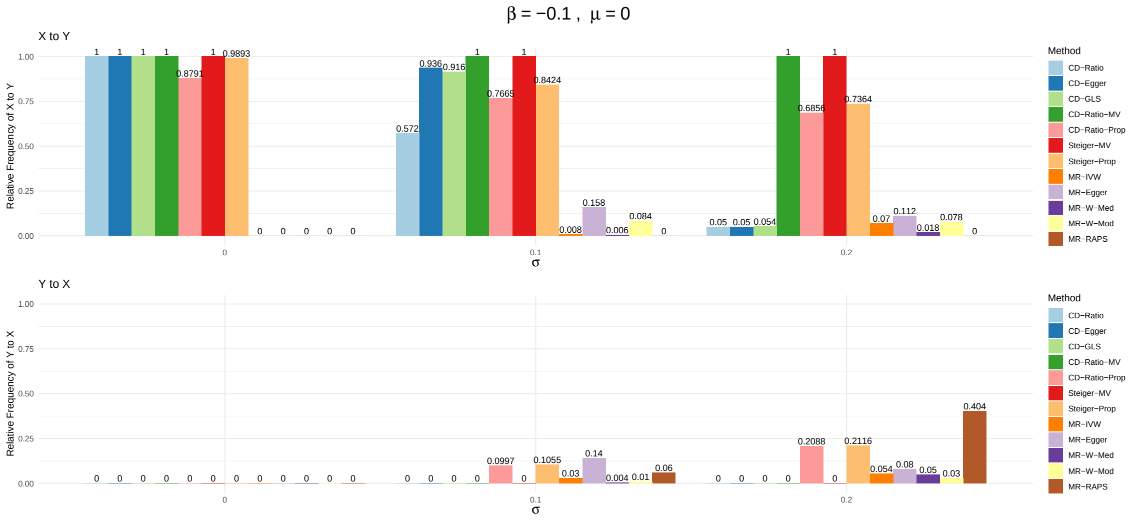

Figure S14: Relative frequencies of decisions for causal direction of all methods: $\beta_{YX} = -0.1$, $\mu_\alpha = 0.2$

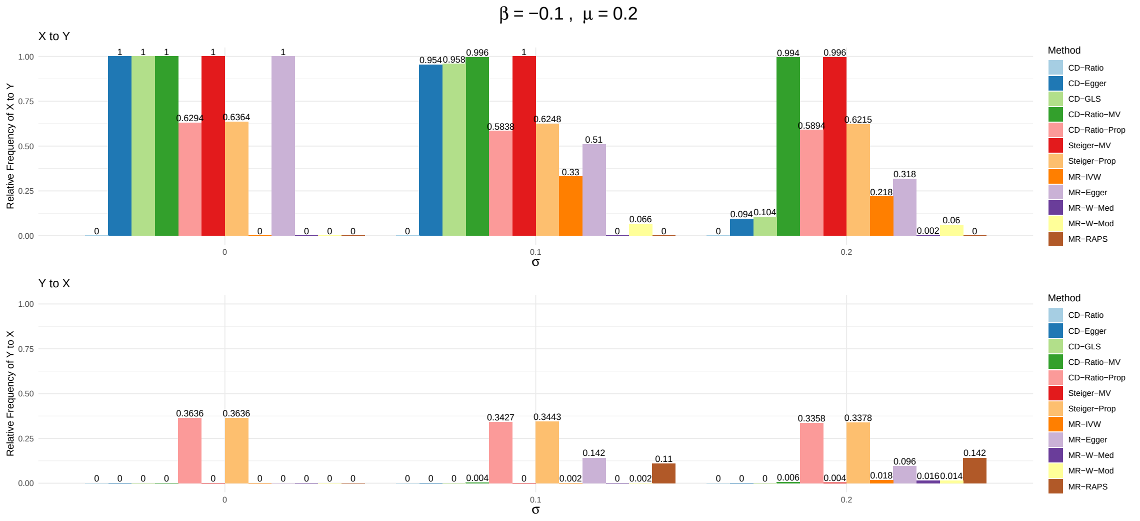


Figure S15: Relative frequencies of decisions for causal direction of all methods: $\beta_{YX} = -0.1$, $\mu_\alpha = 0.4$

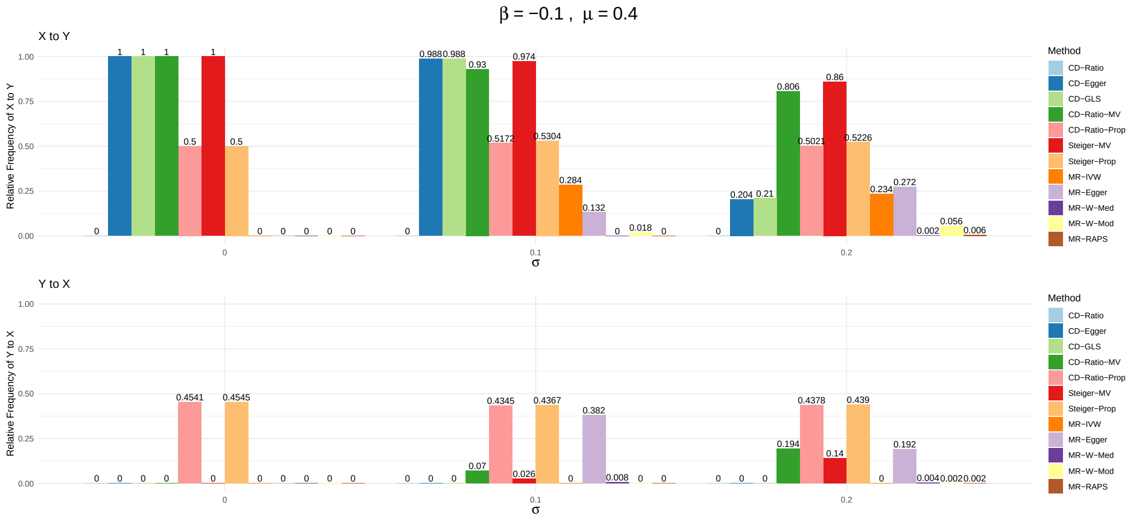

Figure S16: Relative frequencies of decisions for causal direction of all methods: $\beta_{YX} = -0.1$, $\mu_\alpha = 0.6$

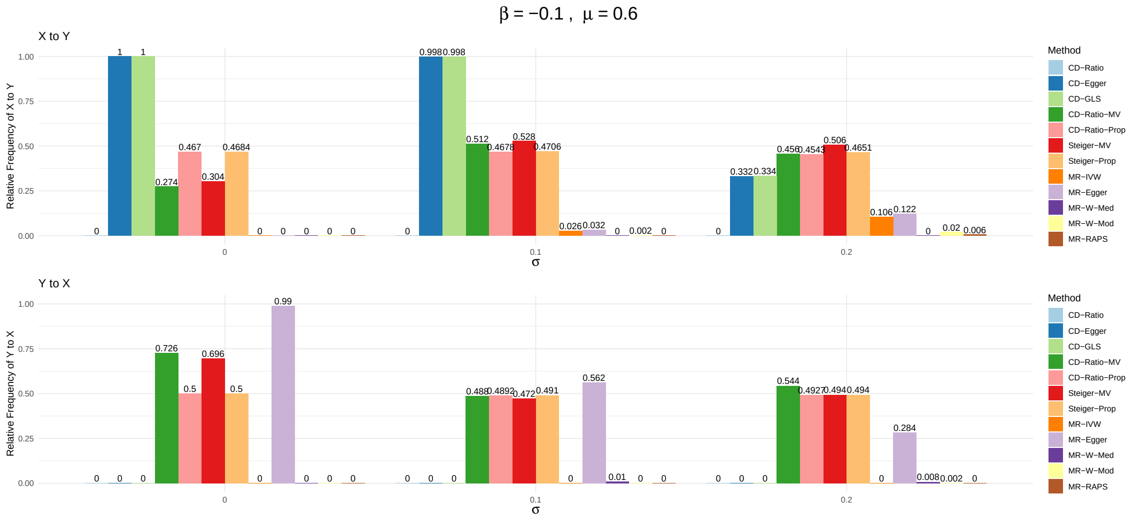


Figure S17: Relative frequencies of decisions for causal direction of all methods: $\beta_{YX} = -0.1$, $\mu_\alpha = 1$

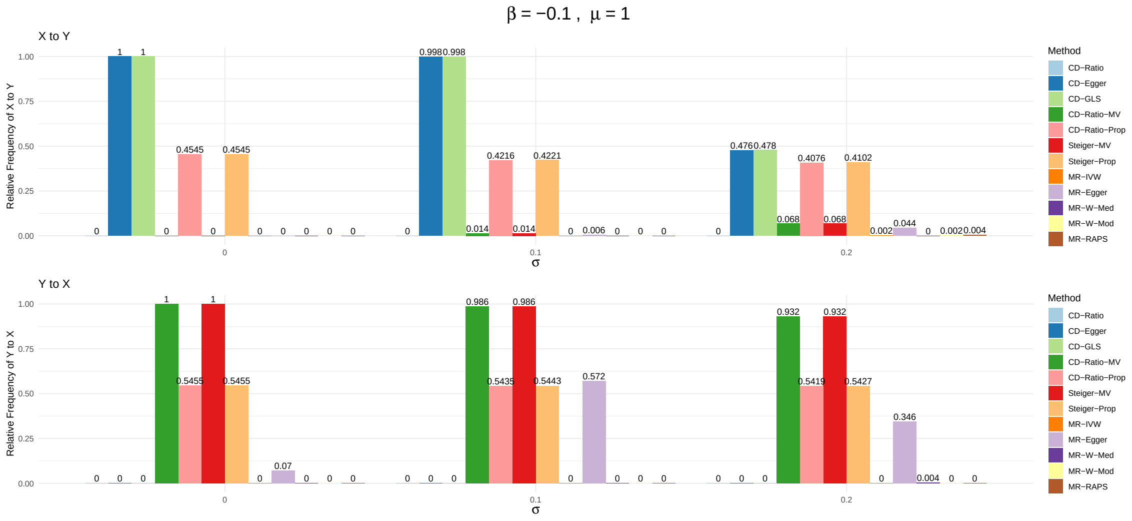

Figure S18: Relative frequencies of decisions for causal direction of all methods: $\beta_{YX} = 0.2$, $\mu_\alpha = 0$

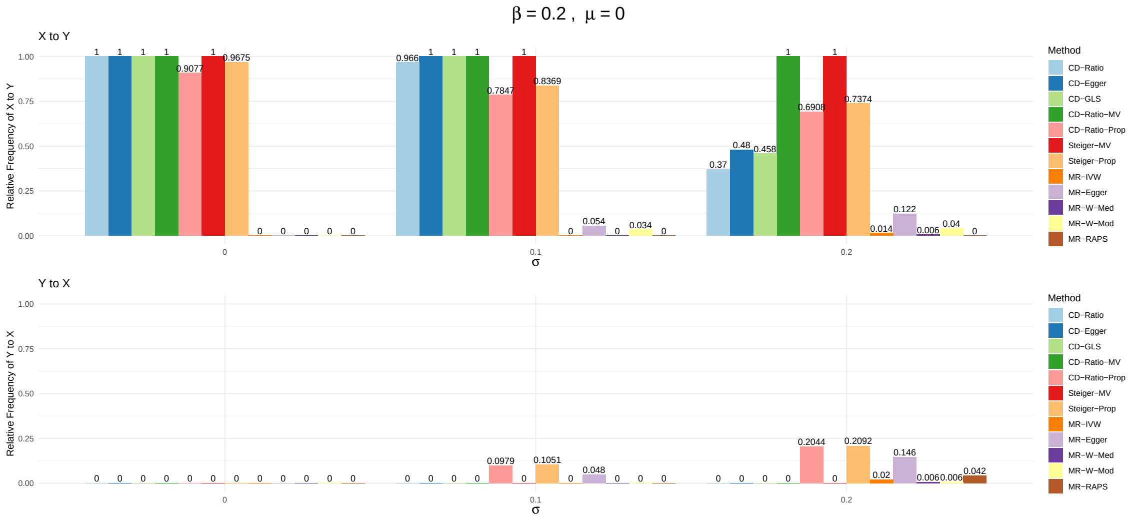


Figure S19: Relative frequencies of decisions for causal direction of all methods: $\beta_{YX} = 0.2$, $\mu_\alpha = 0.2$

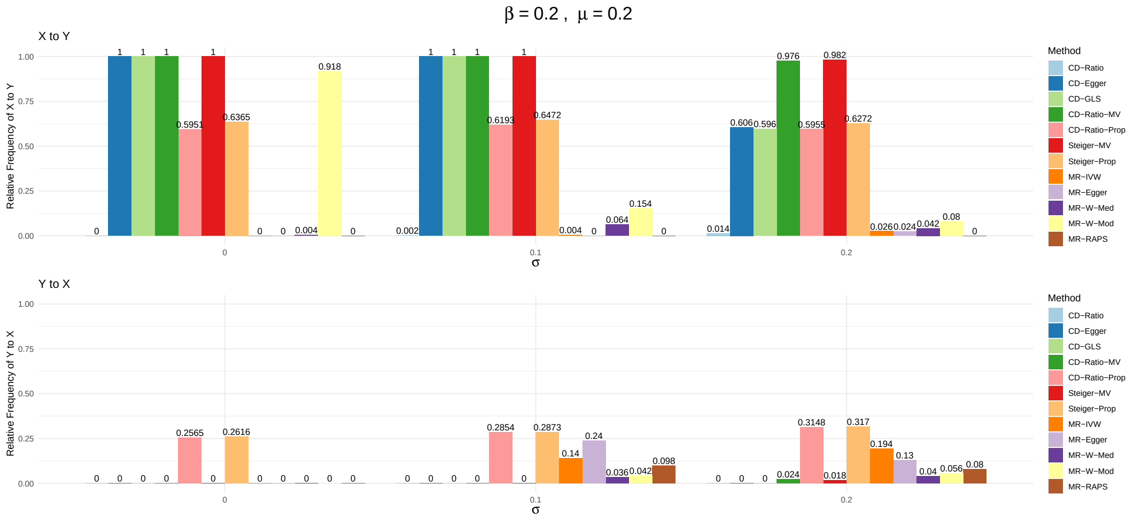

Figure S20: Relative frequencies of decisions for causal direction of all methods: $\beta_{YX} = 0.2$, $\mu_\alpha = 0.4$

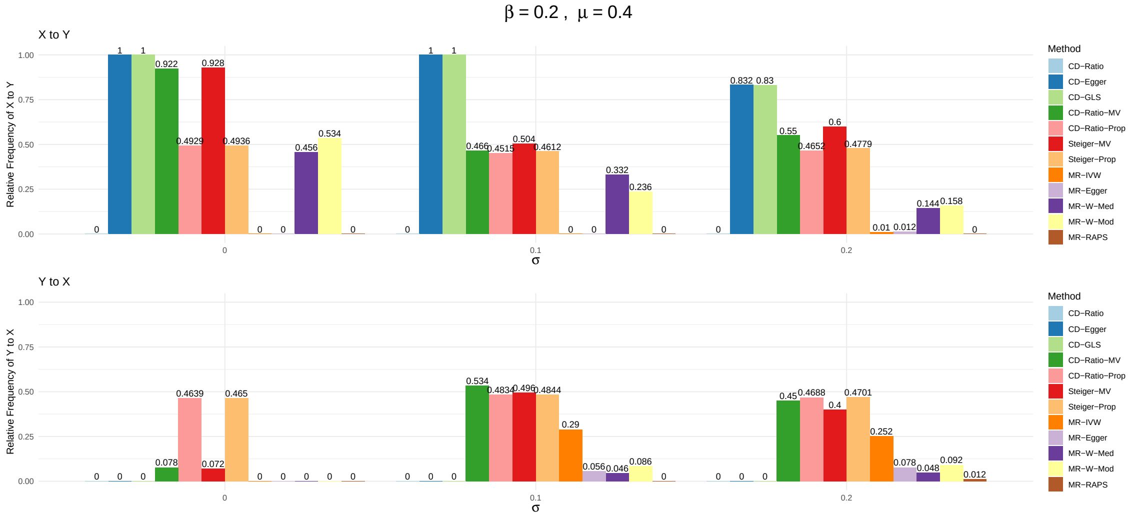


Figure S21: Relative frequencies of decisions for causal direction of all methods: $\beta_{YX} = 0.2$, $\mu_\alpha = 0.6$

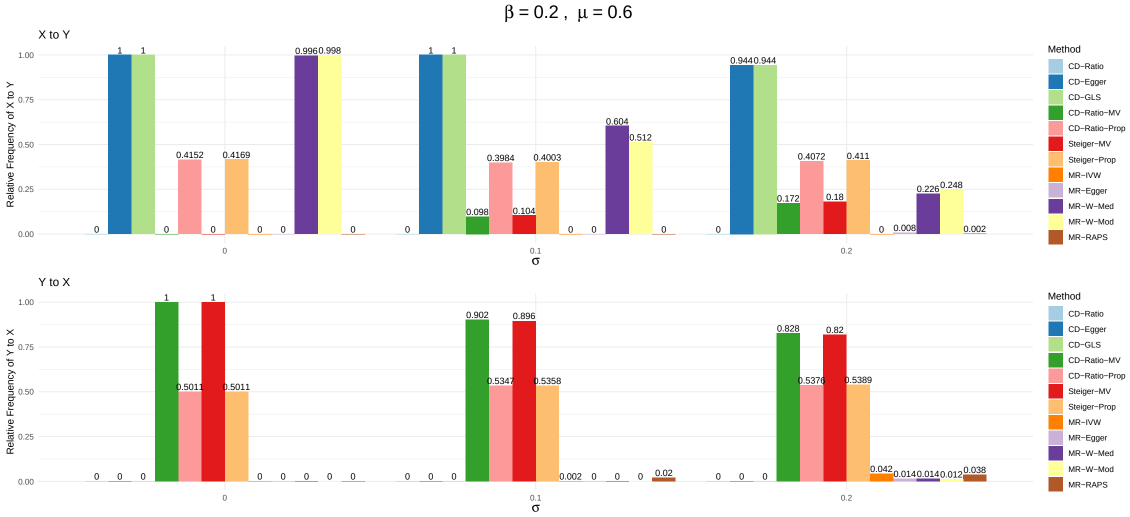

Figure S22: Relative frequencies of decisions for causal direction of all methods: $\beta_{YX} = 0.2$, $\mu_\alpha = 1$

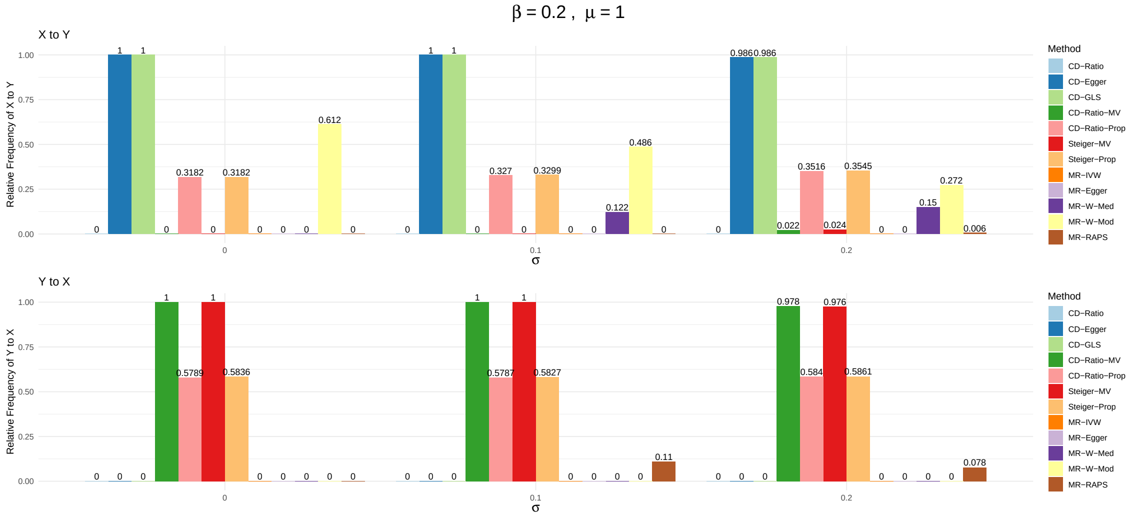


Figure S23: Relative frequencies of decisions for causal direction of all methods: $\beta_{YX} = -0.2$, $\mu_\alpha = 0$

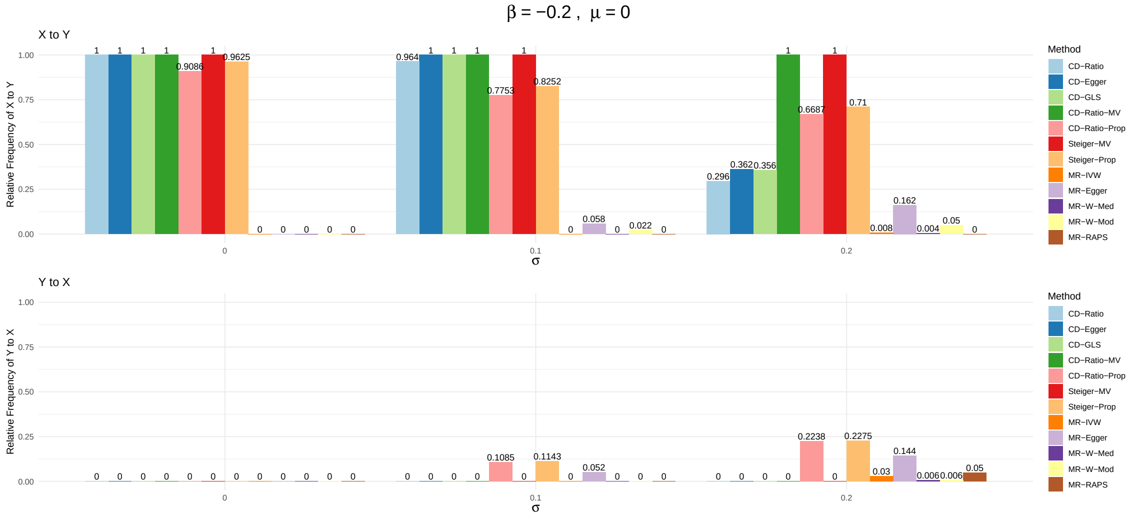

Figure S24: Relative frequencies of decisions for causal direction of all methods: $\beta_{YX} = -0.2$, $\mu_{\alpha} = 0.2$

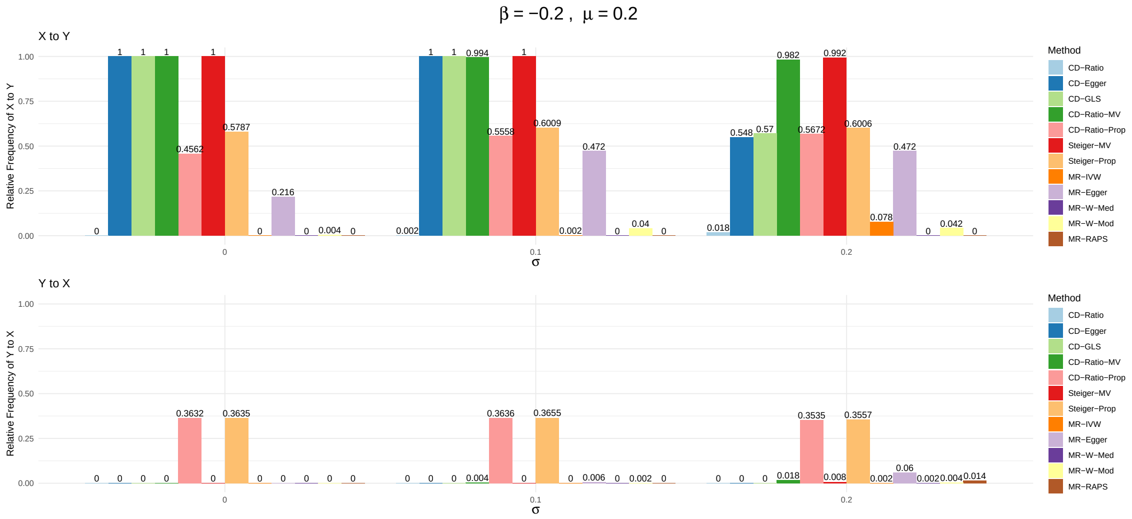


Figure S25: Relative frequencies of decisions for causal direction of all methods: $\beta_{YX} = -0.2$, $\mu_{\alpha} = 0.4$

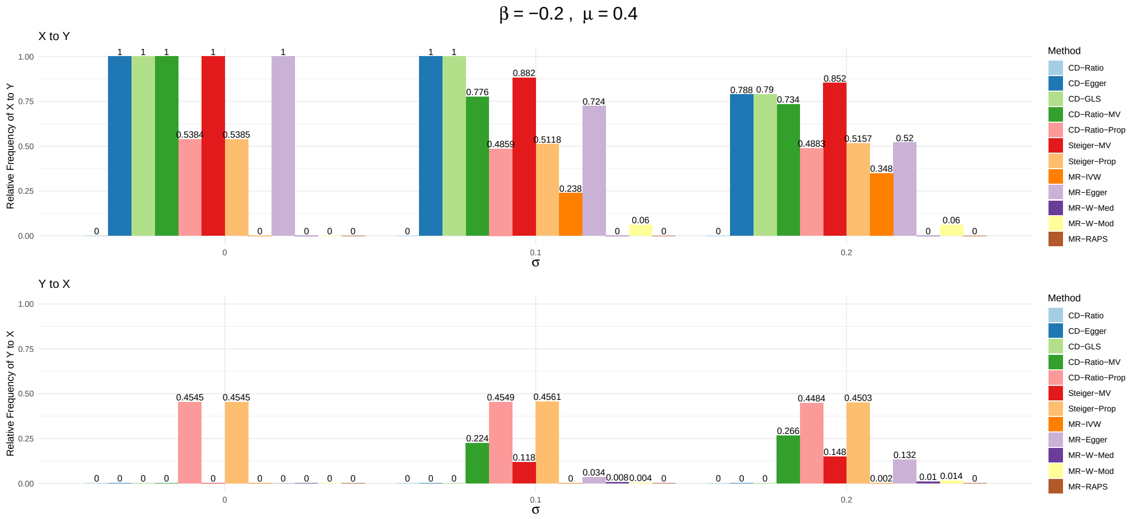

Figure S26: Relative frequencies of decisions for causal direction of all methods: $\beta_{YX} = -0.2$, $\mu_\alpha = 0.6$

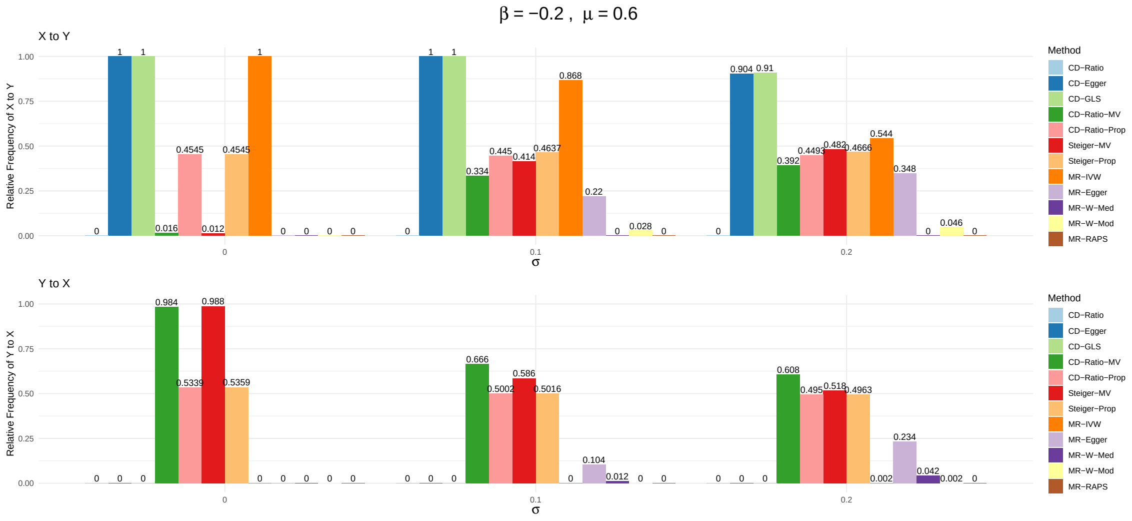


Figure S27: Relative frequencies of decisions for causal direction of all methods: $\beta_{YX} = -0.2$, $\mu_\alpha = 1$

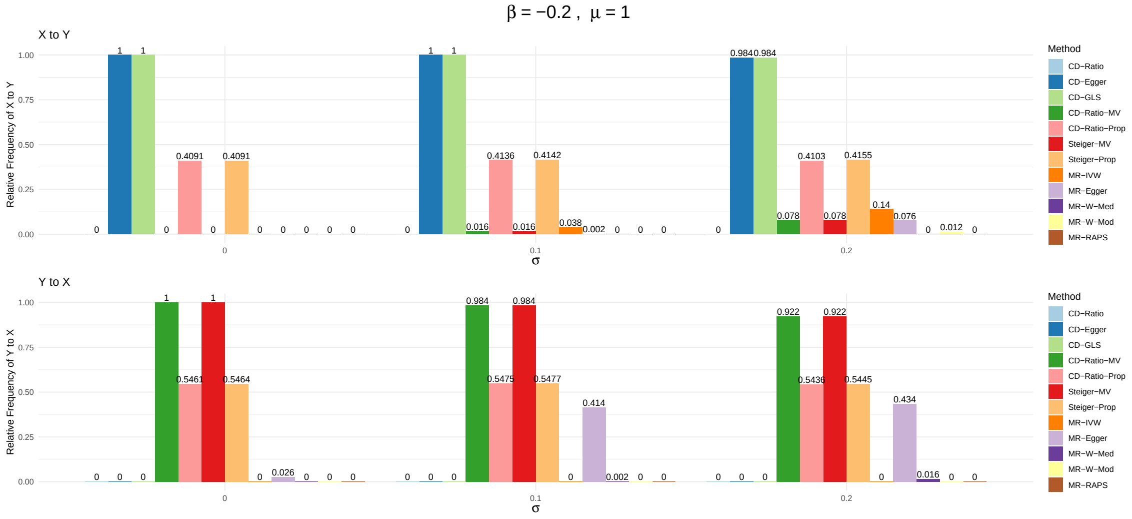


### 7 Simulations for Bi-directional Causal Effects

We set the sample size $n = 10000$, generated $g_1$ and $g_2$ independently with a minor allele frequency of 0.3. We generated the independent error terms $\varepsilon_1$ and $\varepsilon_2$ from a normal distribution with mean 0 and variance 3.2, and generated the confounder $U$ from a normal distribution with mean 0 and variance 0.8. We set $\gamma_1 = 1$, $\gamma_2 = 1$, $\beta_{XU} = 1$, and $\beta_{YU} = 1$. With different $\beta_1$ and $\beta_2$, we generated two independent samples of $X$ and $Y$ from the reduced form of the models (25) in the main text, using the first sample to get summary statistics for $X$ and the second for $Y$, and applied **CD-Ratio** to both directions, leading to out **bi-CD-Ratio**. For comparison, we also applied **MR-Steiger** and **MR-Wald-Ratio** to both directions. When considering the candidate direction of $X$ to $Y$, we used $g_1$ as the IV; for $Y$ to $X$, we used $g_2$ as the IV. For each setup we did simulations 1000 times, and drew conclusions on both directions based on the 95% confidence intervals. Table S38 shows the simulation results of estimating $K_1$, $K_2$ and their standard errors with bi-CD-Ratio, and Table S39 compares *bi-directional) CD-Ratio, MR-Steiger, and MR-Wald-Ratio for their relative frequencies of concluding with any causal directions.

Table S38: Simulation results for bi-CD-Ratio with different pairs of $\beta_1$ and $\beta_2$. For both directions we show the true value of $K$, the mean and standard deviation of estimates, and mean of standard errors.

| | $X \to Y$ | | | | $Y \to X$ | | | |
|---|---|---|---|---|---|---|---|---|
| $(\beta_1, \beta_2)$ | $K_1$ | Mean($\hat{K}_1$) | sd($\hat{K}_1$) | Mean($se(\hat{K}_1)$) | $K_2$ | Mean($\hat{K}_2$) | sd($\hat{K}_2$) | Mean($se(\hat{K}_2)$) |
| No causal effect | | | | | | | | |
| (0,0) | 0 | 0.001 | 0.032 | 0.033 | 0 | 0.001 | 0.032 | 0.032 |
| Unidirectional causal effect from $X$ to $Y$ | | | | | | | | |
| (-0.2,0) | -0.203 | -0.203 | 0.032 | 0.033 | 0 | 0.001 | 0.032 | 0.032 |
| (0.2,0) | 0.19 | 0.191 | 0.032 | 0.033 | 0 | 0.001 | 0.034 | 0.034 |
| Unidirectional causal effect from $Y$ to $X$ | | | | | | | | |
| (0,-0.2) | 0 | 0.001 | 0.031 | 0.032 | -0.203 | -0.203 | 0.033 | 0.033 |
| (0,0.2) | 0 | 0.001 | 0.034 | 0.034 | 0.19 | 0.191 | 0.032 | 0.033 |
| Bi-directional causal effect | | | | | | | | |
| (-0.2,-0.2) | -0.2 | -0.199 | 0.031 | 0.032 | -0.2 | -0.199 | 0.032 | 0.032 |
| (-0.2,0.2) | -0.214 | -0.214 | 0.034 | 0.035 | 0.187 | 0.188 | 0.032 | 0.032 |
| (0.2, -0.2) | 0.187 | 0.188 | 0.032 | 0.032 | -0.214 | -0.214 | 0.035 | 0.035 |
| (0.2, 0.2) | 0.2 | 0.202 | 0.034 | 0.035 | 0.2 | 0.201 | 0.034 | 0.035 |

Table S39: Comparison of (bi-directional) CD-Ratio, MR-Steiger and MR-Wald-Ratio for the relative frequencies of their conclusions on the causal directions.

| | | | | | | |
|---|---|---|---|---|---|---|
| No causal effect | | | | | | |
| | $X \to Y$ | | | $Y \to X$ | | |
| $(\beta_1, \beta_2)$ | CD-Ratio | MR-Steiger | MR-Wald-Ratio | CD-Ratio | MR-Steiger | MR-Wald-Ratio |
| (0,0) | 0.035 | 1 | 0.035 | 0.047 | 1 | 0.047 |
| Unidirectional causal effect from $X$ to $Y$ | | | | | | |
| | $X \to Y$ | | | $Y \to X$ | | |
| $(\beta_1, \beta_2)$ | CD-Ratio | MR-Steiger | MR-Wald-Ratio | CD-Ratio | MR-Steiger | MR-Wald-Ratio |
| (-0.2,0) | 1 | 1 | 1 | 0.047 | 1 | 0.047 |
| (0.2,0) | 1 | 1 | 1 | 0.047 | 1 | 0.047 |
| Unidirectional causal effect from $Y$ to $X$ | | | | | | |
| | $X \to Y$ | | | $Y \to X$ | | |
| $(\beta_1, \beta_2)$ | CD-Ratio | MR-Steiger | MR-Wald-Ratio | CD-Ratio | MR-Steiger | MR-Wald-Ratio |
| (0,-0.2) | 0.035 | 1 | 0.035 | 1 | 1 | 1 |
| (0,0.2) | 0.035 | 1 | 0.035 | 0.999 | 1 | 0.999 |
| Bi-directional causal effect | | | | | | |
| | $X \to Y$ | | | $Y \to X$ | | |
| $(\beta_1, \beta_2)$ | CD-Ratio | MR-Steiger | MR-Wald-Ratio | CD-Ratio | MR-Steiger | MR-Wald-Ratio |
| (-0.2,-0.2) | 1 | 1 | 1 | 1 | 1 | 1 |
| (-0.2,0.2) | 1 | 1 | 1 | 0.999 | 1 | 0.999 |
| (0.2,-0.2) | 1 | 1 | 1 | 1 | 1 | 1 |
| (0.2,0.2) | 1 | 1 | 1 | 0.999 | 1 | 0.999 |

From Table S38, we can see that for all situations, our proposed bi-CD-Ratio could estimate the true $K_1$ and $K_2$ pretty well, and the means of $\text{se}(\hat{K}_1)$ and $\text{se}(\hat{K}_1)$ were close to $\text{sd}(\hat{K}_1)$ and $\text{sd}(\hat{K}_2)$. From Table S39, when there was no causal relationship, both the bi-CD-Ratio and MR-Wald-Ratio could control the Type-I Errors around 0.05; when there was a causal direction, both methods could always detect it with a relative frequency of 1. MR-Steiger always concluded with the bi-directional causal effect due to the following reason: for $X$ to $Y$ we used $g_1$ as the valid instrument; $g_1$ always had a larger correlation with $X$ than that with $Y$ no matter whether $X$ had a causal effect on $Y$ or not; hence Steger's method would always conclude with a causal direction from $X$ to $Y$.; similarly, when considering $Y$ to $X$ with $g_2$ as the instrument, it would always conclude with a causal direction of $Y$ to $X$. and same for $Y$ to $X$. In contrast, based on correlation ratios, (bi-directional) CD-Ratio could determine the existence of a causal effect correctly by comparing the ratio with 0; on the other hand, MR-Steiger, based on differences of correlations, could not correctly determine the existence of a causal relationship under this situation.

### 8 Simulations for TWAS with Small Sample Sizes

We did a simulation study to investigate how the asymptotic theory for the sample correlations and their ratios performs with smaller sample sizes as typical with molecular endophenotypes as in TWAS. To mimick real TWAS, we used the fitted model from the ADNI gene expression and genotype data for 712 individuals as the data-generating model. For gene **PSPH** on chromosome 7, we identified 10 *SNP*s collectively explaining around 20% of the gene's expression variation in the fitted linear regression model: rs35515795, rs79278832, rs12154781, rs7791829, rs1057603, rs148444659, rs2242509, rs2908543, rs816417 and rs56875346. Figure S28 shows the correlation matrix of these 10 SNPs based on 712 individuals. From the fitted the linear regression model, we obtained the estimated regression coefficients $\hat{\beta}$'s for these 10 SNPs, and the estimated standard deviation of the error term $\hat{\sigma} = 0.803$. We used $\hat{\beta}$'s and $\hat{\sigma}$ in the corresponding linear model to generate realistic simulated data.

Figure S28: Correlation Structure of 10 SNPs in gene PSPH

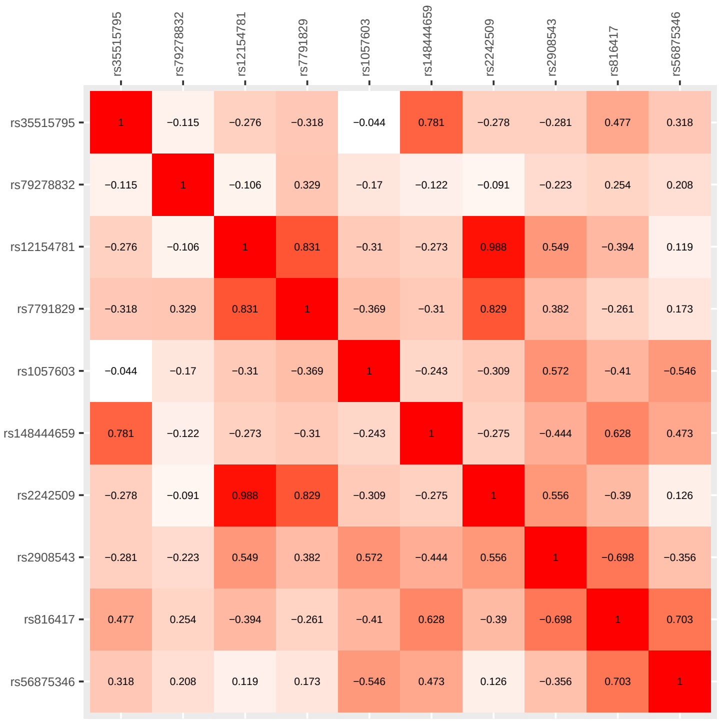


We generated the simulated data as the following:

$$
\begin{aligned}
&U \sim N(0, \hat{\sigma}^2/4) \\
&X = \sum_{j=1}^{10} \hat{\beta}_j \cdot SNP_j + U + \varepsilon_X, \varepsilon_X \sim N(0, \hat{\sigma}^2) \qquad (47) \\
&Y = \beta_{YX} \cdot X + U + \varepsilon_Y, \varepsilon_Y \sim N(0, \hat{\sigma}^2)
\end{aligned}
$$

Here $U$ is a confounder. We generated two independent samples. For the first sample, we randomly chose 200 out of the 712 ADNI individuals, and duplicated their genotypes $k \geq 1$ times to possibly increase the sample size. For the second sample, we randomly chose 150 from the remaining 512 individuals, and duplicated their genotypes $k$ times. With the first sample we obtains the sample correlations between $X$ and the 10 SNPs, and with the second sample the correlations between $Y$ and the 10 SNPs. Then we applied our CD-Ratio method to the two samples of these correlations. We applied CD-Ratio to each of 10 SNPs, and to combine their results; we also applied it to combine only those of the 9 SNPs without rs2242509, which had a high correlation $> 0.9$ with rs12154781. For each combination of $k = 1, 2, 3, 4, 5$ and $\beta_{YX} = -0.2, -0.1, 0, 0.2, 0.1$, we did simulations 1000 times. Each time we got a 95% CI for $K_{YX}$, then if it was completely inside [-1,0) or (0,1], we concluded that $X$ had a causal effect on $Y$; if it covered 0, we concluded that $X$ had no causal effect on $Y$. We also apply the MR Steiger method with each single SNP for comparison. Note that here, for simplicity, differing from our other numerical studies, we only considered either no or one causal direction from $X$ to $Y$. Table S40 shows the simulation results.

Table S40: Relative frequencies of concluding with $X$ having a causal effect on $Y$ from 1000 simulations for each setup of $(\beta_{YX}, k)$. For CD-Ratio, the results combining over 10 or 9 SNPs, and the maximum (max), mean and minimum (min) relative frequencies using each of the 10 single SNPs are shown. For MR-Steiger, the maximum, mean and minimum of relative frequencies using each of 10 SNPs are shown.

| $\beta_{YX} = 0$ | | | | | | | | |
|---|---|---|---|---|---|---|---|---|
| | CD-Ratio | | | | | MR-Steiger | | |
| $k$ | 10 SNPs | 9 SNPs | max | mean | min | max | mean | min |
| 1 | 0.021 | 0.022 | 0.004 | 4e-04 | 0 | 0.313 | 0.0417 | 0 |
| 2 | 0.033 | 0.029 | 0.015 | 0.0015 | 0 | 0.797 | 0.115 | 0 |
| 3 | 0.048 | 0.039 | 0.035 | 0.0043 | 0 | 0.958 | 0.1551 | 0 |
| 4 | 0.051 | 0.043 | 0.043 | 0.006 | 0 | 0.994 | 0.1818 | 0 |
| 5 | 0.05 | 0.048 | 0.047 | 0.006 | 0 | 0.999 | 0.1925 | 0 |
| $\beta_{YX} = 0.1$ | | | | | | | | |
| | CD-Ratio | | | | | MR-Steiger | | |
| $k$ | 10 SNPs | 9 SNPs | max | mean | min | max | mean | min |
| 1 | 0.024 | 0.024 | 0.004 | 4e-04 | 0 | 0.327 | 0.0441 | 0 |
| 2 | 0.066 | 0.075 | 0.023 | 0.0023 | 0 | 0.775 | 0.1131 | 0 |
| 3 | 0.112 | 0.127 | 0.056 | 0.0061 | 0 | 0.93 | 0.1511 | 0 |
| 4 | 0.133 | 0.12 | 0.085 | 0.0106 | 0 | 0.978 | 0.1801 | 0 |
| 5 | 0.134 | 0.134 | 0.112 | 0.0137 | 0 | 0.993 | 0.1892 | 0 |
| $\beta_{YX} = -0.1$ | | | | | | | | |
| | CD-Ratio | | | | | MR-Steiger | | |
| $k$ | 10 SNPs | 9 SNPs | max | mean | min | max | mean | min |
| 1 | 0.032 | 0.033 | 0.005 | 6e-04 | 0 | 0.306 | 0.0405 | 0 |
| 2 | 0.079 | 0.072 | 0.023 | 0.0024 | 0 | 0.756 | 0.1085 | 0 |
| 3 | 0.103 | 0.108 | 0.06 | 0.0067 | 0 | 0.938 | 0.155 | 0 |
| 4 | 0.153 | 0.161 | 0.086 | 0.0109 | 0 | 0.979 | 0.1755 | 0 |
| 5 | 0.153 | 0.151 | 0.108 | 0.0143 | 0 | 0.997 | 0.1905 | 0 |
| $\beta_{YX} = 0.2$ | | | | | | | | |
| | CD-Ratio | | | | | MR-Steiger | | |
| $k$ | 10 SNPs | 9 SNPs | max | mean | min | max | mean | min |
| 1 | 0.051 | 0.045 | 0.001 | 1e-04 | 0 | 0.292 | 0.0405 | 0 |
| 2 | 0.166 | 0.171 | 0.044 | 0.0046 | 0 | 0.693 | 0.1001 | 0 |
| 3 | 0.306 | 0.3 | 0.124 | 0.0133 | 0 | 0.843 | 0.1363 | 0 |
| 4 | 0.335 | 0.346 | 0.2 | 0.0233 | 0 | 0.926 | 0.1672 | 0 |
| 5 | 0.431 | 0.437 | 0.295 | 0.034 | 0 | 0.973 | 0.1819 | 0 |
| $\beta_{YX} = -0.2$ | | | | | | | | |
| | CD-Ratio | | | | | MR-Steiger | | |
| $k$ | 10 SNPs | 9 SNPs | max | mean | min | max | mean | min |
| 1 | 0.062 | 0.056 | 0.004 | 6e-04 | 0 | 0.264 | 0.0355 | 0 |
| 2 | 0.196 | 0.203 | 0.06 | 0.0063 | 0 | 0.646 | 0.0949 | 0 |
| 3 | 0.315 | 0.315 | 0.152 | 0.0159 | 0 | 0.844 | 0.1405 | 0 |
| 4 | 0.424 | 0.44 | 0.273 | 0.0311 | 0 | 0.926 | 0.1618 | 0 |
| 5 | 0.495 | 0.502 | 0.324 | 0.0392 | 0 | 0.976 | 0.1807 | 0 |

From Table S40 we can see that, when there was no causal effect, i.e. $\beta_{YX} = 0$, and when $k = 1$, i.e.

the sample sizes were small, CD-Ratio was a little conservative. As $k$ increased to 5, its empirical Type-I error rates would approach the nominal level 0.05. When there was a causal effect, i.e. $\beta_{YX} \neq 0$, as $k$ increased the power of CD-Ratio also increased, and the power with 10 or 9 SNPs was close and was higher than the maximum of using only single SNPs. For MR-Steiger, the maximum and mean relative frequencies from the 10 single SNPs gave some inflated Type-I error rates when $\beta_{YX} = 0$. Again it was because the SNPs had larger absolute correlations with $X$ than with $Y$, regardless of whether $X$ had a causal effect on $Y$ or not. Given its inflated type I error rates, the high power of MR-Steiger was not meaningful here. Note also here we considered only one possible causal direction of $X$ to $Y$.

### 9 Simulations to Study Selection Bias

Table S41: Simulation results for estimating $K_{YX}$ with no pleiotropy. The means, standard deviations (sd) and SEs (se) from each method are shown for each set-up based on 500 simulated datasets. $\mu_\alpha$ represents the mean of the pleiotropic/direct effects (with standard deviation $\sigma_\alpha = 0$). $N$ represents the number of top SNPs that are used.

| No pleiotropy | | | | | | | | | | | |
|---|---|---|---|---|---|---|---|---|---|---|---|
| | | | CD-Ratio | | | CD-Egger | | | CD-GLS | | |
| $\mu_\alpha$ | $K_{YX}$ | $N$ | Mean($\hat{K}$) | sd($\hat{K}$) | Mean$(se(\hat{K}))$ | Mean($\hat{K}$) | sd($\hat{K}$) | Mean$(se(\hat{K}))$ | Mean($\hat{K}$) | sd($\hat{K}$) | Mean$(se(\hat{K}))$ |
| 0 | 0 | 10 | -0.0001 | 0.0069 | 0.0052 | 0 | 0.007 | 0.0063 | 0 | 0.007 | 0.0063 |
| | | 15 | 0.0001 | 0.0048 | 0.0042 | 0.0001 | 0.0048 | 0.0047 | 0.0001 | 0.0048 | 0.0047 |
| | | 22 | 0.0001 | 0.0034 | 0.0035 | 0.0001 | 0.0034 | 0.0036 | 0.0001 | 0.0034 | 0.0036 |
| 0 | 0.2257 | 10 | 0.2261 | 0.0038 | 0.0041 | 0.226 | 0.0039 | 0.0042 | 0.226 | 0.0039 | 0.0043 |
| | | 15 | 0.2261 | 0.0036 | 0.0038 | 0.2261 | 0.0036 | 0.0039 | 0.2261 | 0.0036 | 0.0039 |
| | | 22 | 0.2256 | 0.0033 | 0.0036 | 0.2256 | 0.0033 | 0.0036 | 0.2256 | 0.0033 | 0.0036 |
| 0 | -0.2344 | 10 | -0.2346 | 0.0038 | 0.0042 | -0.2346 | 0.0039 | 0.0042 | -0.2345 | 0.0039 | 0.0043 |
| | | 15 | -0.2346 | 0.0035 | 0.0038 | -0.2346 | 0.0035 | 0.0039 | -0.2346 | 0.0035 | 0.0039 |
| | | 22 | -0.2341 | 0.0032 | 0.0036 | -0.2341 | 0.0032 | 0.0036 | -0.2341 | 0.0032 | 0.0036 |

Table S42: Simulation results for estimating $K_{YX}$ with balanced pleiotropy. The means, standard deviations (sd) and SEs (se) from each method are shown for each set-up based on 500 simulated datasets. $\mu_\alpha$ represents the mean of the pleiotropic/direct effects (with standard deviation $\sigma_\alpha = 0.1$). $N$ represents the number of top SNPs that are used.

| Balanced pleiotropy | | | | | | | | | | | |
|---|---|---|---|---|---|---|---|---|---|---|---|
| | | | CD-Ratio | | | CD-Egger | | | CD-GLS | | |
| $\mu_\alpha$ | $K_{YX}$ | $N$ | Mean($\hat{K}$) | sd($\hat{K}$) | Mean$(se(\hat{K}))$ | Mean($\hat{K}$) | sd($\hat{K}$) | Mean$(se(\hat{K}))$ | Mean($\hat{K}$) | sd($\hat{K}$) | Mean$(se(\hat{K}))$ |
| 0 | 0 | 10 | 0.0007 | 0.1023 | 0.0053 | -0.0057 | 0.0801 | 0.0741 | -0.0056 | 0.0803 | 0.0741 |
| | | 15 | 0.0013 | 0.0767 | 0.0044 | -0.0017 | 0.0579 | 0.0547 | -0.0017 | 0.0587 | 0.0551 |
| | | 22 | 0.0013 | 0.0563 | 0.0037 | -0.0007 | 0.0431 | 0.041 | -0.0005 | 0.0436 | 0.0419 |
| 0 | 0.2257 | 10 | 0.2801 | 0.0615 | 0.0046 | 0.2531 | 0.0586 | 0.0591 | 0.2535 | 0.0587 | 0.059 |
| | | 15 | 0.2479 | 0.0579 | 0.0041 | 0.2339 | 0.0508 | 0.0503 | 0.2341 | 0.051 | 0.0504 |
| | | 22 | 0.2181 | 0.0531 | 0.0037 | 0.2201 | 0.0397 | 0.0393 | 0.2203 | 0.0402 | 0.0401 |
| 0 | -0.2344 | 10 | -0.2871 | 0.0636 | 0.0047 | -0.265 | 0.0615 | 0.0628 | -0.2652 | 0.0615 | 0.0627 |
| | | 15 | -0.2545 | 0.0595 | 0.0041 | -0.2454 | 0.0524 | 0.0527 | -0.2455 | 0.0525 | 0.0527 |
| | | 22 | -0.2233 | 0.0545 | 0.0037 | -0.2296 | 0.0409 | 0.0408 | -0.2295 | 0.0414 | 0.0416 |

Table S43: Simulation results for estimating $K_{YX}$ with directional pleiotropy. The means, standard deviations (sd) and SEs (se) from each method are shown for each set-up based on 500 simulated datasets. $\mu_\alpha$ represents the mean of the pleiotropic/direct effects (with standard deviation $\sigma_\alpha = 0.1$). $N$ represents the number of top SNPs that are used.

| Directional pleiotropy | | | | | | | | | | | |
|---|---|---|---|---|---|---|---|---|---|---|---|
| | | | CD-Ratio | | | CD-Egger | | | CD-GLS | | |
| $\mu_\alpha$ | $K_{YX}$ | $N$ | Mean($\hat{K}$) | sd($\hat{K}$) | Mean$(se(\hat{K}))$ | Mean($\hat{K}$) | sd($\hat{K}$) | Mean$(se(\hat{K}))$ | Mean($\hat{K}$) | sd($\hat{K}$) | Mean$(se(\hat{K}))$ |
| 0.2 | 0 | 10 | -0.0908 | 0.1345 | 0.0053 | 0.02 | 0.0529 | 0.0618 | 0.02 | 0.0529 | 0.0617 |
| | | 15 | -0.0347 | 0.0843 | 0.0044 | -0.0017 | 0.0428 | 0.0508 | -0.0017 | 0.0428 | 0.0507 |
| | | 22 | -0.0112 | 0.0538 | 0.0039 | -0.0005 | 0.0386 | 0.0368 | -0.0004 | 0.0392 | 0.0375 |
| 0.4 | 0 | 10 | -0.1389 | 0.1189 | 0.0051 | 0.0297 | 0.041 | 0.086 | 0.0297 | 0.041 | 0.0859 |
| | | 15 | -0.0857 | 0.0824 | 0.0045 | -0.0029 | 0.0381 | 0.0689 | -0.0029 | 0.0381 | 0.0687 |
| | | 22 | -0.0183 | 0.0448 | 0.0041 | -0.0004 | 0.0306 | 0.0291 | -0.0003 | 0.0311 | 0.0297 |
| 0.6 | 0 | 10 | -0.2007 | 0.1215 | 0.0052 | 0.0419 | 0.0363 | 0.0991 | 0.0418 | 0.0363 | 0.0989 |
| | | 15 | -0.1052 | 0.0673 | 0.0045 | -0.009 | 0.032 | 0.0783 | -0.009 | 0.032 | 0.0781 |
| | | 22 | -0.0317 | 0.0357 | 0.0042 | -0.0003 | 0.024 | 0.0228 | -0.0002 | 0.0244 | 0.0233 |
| 1 | 0 | 10 | -0.2569 | 0.1313 | 0.0053 | 0.0553 | 0.0306 | 0.1098 | 0.0551 | 0.0306 | 0.1095 |
| | | 15 | -0.1162 | 0.0461 | 0.0045 | -0.0106 | 0.0234 | 0.0854 | -0.0108 | 0.0234 | 0.085 |
| | | 22 | -0.0506 | 0.0241 | 0.0043 | -0.0002 | 0.0161 | 0.0153 | -0.0002 | 0.0164 | 0.0156 |
| 0.2 | 0.2027 | 10 | 0.3346 | 0.1275 | 0.0055 | 0.0563 | 0.0751 | 0.0692 | 0.0563 | 0.075 | 0.0691 |
| | | 15 | 0.2443 | 0.0844 | 0.0045 | 0.1211 | 0.0499 | 0.0519 | 0.1212 | 0.0499 | 0.0518 |
| | | 22 | 0.1898 | 0.0514 | 0.0039 | 0.1987 | 0.0369 | 0.0355 | 0.1989 | 0.0375 | 0.0362 |
| 0.4 | 0.1614 | 10 | 0.1881 | 0.1075 | 0.005 | 0.0984 | 0.0587 | 0.0951 | 0.0983 | 0.0587 | 0.095 |
| | | 15 | 0.1739 | 0.0857 | 0.0045 | 0.0996 | 0.0412 | 0.0698 | 0.0996 | 0.0412 | 0.0696 |
| | | 22 | 0.1438 | 0.0448 | 0.0041 | 0.1593 | 0.0301 | 0.0285 | 0.1595 | 0.0307 | 0.029 |
| 0.6 | 0.1272 | 10 | 0.0887 | 0.1052 | 0.005 | 0.1067 | 0.0511 | 0.1023 | 0.1066 | 0.0512 | 0.102 |
| | | 15 | 0.0851 | 0.0812 | 0.0044 | 0.0939 | 0.0342 | 0.0788 | 0.0938 | 0.0342 | 0.0786 |
| | | 22 | 0.0944 | 0.0362 | 0.0042 | 0.1261 | 0.024 | 0.0225 | 0.1263 | 0.0244 | 0.023 |
| 1 | 0.0856 | 10 | 0.0104 | 0.1078 | 0.0051 | 0.1021 | 0.0307 | 0.1091 | 0.1019 | 0.0307 | 0.1089 |
| | | 15 | -0.0067 | 0.0553 | 0.0044 | 0.0636 | 0.0235 | 0.0869 | 0.0634 | 0.0235 | 0.0865 |
| | | 22 | 0.0313 | 0.0244 | 0.0043 | 0.0851 | 0.0162 | 0.0152 | 0.0852 | 0.0165 | 0.0155 |
| 0.2 | -0.2093 | 10 | -0.4414 | 0.0938 | 0.0052 | -0.1217 | 0.0642 | 0.0681 | -0.1217 | 0.0642 | 0.0681 |
| | | 15 | -0.3054 | 0.0714 | 0.0046 | -0.1886 | 0.0519 | 0.053 | -0.1885 | 0.0519 | 0.0528 |
| | | 22 | -0.2172 | 0.0517 | 0.0039 | -0.2061 | 0.037 | 0.0366 | -0.206 | 0.0373 | 0.0373 |
| 0.4 | -0.1649 | 10 | -0.4235 | 0.0964 | 0.0053 | -0.0929 | 0.046 | 0.0866 | -0.093 | 0.046 | 0.0864 |
| | | 15 | -0.3038 | 0.0687 | 0.0048 | -0.1428 | 0.04 | 0.0679 | -0.1429 | 0.0399 | 0.0676 |
| | | 22 | -0.1796 | 0.0435 | 0.0041 | -0.1636 | 0.0298 | 0.029 | -0.1635 | 0.03 | 0.0296 |
| 0.6 | -0.129 | 10 | -0.4022 | 0.0538 | 0.0054 | -0.0712 | 0.0367 | 0.1002 | -0.0714 | 0.0368 | 0.0999 |
| | | 15 | -0.2979 | 0.0778 | 0.0048 | -0.1129 | 0.032 | 0.0771 | -0.113 | 0.032 | 0.0768 |
| | | 22 | -0.1552 | 0.0351 | 0.0042 | -0.1285 | 0.0236 | 0.0228 | -0.1285 | 0.0238 | 0.0233 |
| 1 | -0.0862 | 10 | -0.4157 | 0.055 | 0.0055 | -0.0247 | 0.0261 | 0.1112 | -0.0248 | 0.0261 | 0.1108 |
| | | 15 | -0.2396 | 0.0762 | 0.0047 | -0.0882 | 0.0252 | 0.0843 | -0.0883 | 0.0252 | 0.0839 |
| | | 22 | -0.1301 | 0.0238 | 0.0043 | -0.0862 | 0.0159 | 0.0153 | -0.0862 | 0.0161 | 0.0156 |

Figure S29: Relative frequencies of decisions for causal direction of proposed methods when there is no causal relationship between $X$ and $Y$ ($K_{YX} = 0$) and no pleiotropic effects ($\mu_\alpha = 0, \sigma_\alpha = 0$) for different $N = 10, 15, 22$.

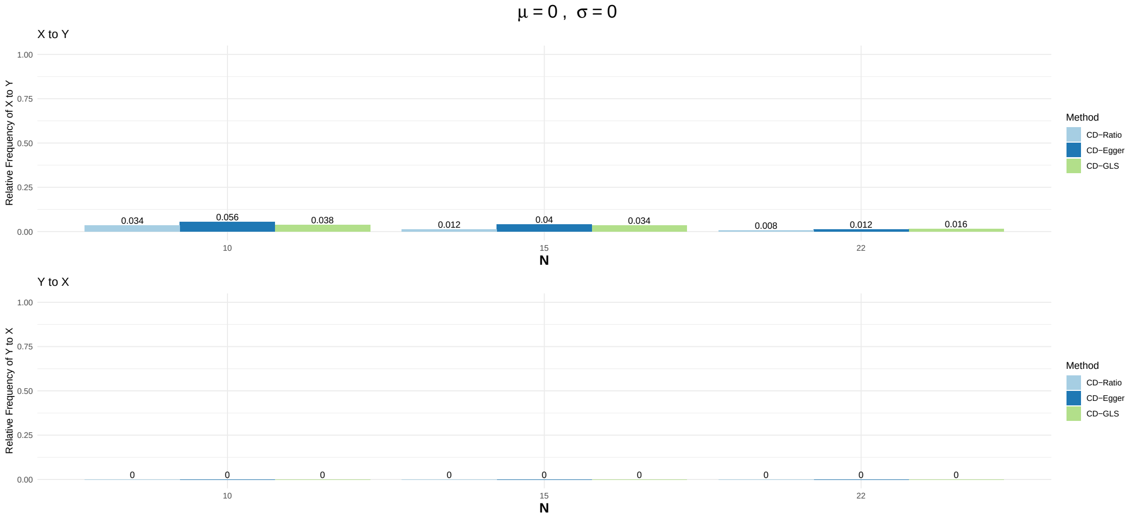


Figure S30: Relative frequencies of decisions for causal direction of proposed methods when there is no causal relationship between $X$ and $Y$ ($K_{YX} = 0$) and balanced pleiotropic effects ($\mu_\alpha = 0, \sigma_\alpha = 0.1$) for different $N = 10, 15, 22$.

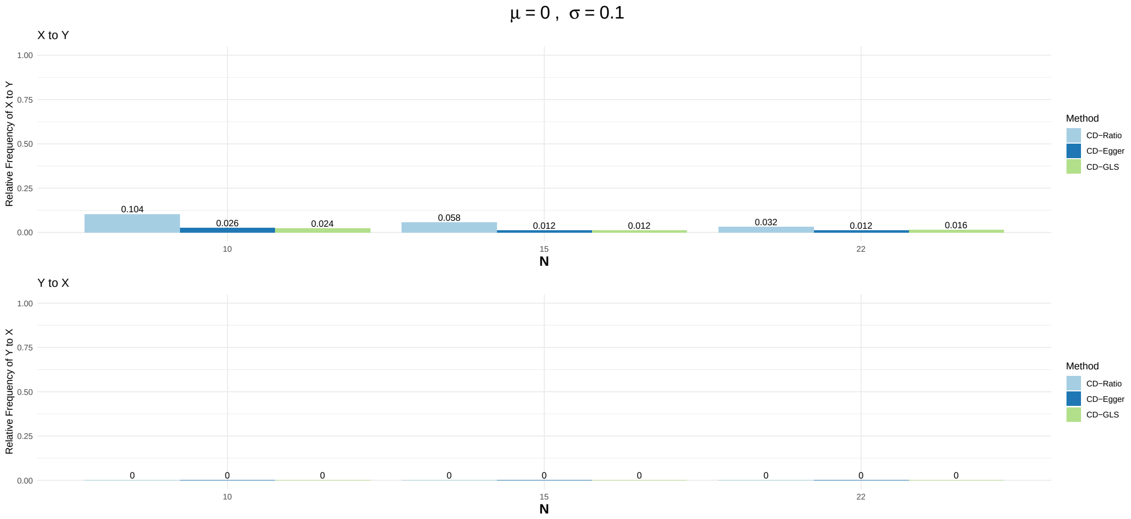

Figure S31: Relative frequencies of decisions for causal direction of proposed methods when there is no causal relationship between $X$ and $Y$ ($K_{YX} = 0$) and directional pleiotropic effects ($\mu_\alpha = 0.2, \sigma_\alpha = 0.1$) for different $N = 10, 15, 22$.

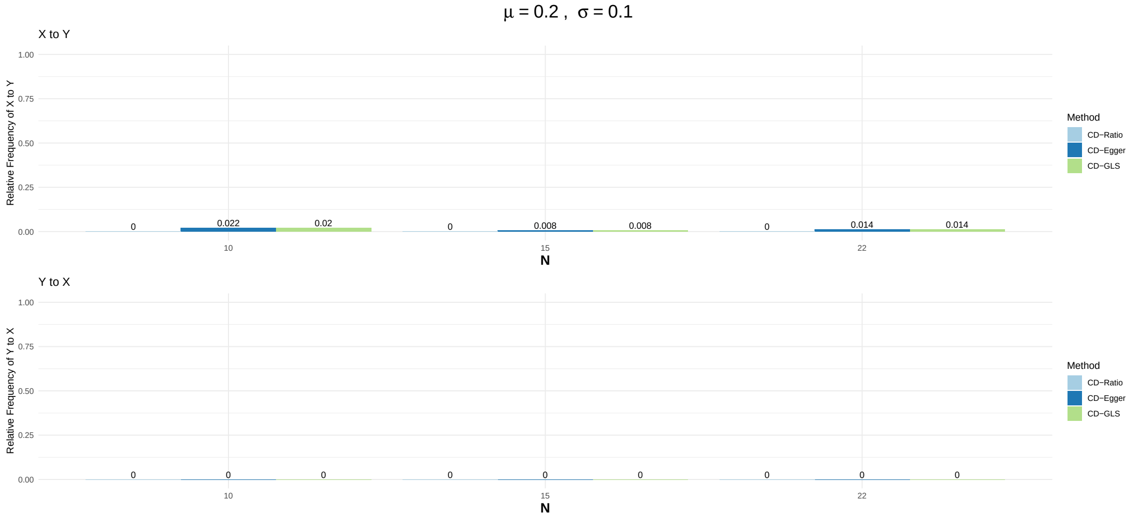


Figure S32: Relative frequencies of decisions for causal direction of proposed methods when there is no causal relationship between $X$ and $Y$ ($K_{YX} = 0$) and directional pleiotropic effects ($\mu_\alpha = 0.4, \sigma_\alpha = 0.1$) for different $N = 10, 15, 22$.

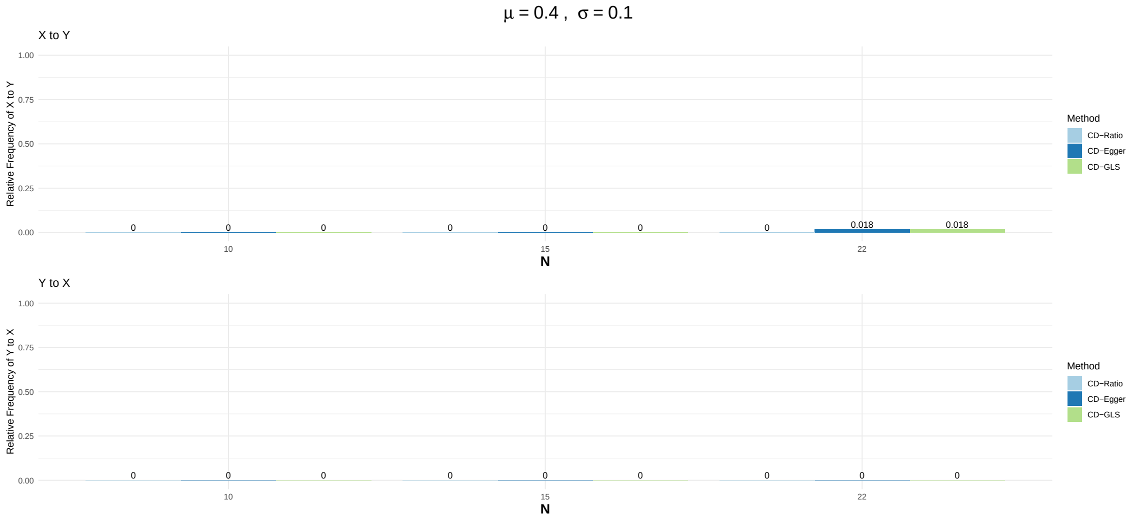

Figure S33: Relative frequencies of decisions for causal direction of proposed methods when there is no causal relationship between $X$ and $Y$ ($K_{YX} = 0$) and directional pleiotropic effects ($\mu_\alpha = 0.6, \sigma_\alpha = 0.1$) for different $N = 10, 15, 22$.

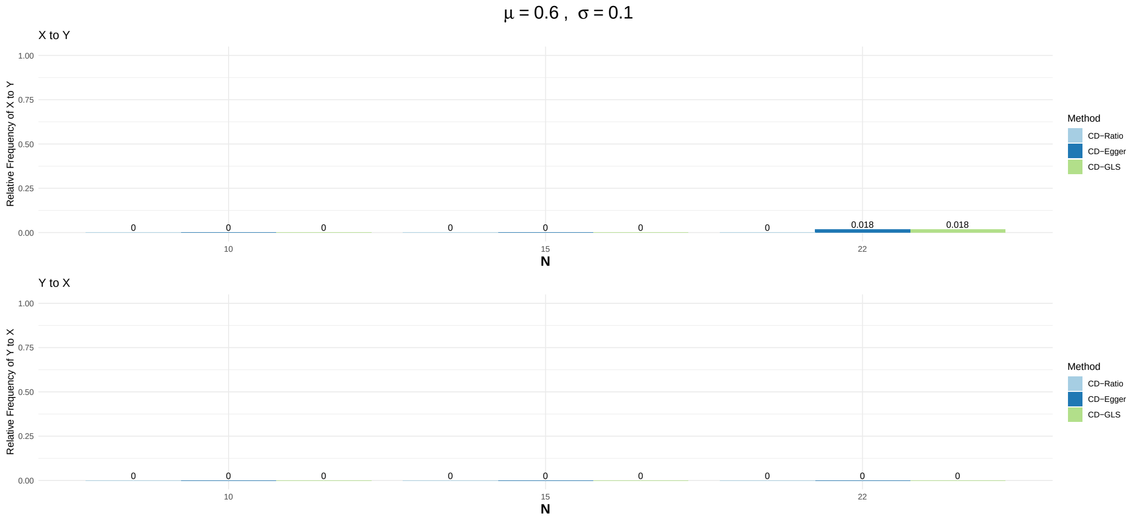


Figure S34: Relative frequencies of decisions for causal direction of proposed methods when there is no causal relationship between $X$ and $Y$ ($K_{YX} = 0$) and directional pleiotropic effects ($\mu_\alpha = 1, \sigma_\alpha = 0.1$) for different $N = 10, 15, 22$.

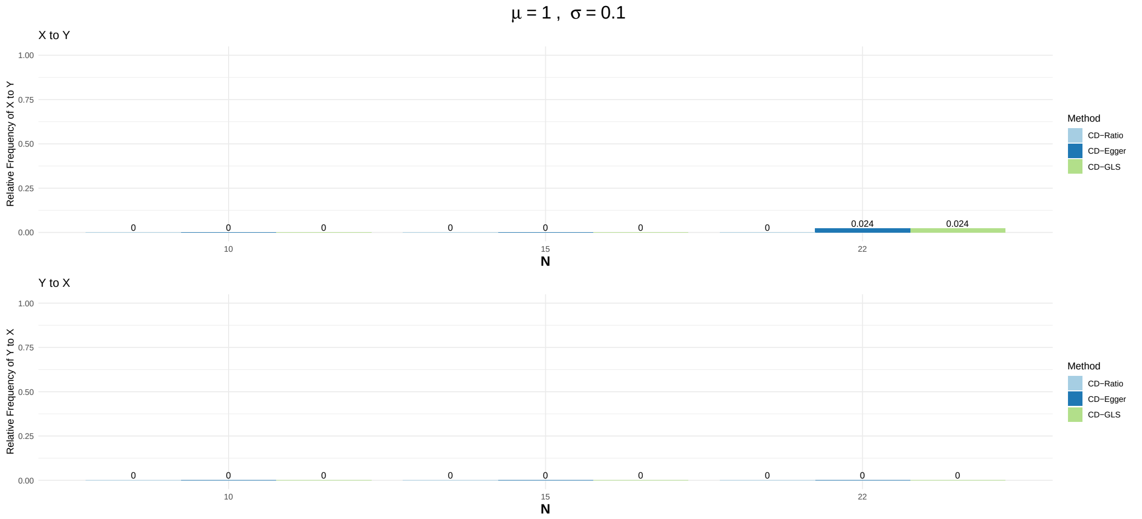


#### 10 Fisher's Transformation to Absolute Correlations

In the left panel of Figure S35, we draw the density function of normal distribution with mean 1 and standard error 1, i.e. $N(1,1)$, with blue color, and density function of its absolute value with red color. The right panel is for normal distribution with mean 1 and standard error 0.5, i.e. $N(1,0.5)$. We can see when standard error is 1, the original density function is different from the density function of absolute value. As standard error decreases to 0.5, the original density function is close to the density function of absolute value.

Figure S35: Comparing original density function and density function of absolute value of normal distributions, left panel is for $N(1,1)$ and right panel is for $N(1,0.5)$.

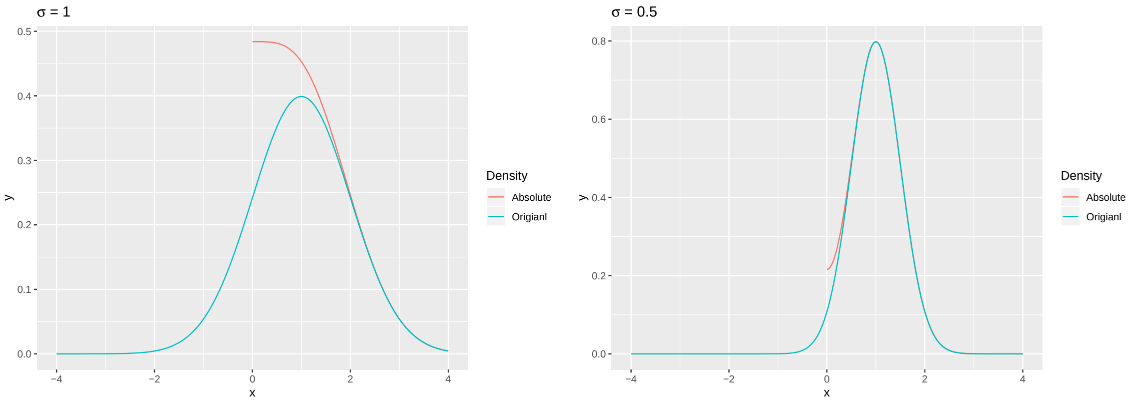
